# A network of bZIP transcription factors and inhibitors enforces differentially proliferative and cytotoxic states in T cells

**DOI:** 10.64898/2026.10.28.762687

**Authors:** Jeshua L. Malroy, Rangapriya Sundararajan, Maria Stegantseva, Sean Hyslop, Pablo García-Díaz, Ashley Zhang, Sarah Zhang, Lechuan Li, Alex Chen, Ying Wang, Leonid S. Metelitsa, Vicky Yao, William H. Hudson, Glenna Wink Foight

## Abstract

The bZIP family of transcription factors (TFs) plays a central role in regulating T-cell exhaustion. Their complex dimeric interactions obscure the roles of individual members, which genetic knockout or overexpression alone poorly resolve. Here, we use overexpression of full-length bZIP family TFs alongside genetically encoded, protein-based inhibitors to probe the effect of increased or decreased bZIP TF activity on CAR T-cell expansion under exhausting conditions. Enrichment of inhibitors of pro-exhaustion TFs (CREM, NFIL3) illustrated the utility of this approach. The screen enriched full-length TFs of established anti-exhaustion relevance and identified underappreciated bZIP TFs such as BATF2. Detailed comparison of several bZIP TFs and inhibitors revealed distinct effects on proliferation, persistence, and cytotoxicity. We investigated combinations of pro-cytotoxic JUN, pro-proliferative BATF2, and a novel inhibitor (AEL-CREM) of pro-dysfunction CREM. Distinct pairs were more advantageous for maintenance of cytotoxicity or proliferation, revealing relationships between BATF2, JUN-, and CREB1-family bZIPs. Using a combination of epistasis, transcriptome, proteome, and AP-MS interactome analyses of BATF2 and AEL-CREM, we discovered functional overlap in their inhibition of pro-dysfunction CREB1-family activity, potentially driven by unexpected binding of CREB1-family members by overexpressed BATF2. Their function diverged in BATF2’s early promotion of pro-proliferation AP-1 activity and later physical and functional interactions with BACH2, contributing to persistence. These findings were mirrored in murine chronic viral infection and syngeneic tumor models, where BATF2 or AEL-CREM overexpression were notably pro-proliferative and anti-exhaustion. Together, these results illuminate new connections and functional roles enabled by bZIP family perturbations, with implications for modulating T-cell state.

**One sentence summary:** Overexpression and protein-based inhibition of bZIP-family transcription factors in T cells reveal a network of BATF-, JUN-, CREB1-, and BACH2-family bZIPs regulating T cell exhaustion.

## INTRODUCTION

T-cell dysfunction, a hypofunctional T-cell state, is characterized by reduced proliferation, expression of inhibitory receptors, and reduced cytotoxicity, which are barriers to T-cell immunotherapy approaches to treating cancer, especially solid tumors. In the context of cancer, T cells are driven to a dysfunctional, exhausted state via several mechanisms, including chronic antigen stimulation, metabolic deficiency, catecholamine and PGE_2_ signaling, among other tumor microenvironment (TME) triggers of dysfunction (*1*, *2*). While multiple types of genetic modifications have been investigated to mitigate T-cell dysfunction, transcription factor (TF) perturbations are particularly promising due to the inherent ability of TFs to induce widespread transcriptional alterations necessary to rewire a dysfunctional transcriptional state (*3*). TFs such as JUN (*4*) or FOXO1 (*5*, *6*) can promote functional effector or memory states, while other TFs promote dysfunction, such as NR4A1/2/3 (*7*, *8*), TOX/TOX2 (*9*), CREM (*10*, *11*), NFAT5 (*12*), PRDM1 (*8*), and XBP1 (*13*, *14*). Several members of the basic leucine zipper (bZIP) family of transcription factors are known to regulate aspects of T-cell function, including effector (JUN, BATF; (*4*, *15*)) and memory (BACH2, BATF3; (*16–18*)) T-cell states, as well as responses to chronic stimulation and stress in TME conditions (XBP1, CHOP (DDIT3), CREM, CREB1, NFIL3, and ATF4 (*10*, *11*, *13*, *14*, *19–23*)). While focused studies on the role of AP-1-family bZIPs (JUN-, FOS-, BATF-, and some ATF-family members) have been performed (*4*, *15*, *18*), there are many other bZIPs with uncharacterized roles in T cells. Furthermore, there are conflicting data concerning the effects of BATF and ATF4 knockout/overexpression (*15*, *23–25*) on T-cell fitness depending on the experimental system, which indicates that bZIPs can act in a context-dependent manner to impede or promote T-cell function. Because the 53 human bZIPs form homo- and/or heterodimers with distinct binding partners which in turn regulate binding to different DNA sites (*26–28*), the bZIP protein-protein interaction network and downstream transcriptional outputs are likely to be highly sensitive to changes in expression level from overexpression or knockout. Due to these intricacies of bZIP combinatorial complexity, uncertainty remains regarding which bZIPs are most central to regulating T-cell dysfunction or promoting functional states, as well as their binding partners and molecular mechanisms.

One widely used method to explore the genetic dependencies of T cell function is genetic screening, with overexpression, knockout, and chimeric TF screens reported in the literature (*22*, *29–31*). While these screens have yielded a wealth of knowledge, none have directly compared the effects of genetic overexpression and knockdown in the same screen or assessed the roles of protein (leucine zipper [LZ]) versus DNA binding domain (basic region) on phenotype. To probe the complexity of bZIP function in T cells, we developed an approach to directly compare the effect of overexpressed full-length bZIP TFs, bZIP subdomains, and genetically encoded, protein-based inhibitors on T-cell proliferation/persistence in a single screen. We leveraged prior work in the bZIP protein engineering field (*32–38*) to systematically design bZIP inhibitors against all 53 human bZIP TFs with the potential to improve T-cell exhaustion resistance and inform our mechanistic understanding of protein and DNA interactions underlying bZIP phenotypic effects.

Through screening this bZIP variant library in CAR T cells under exhausting conditions, we identified multiple full-length bZIP TFs and inhibitor variants that impact T-cell proliferation or persistence. Inhibitors formed by addition of acidic extensions to leucine zippers were the most effective inhibitor variants for altering CAR-T cell proliferation, suggesting that blocking the basic DNA binding domain is required for a strong effect. Upon phenotypic, transcriptomic, proteomic and interactome profiling of screen hits in human CAR T cells in vitro, murine CAR T cells in a solid tumor model, and murine T cells in a chronic viral infection model, we uncovered roles of overexpressed BATF2 as a pro-persistence bZIP that slows exhaustion in part by forming unexpected interactions with BACH2 and CREB1-family members (CREB1, CREM, ATF1). In agreement that inhibition of CREB1-family members enhances proliferation and exhaustion resistance, AEL-CREM, an inhibitor of this family, also showed benefit in vitro and in a murine chronic infection model. Single and paired modifications of T cells with BATF2, JUN, and AEL-CREM demonstrated disparate effects on proliferation, cytotoxicity, and exhaustion suppression, which we trace to interplay between these bZIP subfamilies. Collectively, our results highlight the potential of bZIP engineering as a promising avenue to mitigate T-cell dysfunction and the importance of evaluating effects mediated by altered protein-protein interactions.

## RESULTS

### Diverse bZIP subfamilies and inhibitors impact persistence

To interrogate the role of bZIPs in T-cell dysfunction, we generated a pooled library consisting of full-length, subdomain (bZIP or leucine zipper [LZ] domain), bZIP inhibitors, as well as select non-bZIP TFs with relevance to T cells (Fig.1A, Table S1). The bZIP inhibitors utilized anti-bZIPs or acidic helical extensions. Anti-bZIPs are computationally designed LZs with specificity for binding bZIP subfamilies (*32–34*). Acidic extensions (AE) are engineered peptide sequences that can be fused to the N-terminus of a native LZ or designed anti-bZIP to dimerize with the basic DNA binding domain of a target bZIP, thus preventing DNA binding and serving as strong inhibitors (*35–38*). Two versions of the AE (AEN-or AEL-) were used, depending on the sequence of the putative target(s) of the inhibitor (see Materials and Methods, Table S1). Inhibitors were expected to alter the stoichiometry of protein-protein interactions, with the added affinity of the AEs expected to prevent DNA interactions. The variants were assembled in a gammaretroviral backbone with a truncated NGFR (tNGFR) transduction marker and multiple (mean = 41) unique DNA barcodes associated with each variant to enable tracking of the frequency of library members in the pooled screen (Fig. S1A-D). The transduction marker tNGFR-only was used as a negative control, and a constitutively active IL-7 cytokine receptor (C7R), clinically validated to improve GD2 CAR T-cell persistence (*39–41*), was used as a positive control.

**Figure 1.**
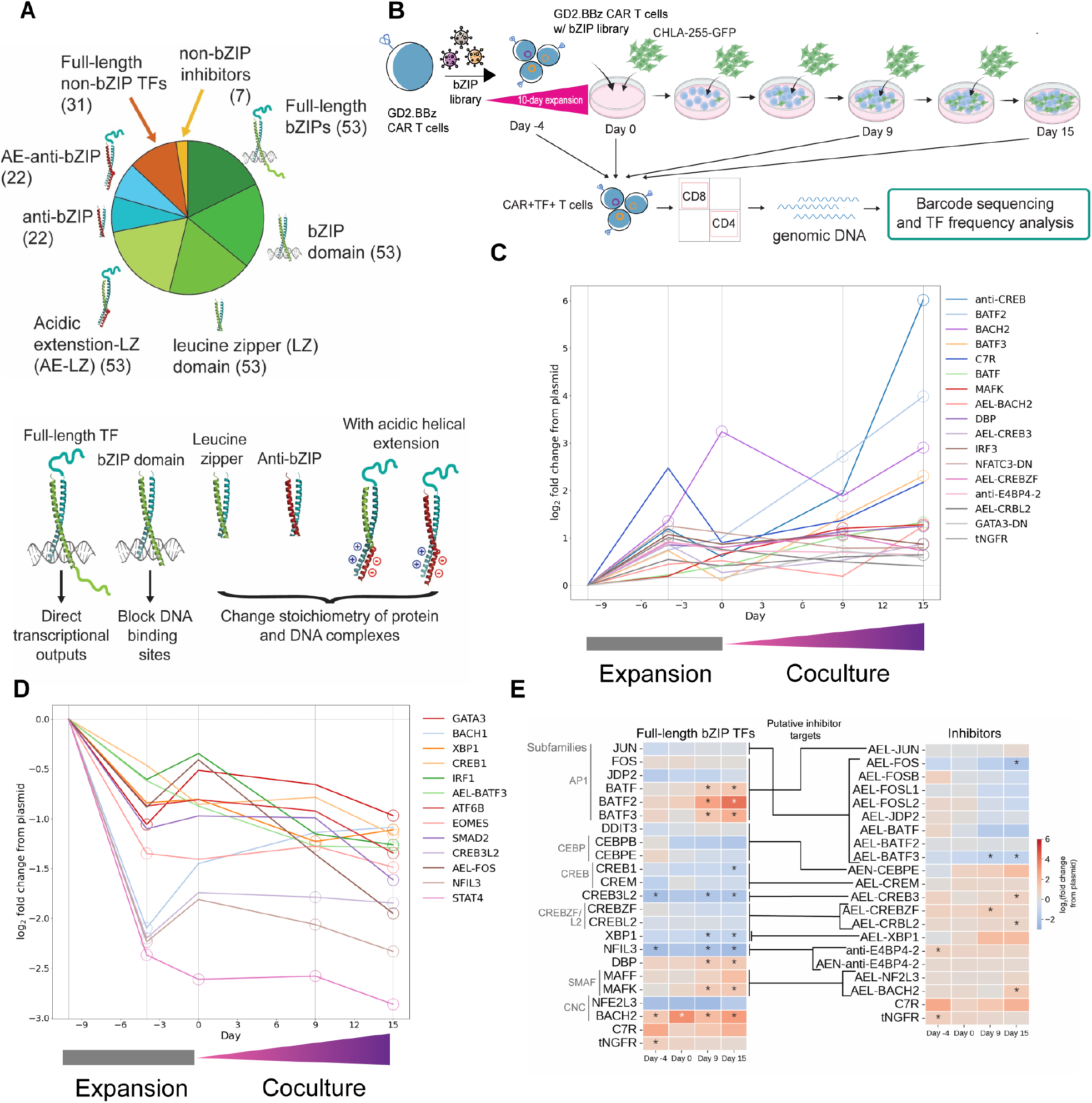
bZIP TFs and inhibitors from multiple subfamilies impact CAR-T cell expansion under exhausting conditions. **(A)** Schematic of bZIP variants and non-bZIP TFs included in the library. **(B)** bZIP library CAR-T cell expansion and persistence screen workflow **(C)** and **(D)** Log_2_ fold change relative to the frequency in the plasmid pool (Day −10) vs. time for bZIP variants significantly enriched **(C)** or de-enriched **(D)** in CD8^+^ T cells. Log_2_ fold change fit by n = 2 donors, 2 replicates and calculated via DESeq2; significance calculated via Wald test in DESeq2. Open circles indicate timepoints with p_adj_ < 0.05. (See Table S2 for all LFC values) **(E)** Comparison of the effects of full-length bZIPs and synthetic bZIP inhibitors on CAR-T cell expansion in the screen.

To generate a model of CAR T cell dysfunction *in vitro*, we benchmarked a GD2 CAR T cell repeated stimulation assay with a neuroblastoma cell line (CHLA-255.GFP) (Fig. 1B). GD2.BBz CAR T cells were selected as a model due to their moderate tonic signaling (*42*) and clinical relevance for multiple trials in our center, as well as the need for additional modifications to achieve durable responses (*39–41*). At later timepoints in the dysfunction assay (days 9-15), GD2 CAR T cells exhibited loss of proliferation and tumor killing (Fig. S2A-B). Repeatedly stimulated CAR^+^CD8^+^ T cells progressed toward a distinct transcriptional state (Fig. S2C), assuming a transcriptional profile associated with T-cell exhaustion (Fig. S2D), including upregulation of exhaustion markers and downregulation of effector and memory markers (Fig. S2E-G). Cumulatively, these data are evidence that GD2 CAR T cells in this assay are driven to exhaustion.

The library was transduced into GD2 CAR T cells and screened in the CAR-T cell dysfunction assay for variants promoting T-cell expansion (Fig. 1B). CAR^+^TF(tNGFR)^+^ T cells were sorted into separate CD8^+^ and CD4^+^ pools for barcode sequencing at day −4 during expansion (day 6 after transduction), and days 0, 9, and 15 of coculture. The time course enabled the identification of variants that promote or restrict T-cell expansion during *ex vivo* expansion and coculture. In CD8^+^ GD2 CAR T cells, the C7R positive control and multiple full-length TFs such as BATF2, BACH2, BATF3, and BATF were significantly upregulated relative to their starting frequency in the library plasmid pool. Notably, BATF2 promoted T-cell expansion in the screen to a greater extent than its paralogs BATF and BATF3, which have been previously reported as beneficial to T-cell expansion (*15*, *18*). Other novel hits enriched in the screen were anti-CREB, DBP, AEL-CREBZF, anti-E4BP4 (a NFIL3 inhibitor), and AEL-CRBL2 (Fig. 1C, Table S2). Other TFs such as CREB1, STAT4, and NFIL3 were strongly selected against, consistent with their established roles in promoting T-cell dysfunction (Fig. 1D, S3E) (*20–22*, *43*).

In the screen, full-length bZIP overexpression and inhibition via acidic extensions affected CAR T cell expansion more strongly than bZIP domains or leucine zippers/anti-bZIPs lacking acidic extensions (Fig. S3A-D). Including full-length bZIPs and their corresponding inhibitors let us directly compare overexpression and knockdown effects. JUN heterodimerizes with BATF and FOS subfamily bZIPs (Fig. S3F) (*26*) and acidic extensions fused to BATF/FOS-family leucine zippers should inhibit JUN DNA binding (*26*, *36*). Multiple acidic extension JUN inhibitors (e.g. AEL-BATF3, AEL-FOS, AEL-BATF) dropped sharply in frequency, suggesting that JUN inhibition is detrimental to T cell expansion. While JUN inhibition markedly impaired GD2 CAR T cell persistence, putative BATF and FOS inhibitors did not strongly affect expansion (Fig. 1E, Fig. S3D). Collectively, these inhibitor behaviors suggest that the amount of JUN is limiting for T-cell expansion, while BATF/FOS-family expression is in excess.

The TF variants significantly enriched or de-enriched in the CD8^+^ screen (Fig. 1C,D, S4A) followed a similar trajectory in CD4^+^ CAR T cells, though collection of fewer cells produced noisier data with only BATF2 enrichment reaching significance (Fig. S4B-D, Table S2).

### bZIP variants induce disparate phenotypic and transcriptomic effects

To prioritize bZIP inhibitors to validate, we performed trajectory analyses of TF expression in chronically stimulated GD2 CAR T cells (Fig. S5A) and overexpressed TF variant frequency in the screen (Fig. S5B). For enriched inhibitors, we looked for depletion of their full-length targets in the screen and an increase in target gene transcripts with exhaustion in the RNA-seq dataset. This analysis nominated putative inhibitors of related XBP1, CREBZF, and CREBL2, as well as inhibitors of CREB1-family members (CREB1, CREM, ATF1), NFIL3, and CEBPE/A/B/D. For initial validation, TF variants were transduced in parallel into GD2.mKate2 CAR T cells before chronic stimulation with CHLA-255.GFP tumor cells and tracking of CAR-T cell expansion and tumor killing via fluorescent protein. BATF2, BACH2, and the CREB1-family inhibitor AEL-CREM improved CAR-T cell expansion the most while JUN overexpression improved tumor killing (Fig. S5C-F). Several additional inhibitors, including putative inhibitors of NFIL3 and XBP1, improved proliferation when tested in a more rigorous flow cytometry-based assay (Fig. S5G-H).

Based on their differential effects on expansion and cytotoxicity in this initial assay, BATF2, JUN, and AEL-CREM were selected for further characterization. AEL-CREM, composed of an acidic extension fused to the LZ of CREM, was prioritized because CREB1 and CREM were de-enriched in the screen and are known contributors to T-cell dysfunction pathways (*10*, *11*, *20*). BATF2 overexpression significantly boosted CAR-T cell expansion, while AEL-CREM and JUN overexpression showed no significant improvement (Fig. 2A). Despite BATF2 improving CAR-T cell expansion, tumor killing mildly diminished, and neither JUN nor AEL-CREM improved tumor clearance (Fig. 2B). We performed Annexin V staining and CellTrace^TM^ Violet (CTV) dilution assays to assess the contribution of survival versus proliferation. The frequency of late-stage apoptotic (7-AAD^+^AnnV^+^) CAR T cells was higher for BATF2 and AEL-CREM (Fig. 2C), and all TF variants caused greater proliferation based on CTV dilution than CAR-only cells (Fig. 2D). These results are consistent with BATF2 and AEL-CREM stimulating frequent cell turnover.

**Figure 2.**
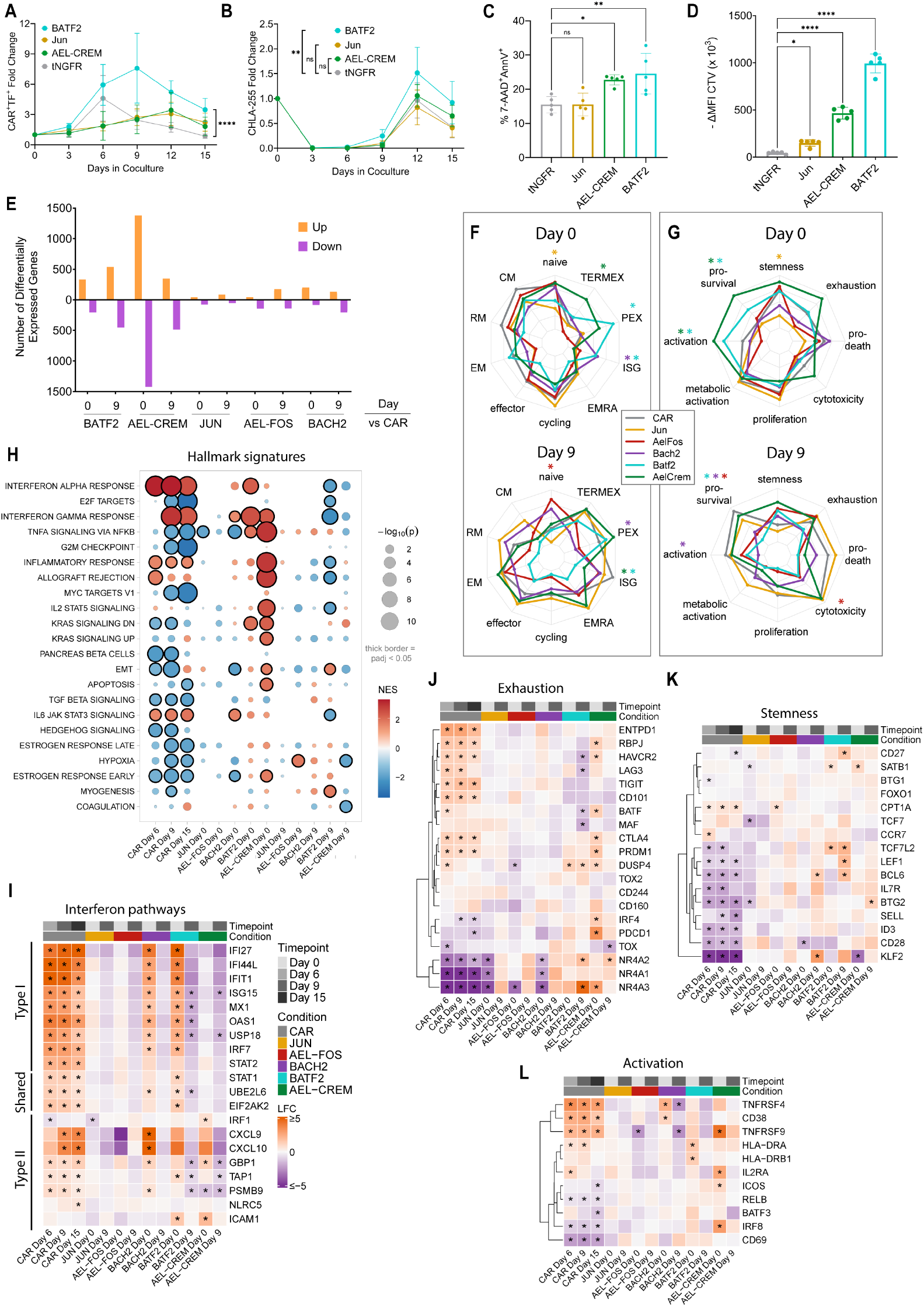
Parallel analysis of bZIP variants reveals pleotropic phenotypic and transcriptomic effects in CAR T cells. Fold change expansion of **(A)** TF-overexpressing CAR T cells and **(B)** CHLA-255.GFP tumor cells. n = 4-5 donors; error bars show SD; significance calculated by two-way ANOVA with Tukey’s adjustment for multiple comparisons. **(C)** Frequency of 7-AAD^+^/AnnV^+^ TF^+^/CAR^+^ T cells in culture. **(D)** CAR-T cell proliferation as measured by change in CellTrace^TM^ Violet (CTV) gMFI from baseline to 3 days after 1A7-idiotype antibody stimulation. Taller bars indicate more proliferation. (C,D) n = 5 donors; mean ± SD; significance from one-way ANOVA with Dunnett’s test for multiple comparisons. (A-D) ns = not significant, *p < 0.05, ***p < 0.001, ****p < 0.0001. **(E)** Number of differentially expressed genes for CD8^+^ CAR T cells expressing each TF versus CAR-only cells on the same day (n=3-5 donors, |LFC|>0.5, p_adj_<0.05 from DESeq2 Wald test with FDR correction). Radar plots of z-score normalized gene set variation analysis (GSVA) scores (mean across donors) of TF-overexpressing CAR T cells or CAR-only at day 0 or day 9 compared to **(F)** gene signatures derived from human tumor-infiltrating lymphocyte datasets (*96–98*) (linear mixed-effect model, *p_adj_<0.05), or **(G)** functional modules (linear mixed-effect model, *p<0.05). **(H)** Gene set enrichment analysis of Hallmark gene signatures on differentially expressed genes of contrasts: TF day 0 vs. CAR day 0, TF day 9 vs. CAR day 9, or CAR day 6, 9, or 15 vs CAR day 0. **(I-L)** Heatmaps of log_2_ fold change (LFC) values for gene expression changes (CAR day 6-15 vs. CAR day 0, TF day 0 vs. CAR day 0, or TF day 9 vs. CAR day 9) for the indicated pathways (*p_adj_<0.05 from DESeq2 Wald test with FDR correction).

To compare the transcriptional changes caused by overexpression of bZIP variants chosen based on their performance in the screen or their potential to yield mechanistic insight, we performed RNA-seq on CD8^+^ GD2 CAR T cells overexpressing BATF2, BACH2, JUN, AEL-CREM, or a JUN inhibitor, AEL-FOS, on day 0 (after an acute 3-hour stimulation) and day 9 of coculture. Transgene overexpression for TF variants was confirmed (Fig. S6A). For comparison, we evaluated the expression trajectories of endogenous bZIP TFs in control GD2 CAR T cells (Fig. S6B-E), noting a decline in JUN expression with duration in coculture, while BATF2 strongly increased with time in coculture, potentially consistent with its role as a type-I interferon-stimulated gene (*44*). BATF2 is rarely expressed at detectable levels in scRNA-seq datasets of human TILs (Fig. S16). Differential gene expression analysis revealed that AEL-CREM induced over 1000 differentially expressed genes (DEGs) in each direction, demonstrating the ability of acidic extension variants to potently alter gene expression (Fig. 2E, Table S4). BATF2 overexpression also induced widespread changes with a large number of DEGs on day 9, indicating both activating and repressing roles for BATF2. BATF2 and AEL-CREM samples also diverged modestly from the CAR-only samples by principal component analysis (Fig. S6F).

Analysis of DEG enrichment in or functional modules or T-cell state gene signatures derived from studies of human tumor-infiltrating lymphocytes (TILs) (Table S3) highlighted that BATF2 and BACH2 elevate an interferon-stimulated gene (ISG) signature on day 0 but BATF2 attenuates it on day 9 (Fig. 2F). AEL-CREM and BATF2 promote activation and survival expression modules at day 0 (Fig. 2G, S6G), perhaps indicating an attempt to resolve activation-induced stress. JUN promoted lower naive state and stemness module expression, while AEL-FOS and BACH2 moderately promoted these states (Fig. 2F-G). While blocking of JUN activity, either via direct binding of JUN (AEL-FOS) or competition with AP-1 sites (BACH2) (*17*, *36*), supports more naive T cell states, BACH2 diverged from AEL-FOS in some attributes, indicating functions independent of JUN competition.

To investigate the signaling pathways and TF activities behind these T-cell state changes, we analyzed DEG enrichment in MsigDB Hallmark pathways (Fig. 2H) and CollecTRI TF regulons (Fig. S6H) (*45*, *46*). JUN and AEL-FOS yielded few enriched pathways, likely due to the low number of DEGs. CAR-only cells strongly upregulated interferon alpha and gamma response signatures and TF regulons associated with inflammatory response (STAT1, IRF1/3) at days 6, 9, and 15 relative to day 0. BATF2 and BACH2 had already upregulated these inflammatory signatures at day 0 relative to CAR-only cells, echoing the elevated ISG state. BATF2 and BACH2 both upregulate type I-interferon and type II-interferon genes, while AEL-CREM boosted expression of type II-but not type I-interferon genes (Fig. 2I). Notably, BATF2 and AEL-CREM suppress interferon genes at day 9 (Fig. 2F,I). Prolonged type-I interferon signaling can contribute to exhaustion (*47*, *48*), so BATF2’s suppression of this pathway may contribute to its moderate anti-exhaustion effect (Fig. 2F,G,J). In line with a pro-persistence function for BATF2 and BACH2, they upregulate several stemness-associated genes at day 9 (Fig. 2K). BATF2 and AEL-CREM upregulated pathways (Fig. 2H; “TNFα signaling via NF-κB”, “IL-2 STAT5 signaling”), TFs and their regulons (Fig. S6H,I; JUN, FOS, NFATC1, NFKB1, RELA, STAT5A), and genes (Fig. 2L) associated with activation and effector activity at day 0, consistent with the role of BATF2 as an AP-1 member and the expected blocking of suppressive CREB1/CREM activity by AEL-CREM. Taken together, BATF2 and AEL-CREM promote gene signatures consistent with early inflammatory and activated states while BATF2 later promotes partial exhaustion resistance.

We compared the TF transcriptional effects to gene signatures drawn from BATF-family or CREM knockout studies in immune cells (Fig. S6J). BATF2 CAR-T cells were directionally consistent with genes upregulated by BATF- and BATF3-overexpressing T cell, but with overlap of only a few activation genes such as IL12RB2 and CD40LG (Fig. S6K). AEL-CREM T cells also had overlap with BATF/BATF3-upregulated genesets, probably reflecting common upregulation of activation genes (Fig. S6J). AEL-CREM mimicked the effect of CREM knockout in NK cells, with strong upregulation of genes repressed by CREM (Fig. S6J), suggesting that the inhibitor targets CREM. Collectively, these data demonstrate the strong effect of bZIP perturbations on T-cell states, including novel insights into overlap in the transcriptional program induced by overexpression of an AP-1 member, BATF2, an AP-1 competitor, BACH2, and a CREB1-family inhibitor AEL-CREM.

### Effects of JUN, AEL-CREM, BATF2 and paired combinations

Because solid tumors suppress CAR T function via multiple mechanisms, we hypothesized that pairing promising bZIP variants perturbing different dysfunction pathways would yield CAR T cells with superior properties. We overexpressed pairs of JUN, BATF2, and AEL-CREM in GD2 CAR T cells and tested them in the chronic CHLA-255 stimulation assay. Paired overexpression of BATF2 with either JUN or AEL-CREM did not result in additive benefit to CAR T cell expansion or tumor killing (Fig. 3A, Fig. S7A-B). Paired JUN-AEL-CREM overexpression outperformed each bZIP variant alone in augmenting tumor cell clearance without significant improvement in CAR-T cell expansion (Fig. 3A-B).

**Figure 3.**
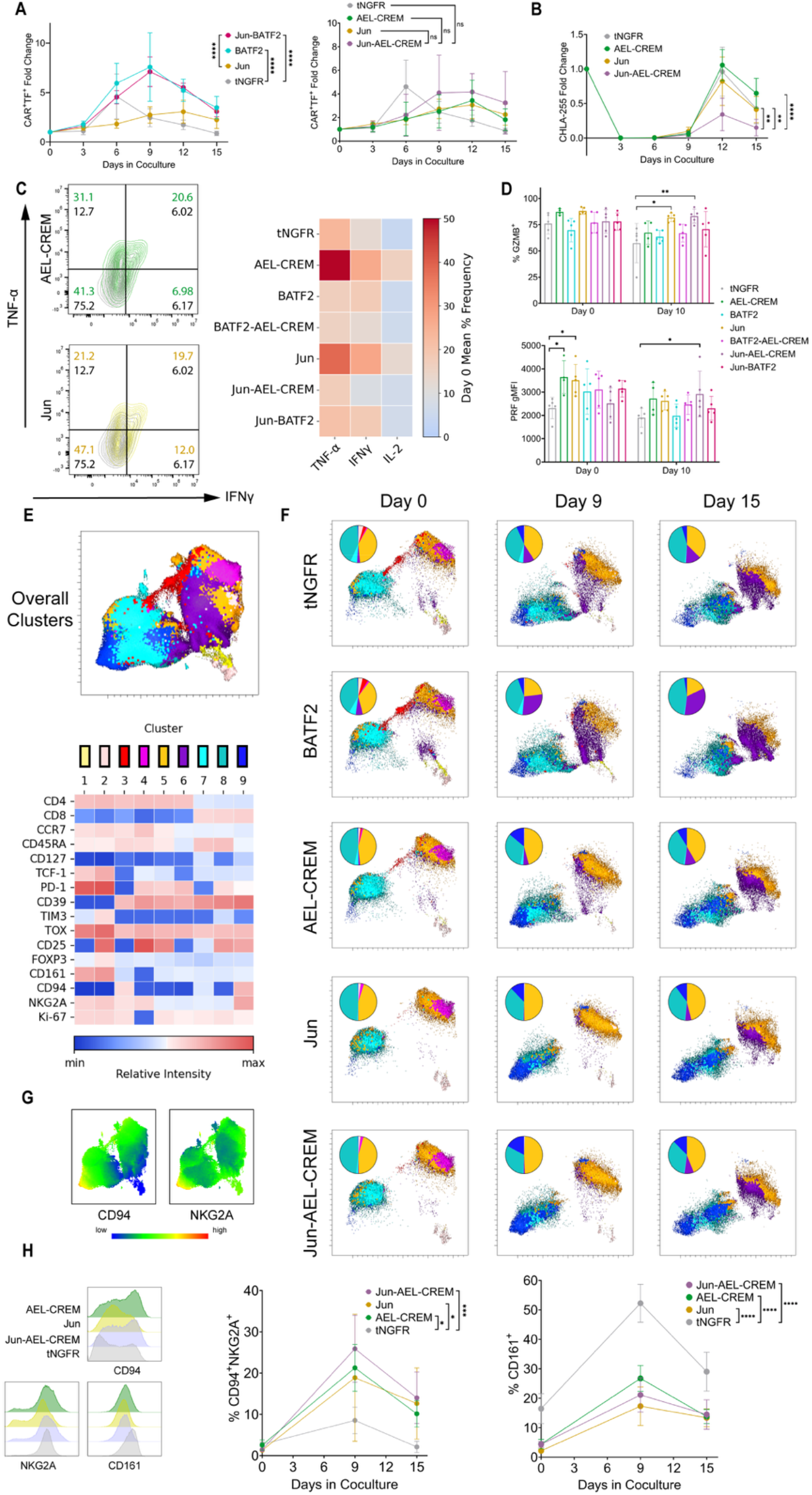
JUN, AEL-CREM, and BATF2 have strong but differential effects on proliferation and cytotoxicity. **(A)** Fold-change expansion of TF-overexpressing CAR T cells; n = 4-5 donors; error bars show SD; significance calculated via two-way ANOVA with Tukey’s test for multiple comparisons; **** p < 0.0001, *** p < 0.001, ns p > 0.05. **(B)** Fold-change residual CHLA-255 tumor cells relative to the tumor cells added for each round of stimulation; n = 4-5 donors; error bars show SD; significance calculated via one-way ANOVA with Tukey’s test for multiple comparisons; **** p < 0.0001, ** p < 0.01 **(C)** Representative flow plots of TNF-α vs IFN-γ and heatmap displaying the change in mean frequency of cytokines at baseline; n = 4-5 donors **(D)** Individual marker quantification of GZMB (% frequency) and PRF (gMFI); n = 4-5 donors, error bars show SD; significance calculated via one-way ANOVA with Dunnett’s multiple comparison’s test; **** p < 0.0001, *** p < 0.001, ** p < 0.01, * p < 0.05. **(E)** Clustered overall UMAP of spectral flow cytometry with relative intensity of each marker in each cluster displayed in the heatmap. Relative intensities are scaled according to the overall intensities of all markers. **(F)** UMAPs for TF variant conditions per timepoint. Pie charts display the frequency of each FLOWSOM cluster. **(G)** CD94 and NKG2A expression mapped onto the overall UMAP for all timepoints **(H)** Flow cytometry histograms (representative donor on Day 9 of coculture) and frequency in coculture for NK-associated markers in CD8^+^ CAR T cells; n = 5 donors; error bars show SD; significance calculated via two-way ANOVA with Dunnett’s multiple comparisons test; **** p < 0.0001, *** p < 0.001, * p < 0.05.

To further assess effector potential, we measured IL-2, IFN-γ, TNF-α, GZMB, PRF, and GNLY expression at baseline (day 0) and after 3 rounds of tumor stimulation (day 10). AEL-CREM and JUN elevated the frequency of cells producing IL-2, TNF-a, and IFN-γ at baseline (Fig. 3C, S7C). The substantial increase in these markers from AEL-CREM overexpression agrees with an increase in TNF-α/NF-κB, STAT5, and IFN-γ pathway activity noted in the transcriptome profiling (Fig. 2H). JUN and JUN-AEL-CREM overexpression increased the frequency of GZMB^+^ cells after repeat stimulation, and JUN-AEL-CREM increased the expression level of PRF (Fig. 3D). No significant differences in GNLY expression were observed in any of the TF conditions (Fig. S7C). Together, these results emphasize the proinflammatory effects of JUN and AEL-CREM and the association between improved GZMB/PRF expression and tumor killing upon paired overexpression of these bZIP variants.

To further characterize CAR T cells overexpressing these bZIP variants, we performed multiparameter spectral flow cytometry on days 0, 9, and 15 of coculture. FLOWSOM (*49*) clustering partitioned CAR^+^(tCD19^+^)TF^+^(tNGFR^+^) cells into 9 clusters (Fig. 3E). Clusters 1-4 were small CD4^+^ subsets present only at baseline, and cluster 5 comprised activated (CD39^high^CD25[IL-2Rα]^high^) CD4^+^ cells. Cluster 6 was CD4^+^CD25^low^, while non-activated (CD25^low^) CD8^+^ cells comprised cluster 7. Cluster 8 contained activated CD8^+^ Temra-like cells (CCR7^−^CD45RA^+^CD25^+^CD39^+^) and cluster 9 delineated NK receptor-positive CD8^+^CD94^+^NKG2A^+^ T cells (Fig. 3F, G). Consistent with the enhanced cytotoxicity of JUN-AEL-CREM, cluster 9 was enriched in JUN, AEL-CREM, and JUN-AEL-CREM conditions relative to CAR.tNGFR or CAR.BATF2 cells (Fig. 3B, F; S7D). CD94 and NKG2A dimerize and can act as inhibitory receptors upon binding tumor HLA-E to drive an NK-like dysfunctional state (*50*, *51*). However, NK-like CD8^+^ T cells have also been associated with increased cytotoxicity in tumors (*52*) and have been linked to effector T cells mediating transplant rejection (*53*). Because IFN-γ, which upregulates HLA-E in neuroblastoma (*54*), was low in our conditions during chronic stimulation (Fig. S7C), the JUN- and AEL-CREM-enriched CD94^+^NKG2A^+^CD8^+^ cells (Fig. 3H) likely represent a functional effector-like state. CD161 expression in CD8^+^ T cells, which is associated with NK-like dysfunction (*50*), was also reduced upon JUN and/or AEL-CREM expression during coculture, while CD25 trended upward (Fig. 3H, S7E), suggesting that JUN and AEL-CREM mitigate CD161-driven dysfunction while driving activation and cytotoxicity. Cluster 6 (CD4^+^CD25^low^) frequency increases with coculture for CAR.BATF2 T cells (Fig. 3F, Fig. S7D), which parallels how the fraction of CD25^+^ cells in the CD4 subset significantly decreased throughout the coculture (Fig. S7E). Potentially due to lower CD25 expression, which forms part of the high-affinity IL-2 receptor that drives proliferation (*55*), the overall frequency of CD4^+^ BATF2-overexpressing CAR T cells declined in coculture (Fig. S7F). Consistent with the BATF2 anti-exhaustion effect seen in the transcriptomic data (Fig. 2F-G, I-J), we also observed downregulation of CD161 but not CD94-NKG2A dimers in CD8^+^ cells (Fig. S7G). While BATF2 restrained exhaustion, it did not alter the fraction of CCR7^+^ memory CD8^+^ T cells (Fig. S7H). Notably, when BATF2 was paired with AEL-CREM or JUN, the BATF2 immunophenotype predominated (Fig. 3F, Fig. S7D), further demonstrating that BATF2 overexpression can override the effects of AEL-CREM and JUN. BATF2 overexpression thus improves expansion and counters exhaustion, but at the expense of long-term CD4 activation and persistence.

To check whether CAR T cell expansion remained cytokine dependent upon JUN/BATF2/AEL-CREM overexpression, T cells were cultured with or without IL-7 and 15. Upon cytokine withdrawal, all conditions failed to expand (Fig. S7I).

Finally, we assessed whether single and paired overexpression of JUN, AEL-CREM, and BATF2 would exhibit similar cytotoxicity effects in a different CAR T cell model of dysfunction (Fig. S8A). As in GD2 CAR T cells, JUN-AEL-CREM overexpression in GPC3.BBz CAR T cells boosted clearance of hepatocellular carcinoma HUH7.GFP cells, as did AEL-CREM and JUN alone (Fig. S8B). Collectively, these results demonstrate that BATF2 overexpression boosts exhaustion resilience and T cell expansion but does not benefit cytotoxicity. In contrast, JUN-AEL-CREM overexpression promotes a cytotoxic NK-like T cell state with elevated GZMB/PRF that is associated with superior tumor clearance.

### Canonical and non-canonical TF interacting partners of AEL-CREM and BATF2

To identify candidate protein complexes that underlie the phenotypic and transcriptomic changes observed upon CREM inhibition or BATF2 overexpression, affinity-purification mass spectrometry (AP-MS) experiments were performed with V5-tagged AEL-CREM and HA-tagged BATF2. V5-AEL-CREM complexes immunoprecipitated from GD2 CAR T lysates without tumor cell exposure and identified 176 significantly enriched proteins over IgG control (Fig. 4A, Table S5). As expected, V5-AEL-CREM bound all three members of the CREB1 subfamily with high confidence (Fig. 4B) (*26*, *56*). Unexpectedly, V5-AEL-CREM also bound JUN-family members JUN and JUNB, as well as ATF2-family members ATF2 and ATF7, although at a lower enrichment magnitude over IgG relative to CREM or ATF1. The pulldown of multiple members of bZIP subfamilies defined by homologous LZs with shared interaction profiles suggests that these interactions are likely mediated by direct LZ interactions. While the CREB1 family is not reported to interact with JUN-or ATF2-families in solution (*26*, *56*), a generic structural model (Boltz-2) (*57*) and two bZIP-specific, data-based models (*34*, *58*), show support for AEL-CREM or the CREM bZIP domain binding these subfamilies (Fig. S9A-E). Non-bZIP TFs of note include TBX21 (T-BET), which is known to promote an activated, effector state (*57*).

**Figure 4.**
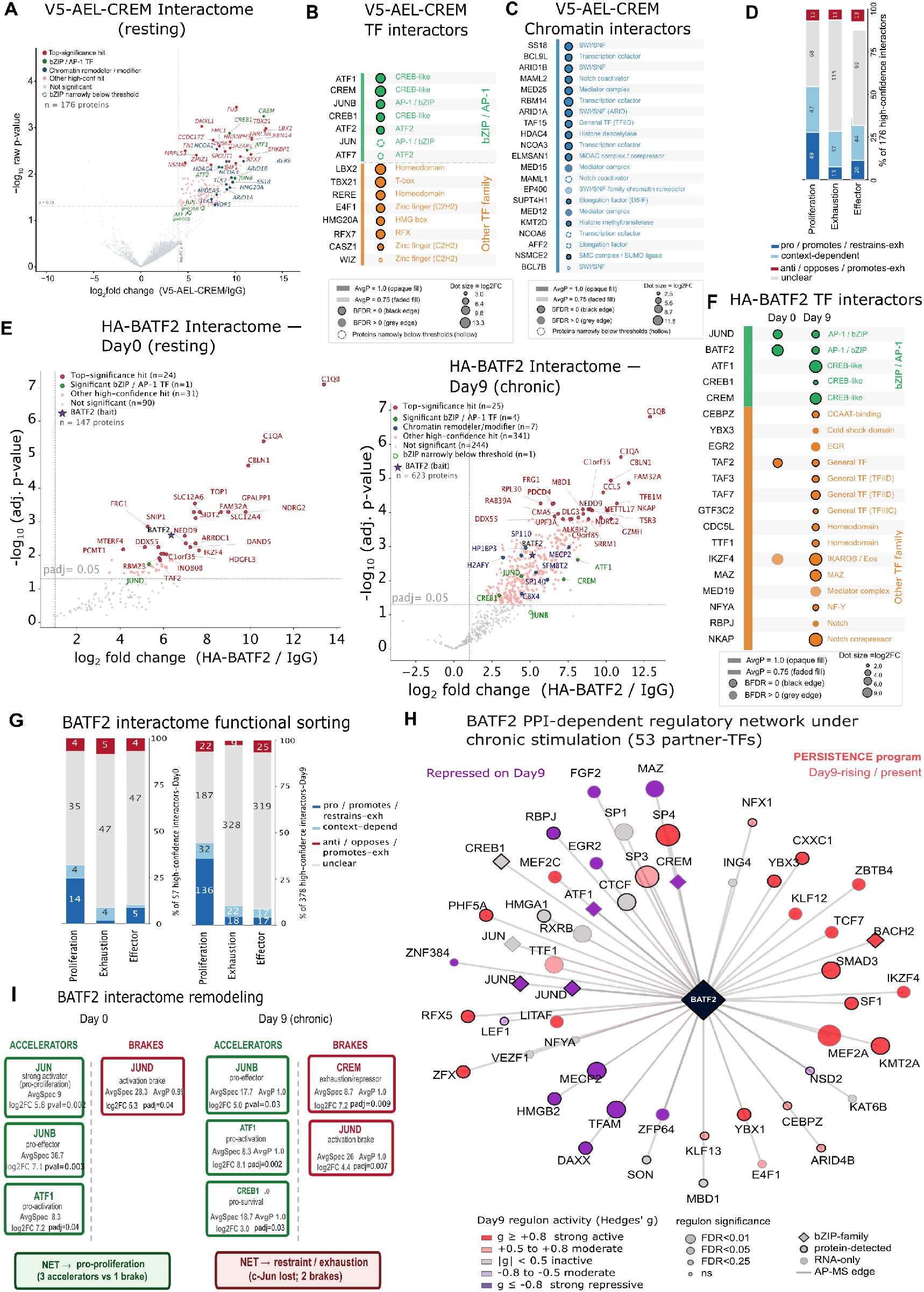
AEL-CREM and BATF2 protein interactomes in GD2 4-1BB CAR-T cells. **(A, E).** Volcano plots of the V5-AEL-CREM (unstimulated Day 0, **A**) and HA-BATF2 co-IP/MS interactome in GD2 CAR-T cells under resting (Day 0) or chronic (Day 9) stimulation with CHLA-255 cells **(E)**. Colored points denote significant interactors with the top 25 highest-confidence hits per condition labeled [(V5-AEL-CREM= SAINTq AvgP ≥ 0.75, BFDR < 0.05, log₂FC ≥ 4.0, p-value ≤ 0.05 vs. matched IgG), (HA-BATF2= log₂FC ≥ 1.0, p-adj ≤ 0.05 vs. day-matched IgG)]. AvgP= Avg probability of interaction, BFDR= Bayesian False Discovery Rate. **(B)** Dot plot of high-confidence AEL-CREM TF interactors and **(C)** chromatin factors (SAINTq AvgP= 1.0, BFDR<0.05). **(D)** AEL-CREM interactors categorized according to pro-or anti-proliferation, exhaustion, and effector functions. **(F)** Dot plot of high-confidence BATF2 TF interactors (SAINTq BFDR<0.05). **(G)** BATF2 interactors categorized according to pro-or anti-proliferation, exhaustion, and effector functions; TF annotations from (*94*) **(H)** BATF2 PPI-dependent regulatory network comprising TFs and chromatin modifying enzymes along with their respective regulon activities. CollecTRI decoupler-ULM ES metric Hedges’ g= bias-corrected standardized enrichment score to compare groups, measures transcriptional upregulation and repression. Edge= AP-MS copurification; node color indicates independently inferred Day 9 regulon activity and direction of activity shift (red= active, grey= inactive, purple= repressed on Day 9). bZIP nodes represented as diamonds. **(I)** A model proposing the balance of key TF interactions with BATF2 on day 0 that promote proliferation, and a subsequent handoff to the CREB1 promoting survival while combining with CREB and JUND brakes with a net result of restrained proliferation under chronic stimulation in BATF2 Day 9 CAR-T cells; Significant log2FC, AvgP and/or AvgSpec (average spectral counts) indicated.

The high-confidence V5-AEL-CREM interactome revealed additional interactions with TF cofactors and chromatin regulators predominantly associated with open chromatin and transcriptional activation (Fig. 4C). The pulldown of NCOA6 and NCOA3 coactivators, cBAF complex (ARID1A/B, BCL7B, and SS18) (*60*, *61*), and Mediator subunits (MED12, MED15, and MED25) (*62*), suggest that AEL-CREM associates with two major RNA Pol II transcriptional complexes (*63*, *64*). Both activating MAML1/2 and repressive HDAC4 appear, suggesting that at baseline, CREM and CREB1 complexes are poised for either activation or repression. Additionally, more interactors were associated with proliferative or effector states than exhaustion, as expected for resting T-cells (Fig. 4D). Thus, AEL-CREM physically associates with chromatin regulators that support open chromatin and transcriptional activation, facilitating a proliferative, effector state.

To assess how chronic stimulation reshapes the BATF2 interactome, we performed AP-MS on HA-BATF2 GD2 CAR T lysates before stimulation (day 0) and after three rounds of stimulation with CHLA-255 cells (day 9), which identified 147 and 623 proteins significantly enriched over IgG control, respectively (Fig. 4E, Table S6A). Within the bZIP family, BATF2 engaged in high-confidence interactions with JUND at day 0 and day 9, and, unexpectedly, with all three members of the CREB1 family at day 9 (Fig. 4F, Fig. 1A). At relaxed stringency, BATF2 interacted with all three JUN-family members (JUN, JUNB, JUND) at day 0, with day 9 marked by JUN loss and gain of modest BACH2 association (Fig. S10B). While unexpected, a BATF2-BACH2 bZIP domain interaction is reported to occur in solution at lower temperatures and is supported by our modeling (Fig. S9A-C) (*26*). The BATF-and CREB1-families are not reported to interact in solution, but two models show scores for interfamily interactions similar to those between BATF2 and BACH2 (Fig. S9A-C). BATF2 interaction with other TF families, cofactors, and chromatin regulators differed substantially between day 0 and day 9 (Fig. 4F, Fig. S10A). Consistent with classical AP-1 complexes, the day 0 HA-BATF2 interactome includes chromatin factors implicated in enhanceosome initiation (HMGB1) (*64*), and transcriptional activation (SUB1, BRD7) (*66*, *67*). By day 9, HMGB1 loss and HMGA1 gain suggest a shift toward a more stable locus-specific enhanceosome assembly (*68*). Furthermore, both activating and repressive transcriptional complexes are BATF2-bound: the MED19 Mediator and TFIID components (TAF2, TAF3, TAF7) alongside SMAD3-SKI, which can form a repressive complex with JUNB (*66*). Additional repressive components include MECP2, RBPJ, CREM, and BACH2.

Functional sorting of the BATF2 interactome revealed a greater association of pro-proliferation factors at day 9 and low recovery of pro-exhaustion and pro-effector factors (Fig. 4G). Additionally, gene set enrichment analysis (GSEA) on Hallmark pathways (*67*) of the HA-BATF2 interactome showed enrichment of Myc-V1/V2 pathways at day 0 consistent with an activation-driven translational burst, and a predominantly ribosome biogenesis program (Myc-V2) at day 9 (Fig. S10C). IFN-α/−γ signatures were enriched at day 0 and de-enriched by day 9, mirroring the transcriptional data (Fig. 2F,H,I, S10C). Further mapping of day 9 BATF2 interactors onto a curated TCR/CAR-proximal network (Fig. S10D, Supplementary Text 1) identified associations with high degree hub nodes CREB1, SMAD3, GZMB, GZMA, and RASA1, suggesting support for survival and modest effector activity alongside RASA1-mediated RAS inactivation (*68*), and SMAD3-associated restraint .

To assess whether TFs interacting with BATF2 were active or inactive, we quantified TF activity via Hedge’s g of CollecTRI TF regulons (*45*), for which negative or positive values indicate less or more inferred TF activity, respectively. Interestingly, the day 0 interactome is dominated by TF/chromatin factors with active transcriptional regulons whereas the day 9 regulons largely drop to baseline activity in CAR-only cells without corresponding changes to interacting TF expression (Fig. 4H, Fig. S11A-C). At day 9, active regulons include BACH2 and SMAD3, while JUNB/D, CREM, and ATF1 regulons are suppressed. Additional Hallmark analyses of mRNA and proteome subsets of high-contribution target genes, which contribute >90% of the transcriptome-derived regulon scores for individual BATF2-interacting active TFs (Fig. S11D, S11E) revealed enrichment of apoptosis, epithelial-mesenchymal-transition, TNF-α signaling via NF-κB, and IL-2-STAT5 signaling modules (Fig. S11F-G). BATF2-interacting TFs and their high-contribution target coverage at the proteome level are shown in Fig. S11H-I. Together, these findings suggest a shift from classical activating AP-1 complexes at day 0 to a broader mix of activating and repressive complexes at day 9.

### Interactions with BATF2 support persistence and limit exhaustion

Given the extensive remodeling of the BATF2 interactome during chronic stimulation (Fig. 4I), we examined interactors that might mediate proliferation or survival and resistance to exhaustion. As an AP-1 complex (*73*), BATF2-JUN cooperation is expected to support proliferation and effector functions of BATF2 CAR T cells. BATF2 transitioning from binding JUN to binding CREB1-members suggests a handoff from JUN-associated proliferation/effector activity to modulation of survival and exhaustion through the CREB1-family (Fig. 5A). Additionally, BATF2 is epistatic to JUN and AEL-CREM for proliferation and cytotoxicity effects, suggesting functional overlap (Fig. 3A, S7A, S8C). We evaluated the evidence for overlap on the proteome and RNA levels (Fig. S11H-J) using a variety of metrics, including GSEA NES to indicate target gene expression direction (Fig. S11K)and Hedge’s g of CollecTRI TF regulons to assess TF activity.

**Figure 5:**
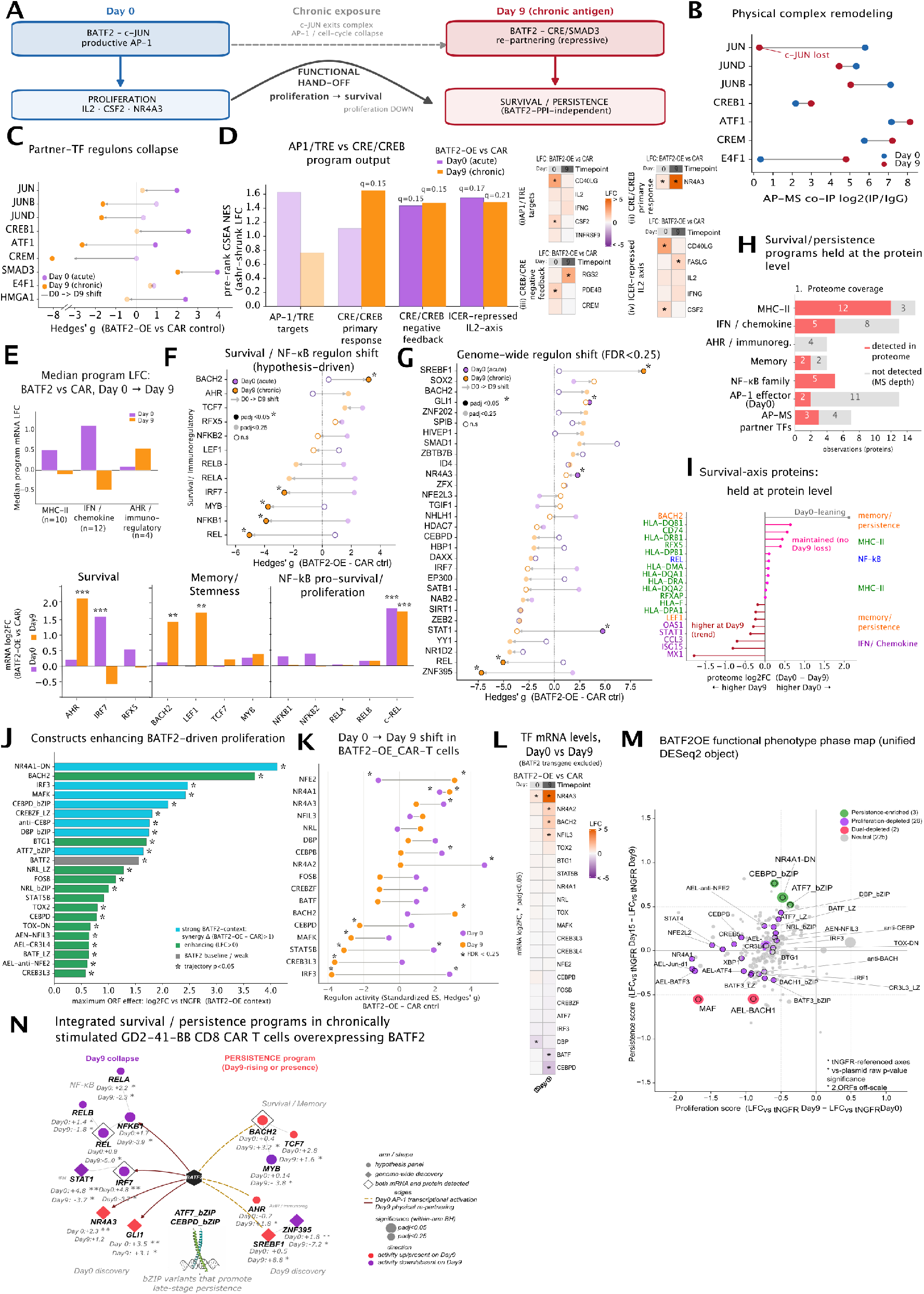
BATF2-PPI-dependent and PPI-independent transcriptional programs in human BATF2 GD2 CAR T cells. **(A-D)** BATF2-TF protein interactions: **(A)** Hypothesis: BATF2-PPI-mediated handoff from an early cJun-dependent effector (Day 0) to a Creb program (Day 9). **(B)** Day 0 to Day 9 shift in BATF2-PPI with Jun- and Creb1 subfamily members [SAINTq AvgP≥0.75, BFDR<0.05, SAINTExpress AvgSpec ≥10, BFDR <0.05, iBAQ log2(IP/IgG) ≥1.0, padj≤0.05]; E4F1= CRE-binding TF. **(C)** Regulon activity of BATF2-interacting TFs in Batf2-OE vs CAR control (CollecTRI-ULM, standardized enrichment score, Hedges’ g). **(D)** Program output by pre-rank GSEA (permutation FDR q <0.25 (solid), q≥0.25 (faded); ICER (Inducible cAMP Early Repressor). Right: top-ranked genes driving the per-day regulon shift. (E-F) Survival, memory and immunoregulatory TFs. **(E)** mRNA levels. **(F)** regulon activity (Hedges’ g, padj <0.25, left); **(G)** Genome-wide screen (ungated) for TFs with active regulons (right, Hedges’ g). **(H)** Protein coverage of survival/persistence program axes in the Batf2-OE Day 0 or Day 9 whole proteomes (LFC >1.0, p-value <0.05); **(I)** Enrichment trends on Day9 vs Day0. **(J-M)** BATF2 proliferation screen: enhancing constructs and regulon behavior of their TFs: TF combinations enhancing CAR-T/BATF2-OE proliferation by maximum ORF-LFCs across timepoint comparisons. **(J)** Enrichment (Δ LFC>0.5, green) or synergy (Δ LFC>1.0, teal)= max_TF_LFC (BATF2-OE – CAR) calculated for the Day 9 and Day 15; *p<0.05=Fisher’s combined trajectory P; **(K)** Day 0 to Day 9 shifts in TF regulon activity; **(L)** Corresponding mRNA levels; **(M)** Functional phase map: TFs positioned by pro-proliferative (ΔLFC_vs_tNGFR: proliferation, Day 9 - Day 0) and pro-persistence (ΔLFC_vs_tNGFR: persistence, Day 15 - Day 9) effects under chronic stimulation. Plasmid library is the DESeq2 reference; tNGFR is the post-hoc enrichment reference (see methods); Point significance: raw p < 0.05 vs-pooled plasmid library; **(N)** Integrated survival and persistence programs in chronically stimulated CAR-T/BATF2 cells. TF node Hedges’ g; exploratory ** and * FDR<0.05, <0.25, respectively. See also Supplementary Figure S15.

JUN association with BATF2 on day 0 (Fig. 5B) coincides with elevated JUN activity (Fig. 5C, Supplementary Text 2) and AP-1/TRE output driving canonical JUN-dependent effector/proliferation genes (Fig. 5D, Table S6). The inferred JUN activity decreases following dissociation on day 9 (Fig. 5B-D). Indeed, combined BATF2-JUN overexpression does not provide added proliferation to GD2 CAR T cells during chronic stimulation (Fig. 3A). CREM activity decreases with its gain of BATF2 association on day 9 and the IL-2 axis normally repressed by CREM/ICER retains activity (Fig. 5C, 5D). Consistent with this functional convergence, BATF2 and AEL-CREM show strong day 0-transcriptomic concordance (Spearman ρ = 0.65) and moderate AP-MS interactome similarity (ρ = 0.39, Fig. S12A-B) suggesting a degree of phenotypic overlap. Furthermore, the BATF2 and AEL-CREM AP-MS interactomes share 18 proteins (Fig. S12C), including the CRE-binding factors CREB1, CREM, ATF1, and E4F1 which are known to promote cell cycle progression and regulate survival (*74*). These CRE-factors gain BATF2 association on day 9 (Fig. 5B) while their inferred activities drop to baseline CAR levels (CREB1), decrease (CREM, ATF1) or stay moderately positive (E4F1) (Fig. 5C). Yet, the CRE/CREB program output remains elevated (Fig. 5D, S11K, Supplementary Text 2). Thus, BATF2 CAR-T cells preserve components of the CREB1 effector program while dampening CREM pro-exhaustion activity.

To identify TFs promoting persistence beyond the JUN- and CREB1-families, we evaluated TF mRNA levels (Fig. 5E, S11C,J, S13A-C), activity (Fig. 5F-G, S11A-B), and AP-MS interactome (Fig. 4F, S10A-B), and sought confirmation by evaluation of a whole proteome dataset from the same BATF2 day 0 and day 9 samples used for AP-MS (Fig. 5H-I, S13D, Table S7). This hypothesis-driven “persistence filter” identified: (a) an AHR immunoregulatory axis implicated in anti-PD1-mediated reinvigoration of precursor exhausted CD8+ T-cells (*71*, *72*), (b) RFX5-dependent MHC-II expression (*73*), (c) type-I IFNs under acute signaling which provide signal 3 necessary for memory formation, (d) a coordinated pro-memory module comprising BACH2, LEF1, TCF7, and MYB implicated in long-term persistence (*17*, *74–76*), and (e) NF-κB-mediated survival. Among these, while MHC-II/RFX5, interferon/IRF7 and NF-κB1/NF-κB2/c-Rel mRNA showed Day 0 induction (Fig. 5E, Fig. S13C), AHR and its canonical target CYP1B1, along with memory TFs BACH2, and TCF7, showed day 9-specific mRNA induction (Fig. 5E) and regulon activity (Fig. 5F). Proteome-level coverage of these programs is represented in Figs. 5H-5I, and Hallmark analyses of the proteome confirmed the enrichment of IFN-γ and TNF-α/NF-κB pathways (Fig. S13D). Complementary BATF2-PPI-independent and ungated analyses neither applying the BATF2-PPI filter, identified additional TFs that may support survival (Fig. Fig. S13A-B, 5G, Supplementary Text 3). Together, this integrated analysis implicated multiple programs potentially associated with long-term persistence of BATF2 GD2-CAR T cells.

We tested whether BATF2 overexpression locks proliferating GD2-CAR-T cells into a transcriptionally restrained state or whether co-expression of a second TF or inhibitory variant could relieve this restraint by repeating the bZIP library proliferation screen in the background of BATF2-OE GD2 CAR T cells. TF variants were evaluated by four criteria: (a) peak enrichment at any time point (LFC≥0.5, p<0.05); (b) trajectory-wide significance, Fisher’s combined p<0.05; (c) projection onto the proliferation-persistence axes to identify TFs promoting either proliferation (day 0-9 enrichment) or persistence (day 9-15 enrichment); (d) comparison with the original CAR-background screen to identify BATF2-specific effects (synergy Δ defined as enrichment over CAR-background screen). Comparing BATF2-CAR and CAR screens identified 22 bZIP/non-bZIP TFs and variants that enhanced proliferation with BATF2 (BATF2-CAR LFC>1.0 on days 9-15; trajectory p<0.05; Fig. 5J, Fig. S14A-B). Hits were consistent with rescue of suppressed regulon activity (CEBPD_bZIP, MAFK, STAT5B, CREB3L3, IRF3 and BATF, S14C-D) or inhibition of pro-dysfunction TF programs (NR4A1-DN targeting NR4A1/2/3) (Fig. 5K-L, S14E). Interestingly, consistent with gain of BATF2-physical association (Fig. S10B), and regulon activity on day 9 (Fig. 5F), BACH2 cooperates with BATF2 to further enhance long-term GD2 CAR T-cell proliferation (Fig. 5J, Fig. S14B) highlighting their functional interaction. Temporal comparisons within BATF2-OE background identified TF variants important for late persistence (NR4A1-DN, ATF7_bZIP) and negative regulators (MAF and AEL-BACH1) (Fig. 5M). Additional TF variants enhanced proliferation under chronic stimulation, defining a functional proliferation-persistence network (Fig. 5M; S15A-H; Supplementary Text 4). A network of experimentally derived nodes in the survival, proliferation, and persistence axes that enhance BATF2 function are summarized in Fig. 5N.

### BATF2 improves GD2 CAR-T cell expansion in vivo

To investigate the effect of BATF2 in an *in vivo* tumor model, we generated a B16F10 tumor cell line expressing the GD2 antigen (B16F10-GD2) by lentiviral transduction of GD3 synthase (*ST8SIA1*) and GD2 synthase (*B4GALNT1*) (Fig. S17A). A murine version of the GD2 CAR and human *BATF2* were transduced into T cells using murine gammaretroviral vectors, with GFP and Thy1.1 as transduction markers, respectively. To determine the effect of CAR and BATF2 on tumor control, mice first received a subcutaneous (s.c.) injection of B16F10-GD2, followed seven days later by CD3^+^ T cells transduced with GD2 CAR alone or co-transduced with BATF2 (Fig. 6A, Fig. S17B). CAR^+^ T cells or CAR^+^BATF2^+^ T cells were sorted before transfer (Fig. S17C). Mice receiving either type of CAR T cells exhibited significantly slower tumor growth (Fig. 6B, C) and higher survival rates (Fig. 6D) compared with the untreated control group. However, BATF2 expression did not provide an additional advantage in tumor control compared with CAR expression alone.

**Figure 6.**
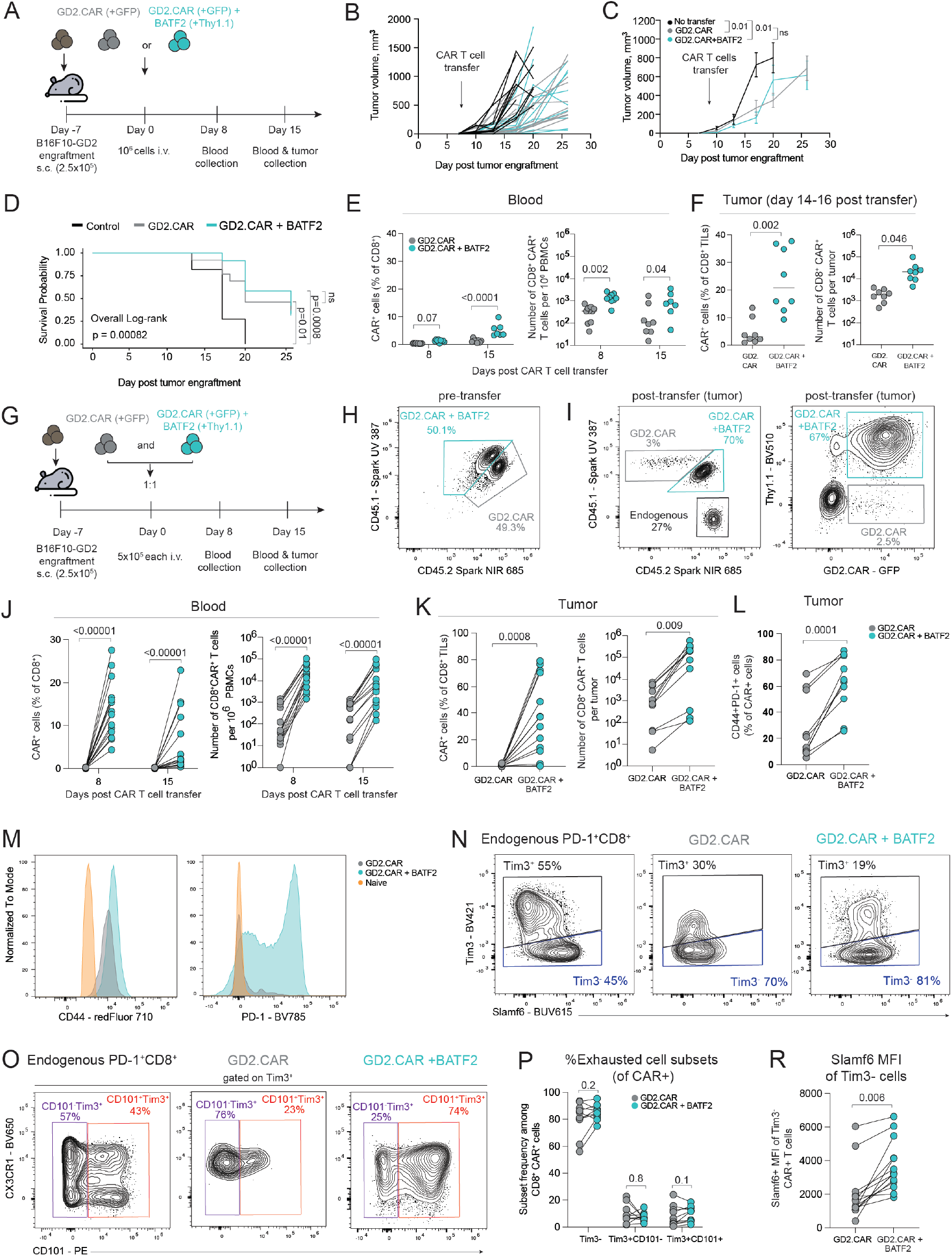
BATF2 effect in murine GD2 CAR T cells in melanoma model. **(A)** Schematic of the experiment to test control of B16F10-GD2 tumors by murine GD2 CAR T cells. **(B)** Individual tumor growth curves after transfer of 10^6^ CD3^+^ T cells transduced with either CAR (grey) (n=13) or CAR + BATF2 (turquoise) (n=12). The no-cell transfer group (black) serves as a tumor growth control (n=12). **(C)** Average tumor growth curves for the same experiment (mean±SEM). **(D)** Kaplan-Meier survival curves from mice with B16F10-GD2 tumor treated with CAR or CAR + BATF2. End point was a tumor size of 500 mm^3^. **(E)** Relative frequency and number of CAR^+^ CD8^+^ T cells in the blood and **(F)** in the tumor at days 8 and 15 post CAR T transfer. **(G)** Schematic of co-transfer of 5×10^5^ murine CAR and CAR + BATF2 T cells in B16F10-GD2 bearing mice. (H) Mix of CAR (CD45.1.2) and CAR + BATF2 (CD45.1.1) T cells before i.v. transfer to mice. **(I)** Representative staining and gating strategy by congenic markers and then based one GFP^+^ (CAR) or GFP^+^Thy1.1^+^ (CAR + BATF2) **(J)** Relative frequency and number of CAR^+^ CD8^+^ T cells in the blood (n=15) at days 8 and 15 and **(K)** in the tumor (n=11) at days 14-16 post co-transfer of the CAR or CAR + BATF2 T cells. **(L)** Expression of activation markers CD44^+^ and PD-1^+^ by CAR^+^CD8^+^ TILs (n=10). **(M)** Representative staining of CD44^+^ and PD-1^+^ by CAR^+^CD8^+^ TILs compared to naïve cells (orange). **(N)** Gating strategy to define Tim3^−^(dark blue), **(O)** Tim3^+^CD101^−^(purple), and Tim3^+^CD101^+^ (red) subsets. **(P)** Relative frequency of each subset within the indicated CAR^+^CD8^+^ T cell population: CAR (grey) (n=10) and CAR + BATF2 (turquoise) (n=10) in the tumor at day 15 post transfer. **(R)** Relative expression of Slamf6 in CAR^+^CD8^+^ TILs. All experiments were repeated 3 times with n = 3-5 mice per group. Statistics are two-way ANOVA (C, E), log-rank test (D), ratio t-test on log-transformed data (E, J, P), Welch’s t-test (F), paired t-test (K, L, R).

To determine whether BATF2 CAR T cells retained their enhanced proliferative capacity *in vivo*, we analyzed PBMCs and TILs for the frequency and number of CAR^+^CD8^+^ T cells. On days 8 and 15 after CAR T-cell transfer, BATF2 CAR CD8^+^ T cells exhibited significantly higher frequencies in PBMCs; this increased from day 8 to day 15 (Fig. 6E). In addition, BATF2 CAR CD8^+^ T cells showed a higher relative frequency and an almost tenfold greater number among TILs (Fig. 6F). There were no significant changes in CAR^+^CD4^+^ T cells in either blood or tumors (Fig. S17D,E). These findings are consistent with our *in vitro* data showing that BATF2 expression enhances CAR T-cell proliferation and persistence.

To investigate why the increased accumulation of BATF2 CAR T cells did not result in more effective tumor control, we performed an additional experiment in which CAR T cells and BATF2 CAR T cells were co-transferred into tumor-bearing mice (Fig. 6G). This model allowed us to characterize the immunophenotypes of both T cell populations within the same tumor microenvironment and under comparable antigen exposure. Murine CD3^+^ T cells carrying distinct congenic markers were sorted and mixed at a 1:1 ratio before transfer (Fig. 6H). Cells were first gated based on their congenic markers and subsequently on CAR (GFP^+^) and/or BATF2 (Thy1.1^+^) expression (Fig. 6I).

In the co-transfer model, BATF2 CAR CD8^+^ T cells exhibited an even greater proliferative advantage over CAR T cells in both the blood and tumors (Fig. 6J,K). By the final time point (day 15 post CAR T cell transfer), they accounted for up to 80% of CD8^+^ TILs, whereas CAR T cells were barely detectable. A similar increase in the relative frequency of CD4+ T cells was also observed in both the blood and tumors of mice receiving BATF2 CAR T cells (Fig. S17F). BATF2 CAR CD8^+^ TILs also exhibited significantly higher expression of the activation-associated markers CD44 and PD-1 (Fig. 6L,M).

To further characterize their differentiation, we examined expression of Slamf6 and Tim3, which characterize progenitor and terminally exhausted T cells, respectively. CAR^+^ T cells exhibited a lower relative frequency of Tim3^+^ cells and a markedly higher frequency of Tim3^−^ cells than endogenous TILs (Fig. 6N–P). Furthermore, BATF2 CAR CD8^+^ T cells showed a trend toward a higher relative frequency of Tim3^−^Slamf6^+^ cells, a phenotype associated with progenitor-exhausted T cells, consistent with their higher Slamf6 expression (Fig. 6R). The increased accumulation of CAR T cells may therefore reflect an expansion of the progenitor-like Slamf6^+^ population rather than an increase in the number of cytotoxic effector cells, pointing towards BATF2 promoting an activated, proliferative, but less-differentiated state.

### BATF2 and AEL-CREM reduce exhaustion in a chronic infection model

To determine if the effects of BATF2 and AEL-CREM are conserved in other exhaustion settings, we overexpressed them in P14 T cells, containing a TCR specific for the GP33 epitope of LCMV, transferred into mice infected with lymphocytic choriomeningitis virus (LCMV) clone 13, a chronic infection model (Fig. 7A). Human BATF2 or AEL-CREM were expressed with Thy1.1 as a transduction marker. We found that BATF2 and AEL-CREM reduced the frequency of terminally exhausted cells and biased CD8^+^ T cells to an intermediate state at day 15 post-infection (Fig. 7B-C). This effect persisted at day 30 (Fig. S18), including in the liver, a tissue where LCMV-specific T cells are typically driven towards the terminally exhausted state (*77*, *78*). Progenitor exhausted T cells were also significantly enriched in the liver at day 15 by BATF2 overexpression, in agreement with its effect in the murine tumor model; though, overall, we see more enrichment of the intermediate state in the LCMV model. The effect of BATF2 overexpression was particularly striking, leading to a distinct Tim3^low^CX3CR1^−^CD101^−^Tcf1^−^state. These cells had reduced expression of TOX, very low granzyme B expression, and lower levels of the Ki-67 proliferation marker in a tissue-dependent manner (Fig. 7D). AEL-CREM overexpression, in comparison, led to comparatively minor changes in these functional markers.

**Figure 7.**
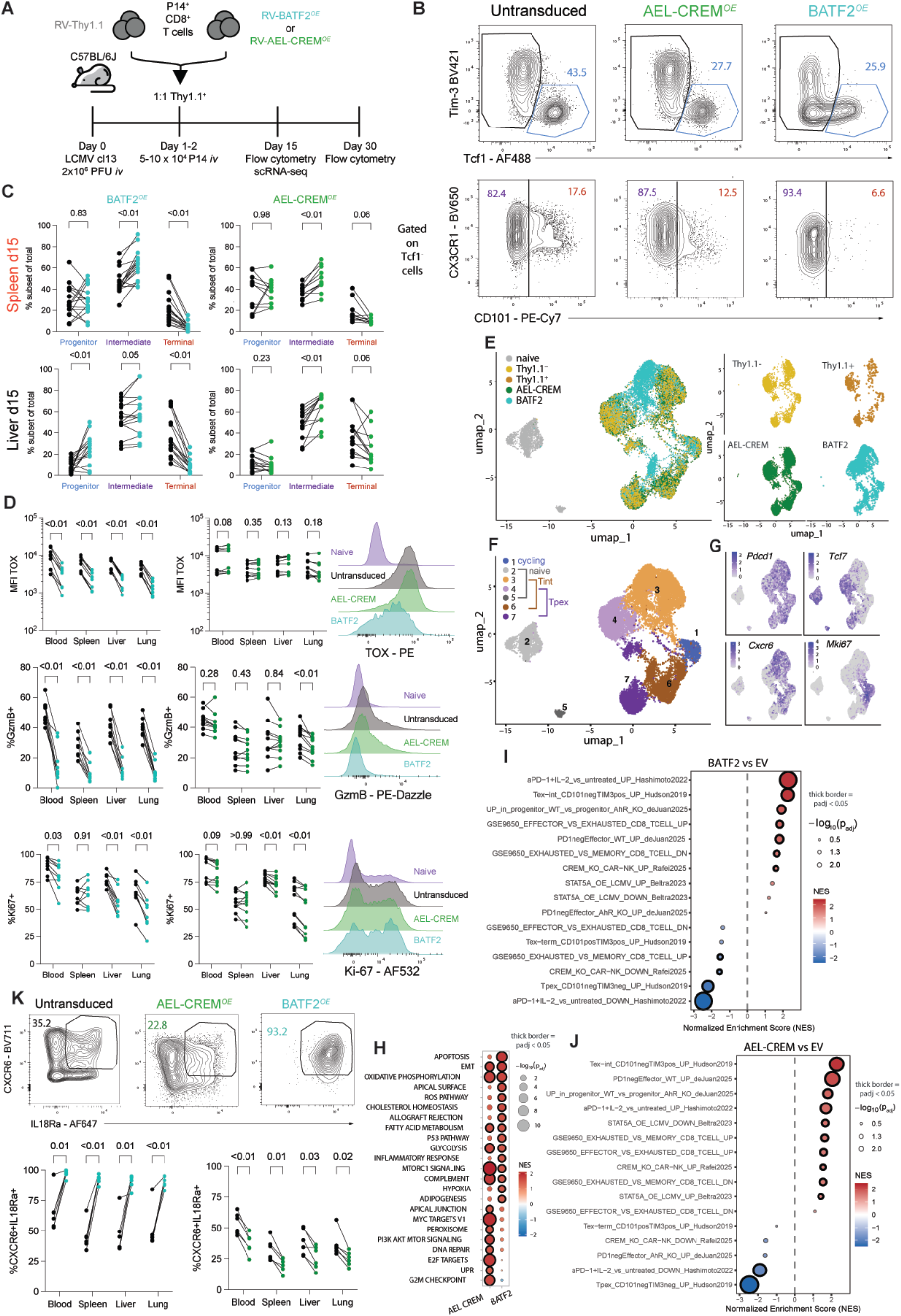
AEL-CREM and BATF2 overexpression in chronic LCMV model. **(A)** Schematic of P14 transfer experiments into mice infected with LCMV clone 13. **(B)** Gating strategy to define progenitor (blue, Tcf1^+^Tim3^−^), intermediate (purple, Tcf1^−^CD101^−^), and terminal (red, Tcf1^−^CD101^+^) subsets. **(C)** Relative frequency of each subset within the indicated P14 population: untransduced (black), AEL-CREM (green), or BATF2 (turquoise) in the spleen and liver at day 15. Data are pooled from 4 experiments of n=3-5 mice per construct. **(D)** Relative expression of TOX, granzyme B, and Ki-67 in the spleen at day 15 (left) with representative flow plots (right). Data are pooled from 3 experiments with n=3-5 mice per construct. **(E)** scRNA-seq UMAP of P14^+^ CD8^+^ T cells extracted from the spleen on day 15 of clone 13 infection, colored by condition. **(F)** UMAP colored by cluster, with mirrored clusters with similar transcriptomic profile colored in different shades: blue (cycling), purple (Tpex), brown (Tint), gray (naive). **(G)** Relative intensity of marker expression on the UMAP. **(H)** GSEA of MsigDB Hallmark gene signatures (*46*) on DEGs of pseudobulk AEL-CREM or BATF2 vs empty vector (EV, Thy1.1^+^). GSEA of DEGs from **(I)** pseudobulk BATF2 vs. EV or **(J)** pseudobulk AEL-CREM vs. EV against murine T cell signatures from ImmuneSigDB and select studies (*72*, *79*, *80*, *95–97*). Thick borders around bubbles indicate p_adj_ < 0.05 for **(H-J) (K)** Representative (top) and quantified (bottom) co-expression of IL18Ra and CXCR6, pooled from 2 experiments of n=3 mice per construct. All statistical comparisons for **(C-D,K)** were performed using matched 2-way ANOVA with Geisser-Greenhouse correction and Sidak’s multiple comparisons test.

To evaluate the effect of BATF2 and AEL-CREM on the transcriptome of murine T cells, we sorted P14 Thy1.1^+^ cells from spleens on day 15. We performed single-cell RNA-sequencing (scRNA-seq) on T cells overexpressing BATF2, AEL-CREM, or transduced with an empty Thy1.1 vector (including Thy1.1^+^ and Thy1.1^−^cells), as well as naive T cells from uninfected mice (17,504 cells passing filtering). AEL-CREM-overexpressing cells largely overlapped with Thy1.1^+/−^empty vector cells in a uniform manifold projection, while BATF2 cells enriched in distinct areas, indicating larger transcriptional changes (Fig. 7E). Clustering identified seven clusters (Fig. 7F, Fig. S19A), and analysis of key markers led us to annotate cluster 1 as cycling (*Mki67*), clusters 2 and 5 as naive, clusters 3 and 6 as intermediate exhausted (Tex-int, markers *Pdcd1*, *Cxcr6*, *Cx3cr1, Gzmb*), and clusters 4 and 7 as precursor exhausted (Tpex, markers *Pdcd1, Tcf7*, *Slamf6*) (Fig. 7F,G, Fig. S19B). All clusters except cluster 1 were split into two mirrored clusters based on differences in UMI counts and nuclear-encoded mitochondrial genes; therefore, we grouped clusters with shared profiles for downstream analyses.

Pseudobulk analysis of BATF2-or AEL-CREM-expressing cells versus empty vector Thy1.1^+^ cells (EV) revealed 1331 DEGs for BATF2 and 212 DEGs for AEL-CREM (Fig. S19C,D). GSEA performed on these DEGs against MsigDB Hallmark pathways showed that cells from both AEL-CREM and BATF2 were more metabolically active than Thy1.1^+^ cells, upregulating glycolysis, oxidative phosphorylation, and fatty acid metabolism pathways (Fig. 7H). This is consistent with promotion of a more active, effector state, which is also supported by upregulation of mTorc1 signaling, allograft rejection, and inflammatory response pathways. AEL-CREM appears to promote a more proliferative state than BATF2, as evidenced by significant upregulation of several cell cycle-related pathways (G2M checkpoint, E2F targets, DNA repair), which is consistent with their transcriptional profile in human CAR T cells (Fig. 2F,G). BATF2 and AEL-CREM expressing cells both show an enrichment in the Tex-int cluster (3_6) and a de-enrichment in the Tpex cluster (4_7) (Fig. S19E). Consistent with this, GSEA on murine T cell state signatures showed enrichment of a Tex-int signature and an active state induced by anti-PD-1 and IL-2 treatment, and de-enrichment of a Tpex signature for both TFs (Fig. 7I,J, Fig. S19F) (*79*). In agreement with the anti-PD-1/IL-2 study, BATF2 overexpression also led to high expression of CXCR6 and IL18Ra on the protein level, proteins associated with AP-1 activity (Fig. 7K). BATF2 DEGs computed within cluster 3_6 showed even stronger enrichment for functional effector states, including enrichment of a less exhausted Tex-int state reported to be caused by STAT5A overexpression (Fig. S19G-J) (*80*).

To directly compare the effect of BATF2 and AEL-CREM overexpression in murine T cells under chronic infection with their effect in human CAR T cells, we evaluated the enrichment of the same human TIL signatures and functional modules we analyzed against CAR T cells in Fig. 2F-G. This emphasized that both TFs cause enrichment of effector and cytotoxic states, while AEL-CREM looks modestly more exhausted (EMRA and TERMEX states) and cycling (Fig. S19K). Both TFs significantly enrich the resident memory (RM) state, which together with their enrichment of the epithelial mesenchymal transition (EMT) pathway, suggests a role in promoting adhesion and tissue residency programs, which is supported by upregulation of several tissue residency-associated genes (Fig. S19K,L).

To evaluate BATF2’s effect more closely, we compared it to gene signatures derived from published studies of BATF or BATF3 overexpressed in murine T cells (*15*, *18*), and the available BATF2 studies in macrophages and hematopoietic stem cells (HSCs) (*85*, *86*). We also generated gene signatures for up-or downregulated genes from our BATF2-overexpressing human CAR T cells at day 0 or day 9 of coculture relative to CAR-only cells. This revealed that BATF2 in murine T cells regulates a program at least partially overlapping that of overexpressed BATF and BATF3 (Fig. S19M,N). The genes bound by BATF2 in HSCs or repressed by BATF2 in macrophages were downregulated or upregulated, respectively by BATF2 overexpression in T cells, indicating differences in transcriptional programs in these cell types. The BATF2 DEGs in murine cells were counter-aligned with the gene signatures from our human BATF2-CAR T cell dataset, with the day 9 gene set being strongly inversely correlated. These results point towards BATF2 acting as a traditional AP-1 factor in murine T cells, driving a functional effector state and reducing exhaustion, which is more in line with its transcriptional profile at day 0 for the CAR T dataset, where it also promoted an activated state.

## DISCUSSION

We employed a pooled library screen containing full-length bZIPs and protein-based inhibitors to identify bZIP variants that promote superior CAR T cell proliferation or persistence. The screen and subsequent validation nominated full-length bZIPs with known benefits to T-cell expansion including BACH2, BATF, and BATF3, and new pro-expansion TFs such as BATF2 and DBP. Pro-expansion inhibitors included AEL-CREM, and several other putative inhibitors of NFIL3, the CEBP family, and XBP1-related bZIPs. Recognizing the importance of bZIP interaction partners to transcriptomic and phenotypic outcomes, we used AP-MS to assess partners of our novel hits BATF2 and AEL-CREM, integrating it with transcriptomic and phenotypic analysis to understand the mechanism underlying their pro-expansion effect. We found that BATF2 overexpression in CAR T cells selectively enhances proliferation without a concomitant increase in cytotoxicity-marker expression or tumor killing in vitro or in vivo indicating an uncoupling of effector functions. Despite enhanced activation markers in vitro and in vivo, which is consistent with its role as an AP-1 member, BATF2 restrains exhaustion in human CAR T cells and promotes a progenitor exhaustion state in murine CAR TILs and an intermediate state in the LCMV model. This was in contrast to the effect of JUN overexpression in human CAR T cells, which yielded reduced stemness and more cytotoxicity, with little effect on exhaustion markers. BATF2’s restraint of terminal differentiation may be due in part to its protein-protein interactions with BACH2 and CREB1-family members. This may explain the overlap in phenotypic outcomes with AEL-CREM, which also binds and blocks the CREB1 family and promotes increased proliferation and restrains terminal differentiation in human CAR T cells and P14 cells. These findings highlight how phenotypic outcomes of bZIP perturbations arise from altered to bZIP complexes and emphasize the utility of our approach comparing overexpression of full-length bZIPs with inhibitors that block specific interactions.

Overexpressed BATF2 appears to have partially overlapping, but mostly distinct effects from its family members BATF and BATF3, which are endogenously highly expressed and better studied in CAR T cells (*15*, *18*, *24*). While endogenous BATF2 expression tends to be low in T cells (*87*, *88*), we detected an increase in expression during chronic stimulation, suggesting an endogenous role in CAR T cells. In general, we observed that BATF2 overexpression is more similar in its effects to BATF than BATF3 overexpression, both in the context of chronic tumor and viral stimulation. Both BATF and BATF2 are associated with pro-effector roles, promoting type I interferon and IL-12 signaling (*15*) but without strong effects on memory markers like IL7R associated with BATF3 (*18*). BATF2 is a type-I interferon response gene activated by STAT1/2 in CD4^+^ T cells (*83*), which parallels our findings in the inflammatory CHLA-255 coculture assay. BATF2 and BATF overexpression also share tissue residency gene signature which appeared in human BATF2-GD2 CAR T cells and BATF2-P14 T cells and was previously reported in BATF-overexpressing TILs (*15*), indicating a tissue-adapted effector state. During acute stimulation, BATF2-overexpressing CAR T cells upregulated pro-activation/effector genes also found upregulated in BATF-overexpressing TILs. BATF2 has a longer C-terminal extension of unknown function beyond the bZIP domain than BATF and BATF3, and this C-terminal segment was a frequent hit in strongly enriched variants in recent persistence screens of chimeric bZIP TFs (*31*). In that study, overexpressed BATF2 and some of the BATF2 C-term variants displayed a strong pro-memory effect in CD19 CAR T cells, which we see elements of, but may be offset by the stronger tonic signaling and pro-inflammatory environment in human GD2 CAR T cells cultured with solid tumor lines. The pro-persistence and -memory effect of BATF2 was emphasized in murine GD2 CAR T cells in the B16-GD2^+^ model, in which BATF2 overexpression greatly increased CAR T cell counts in the tumor and memory marker *Slamf6* expression. These progenitor-like cells have self-renewal potential but may exhibit limited immediate effector function, while their capacity for self-renewal can support responses to immune checkpoint blockade (*84*). The strong proliferative and anti-exhaustion effect of BATF2 overexpression was mirrored in the LCMV model, but there we observed a more activated, intermediate-exhaustion state, emphasizing a potential sensitivity to CAR versus TCR signaling or inflammatory states. In all CAR T models in our study, BATF2 displays no benefit in cytotoxicity and tumor control despite higher T cell counts, which aligns with the cytotoxic deficit induced by overexpression of BACH2, suggesting functional overlap (*17*, *85–87*). Thus, despite the anti-exhaustion, pro-proliferation, and -persistence state induced by BATF2 overexpression in tumor and chronic viral infection models, these states appear to be divorced from benefits to cytotoxicity.

Besides interactions with the Jun subfamily, overexpressed BATF2 exhibited unexpected complexes with CREB1-family bZIPs and BACH2 during chronic stimulation, which may mediate alternate transcriptional programs that tune gene expression to temper activation and avoid exhaustion. Moreover, we observed a masking effect when pairing BATF2 with JUN or AEL-CREM for proliferation, tumor killing, and immunophenotype, which suggests that overexpressed BATF2 overrides the effects of JUN and AEL-CREM at least in part through direct modulation of JUN- and CREB1-family complexes. Whether BATF2-CREB1-family complexes bind DNA and affect gene expression remains to be tested, but considerable overlap in the transcriptional profile of BATF2 and AEL-CREM, which is expected to block DNA binding through AE-basic domain interactions (*37*, *38*), suggests that the effect of overexpressed BATF2 on CREB1-family activity may be through blocking of protein-DNA complexes. BATF2’s strong pro-persistence effect may be aided by the upregulation of BACH2 and potentially direct effects of BATF2-BACH2 complexes, and our integrated transcriptomic and proteomic analysis identified several other BATF2-PPI-independent pathways that may contribute to this effect. These results highlight how overexpression of bZIP TFs can alter their interaction partners and have substantial transcriptomic and phenotypic effects.

AEL-CREM overexpression exhibited a more modest effect on proliferation than BATF2 overexpression in CAR T cells (Fig. 2D), but transcriptomic profiling detected more proliferative pathway upregulation in CAR T cells and P14 T cells. AEL-CREM promoted a tissue-adapted effector state in chronic viral and tumor coculture models. In addition, AEL-CREM overexpression, as well as JUN or the JUN + AEL-CREM pair, promoted an NK-like cytotoxic state in GD2 CAR T cells and improved CAR-T cell tumor clearance *in vitro*. The fact that pro-dysfunction CREM is upregulated by PGE_2_ and catecholamines in the tumor microenvironment (*10*, *88*, *89*) makes AEL-CREM an attractive candidate for application in T-cell therapies for solid tumors. We note that based on the AP-MS data and computational modeling, AEL-CREM also binds weakly to JUN and ATF2 subfamily bZIPs, which warrants further optimization to avoid blocking JUN, which we conclude is a limiting AP-1 component with respect to T-cell function. Inhibiting multiple members of the CREB1 subfamily in T cells could be beneficial if all members contribute to dysfunction in solid tumors, which is a benefit of protein-based inhibitors relative to making multiple CRISPR edits (*90*).

Our results showing the anti-exhaustion potential of several bZIP inhibitors motivate future work to identify inhibitors of bZIPs responding to other sources of T-cell dysfunction in the tumor microenvironment, including reactive oxidative species, ER stress, or nutrient deficiencies (*2*, *13*, *14*, *19*, *23*, *25*), which may not be adequately modeled in our *in vitro* exhaustion assay. Our investigations were in the context of overexpression, which likely alters bZIP interaction stoichiometry relative to endogenous levels, and highlights the potential for rewiring bZIP transcriptional pathways through overexpression and inhibition. The evidence that overexpressed BATF2 alters competition between AP-1, CREB-family, and BACH2 complexes warrants further investigation with endogenous bZIPs including the other BATF-family members.

Overall, this study identified novel ways that both full-length bZIPs and bZIP inhibitors can regulate T-cell states of proliferation, cytotoxicity, activation, persistence, and exhaustion. While we deeply characterized overexpression of BATF2, JUN, AEL-CREM, and paired combinations thereof, other bZIP variants highlighted in this study warrant further investigation. The number of promising bZIP variants identified in this study underscores the potential of bZIP engineering as a promising avenue for engineering T cells with exhaustion resistance.

## Supplemental Figures

**Figure S1.**
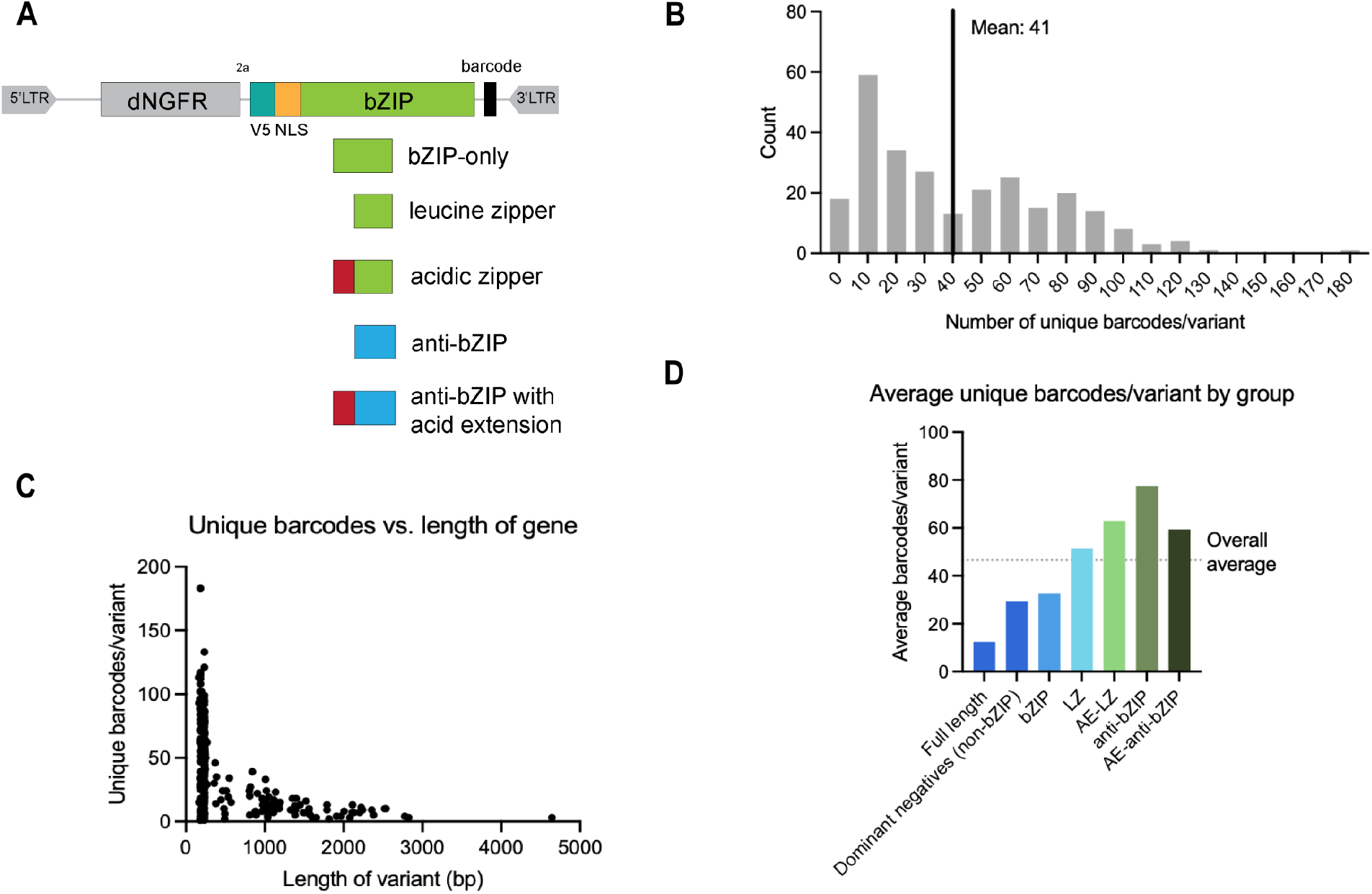
Library construct design and quality control of bZIP variant library. **(A)** Schematic of bZIP library constructs used in the screen **(B)** Distribution of the number of unique barcodes per variant **(C)** Number of unique barcodes versus the length of the library variant **(D)** Average unique barcodes per variant, organized by variant category

**Figure S2.**
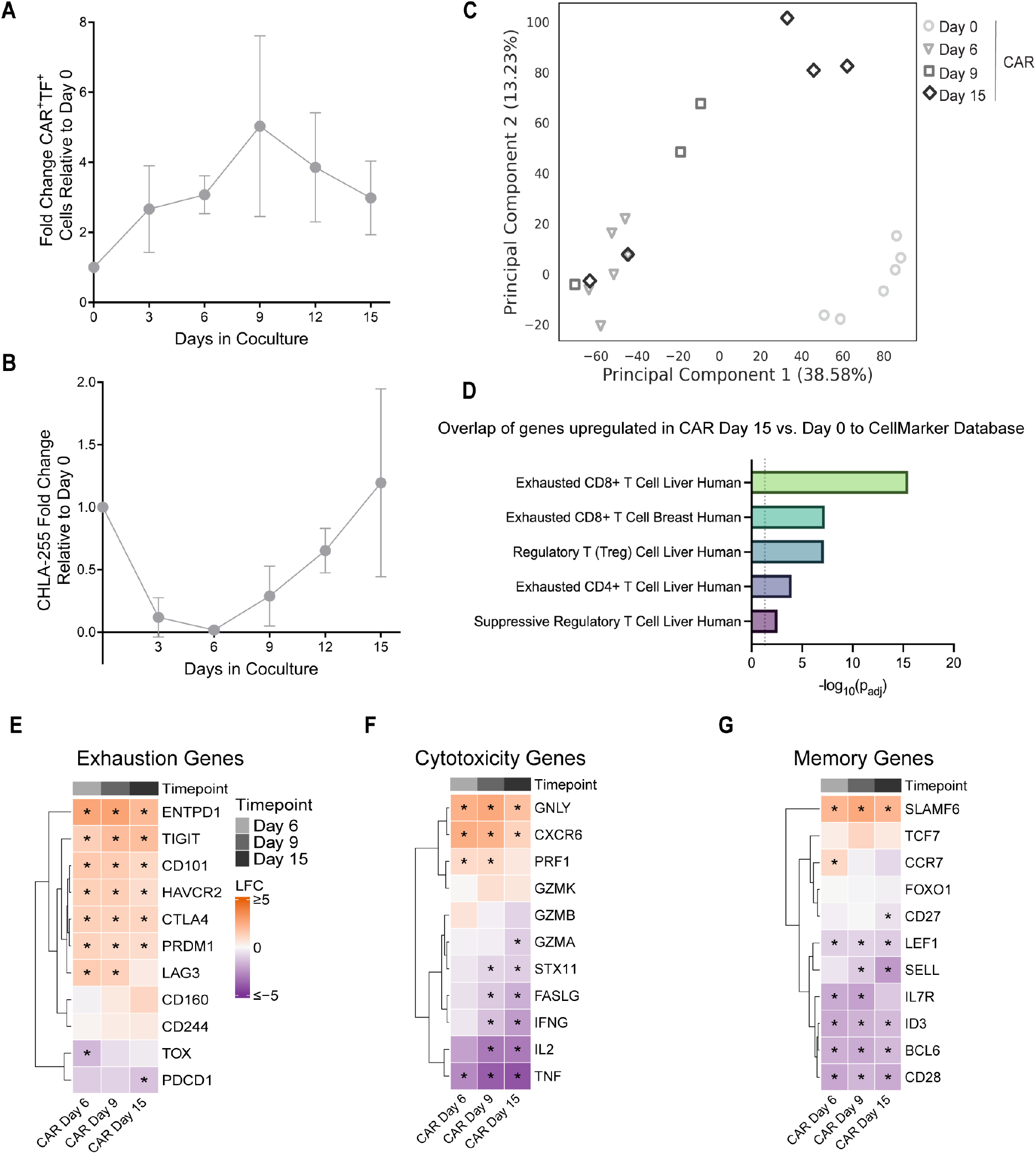
Validation of GD2 CAR T cell in vitro exhaustion assay. **(A)** Fold change of CAR.tNGFR T cell expansion; n = 4 donors; error bars show SD **(B)** Fold change of CHLA-255 tumor cells; n = 4-5 donors; error bars show SD **(C)** PCA plot showing onset of T-cell dysfunction in GD2 CAR-T cell transcriptomes over time in coculture (n= 3-6 donors) **(D)** CellMarker (*103*) gene signatures with significant overlap to genes upregulated in coculture (Day 15 vs. Day 0) **(E-G)**. Heatmaps of log_2_ fold change (LFC) values for gene expression changes (CAR day 6-15 vs. CAR day 0) for the indicated pathways (*padj<0.05 from DESeq2 Wald test with FDR correction).

**Figure S3.**
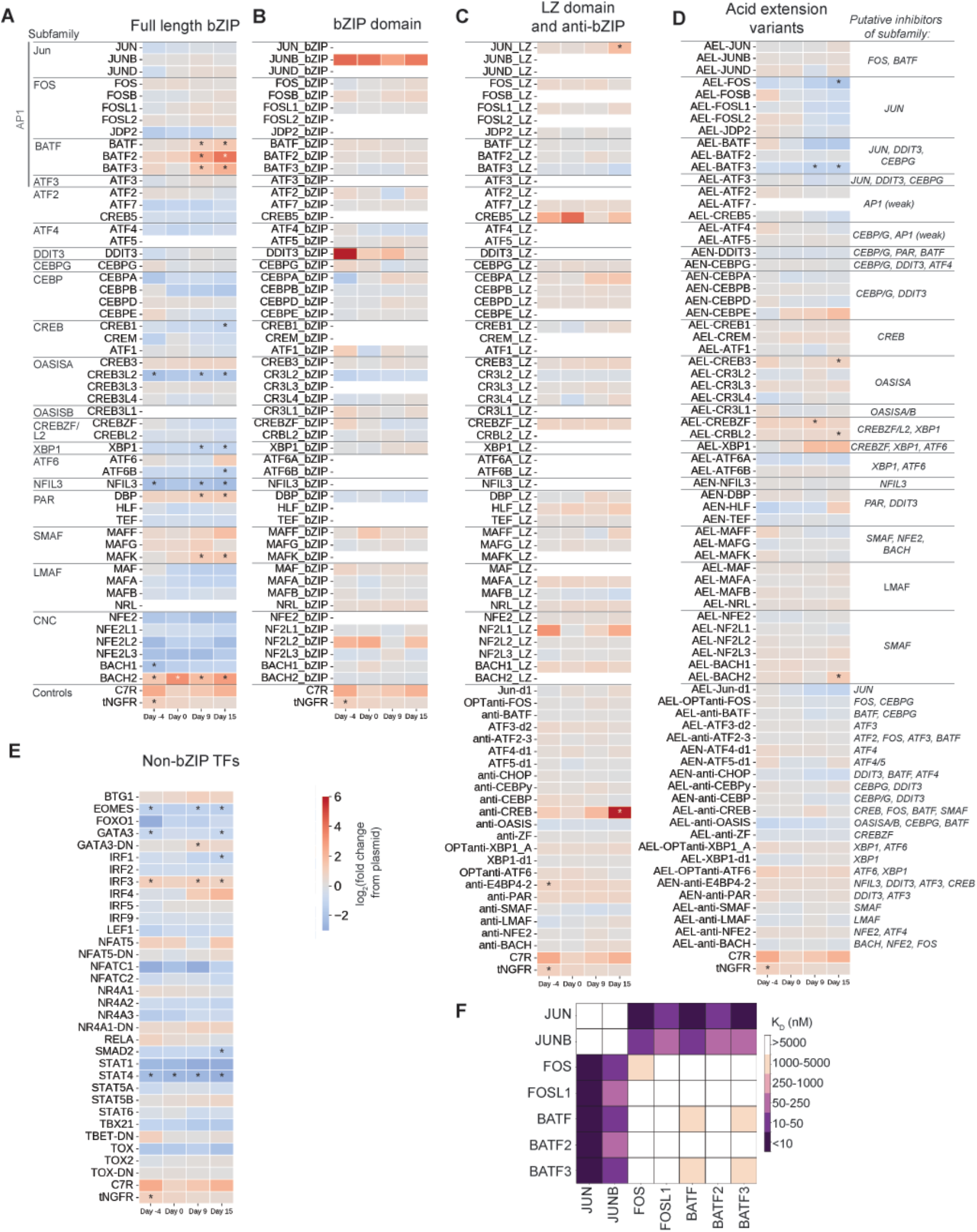
Library results in CD8^+^ GD2 CAR T cells emphasize stronger effect for full-length TFs and acidic extension inhibitors. Change in TF variant abundance across expansion (day −4) and CHLA-255 coculture timepoints (Day 0-15) for the enrichment screen in GD2 CAR T cells for: **(A)** full-length bZIP TFs, **(B)** bZIP domain-only variants, **(C)** leucine zipper (LZ) and anti-bZIP variants, **(D)** acidic extensions on LZ and anti-bZIP variants, and **(E)** non-bZIP TFs and putative dominant negatives (DN). Horizontal lines separate bZIP variants by subfamilies, labeled in (A). Putative target subfamilies of the acidic extension inhibitors are listed to the right in (D), based on *in vitro* binding data for bZIP domains and anti-bZIPs (*26*, *32*). White spaces in the heatmaps indicate variants that dropped out of the library screen **(F)** *In vitro* interaction matrix for AP1 bZIPs reproduced from (*26*). (A-E) log_2_(fold-change) in variant abundance relative to frequency in library plasmid fit from n=4 biological replicates; * denotes adjusted p-value < 0.05.

**Figure S4.**
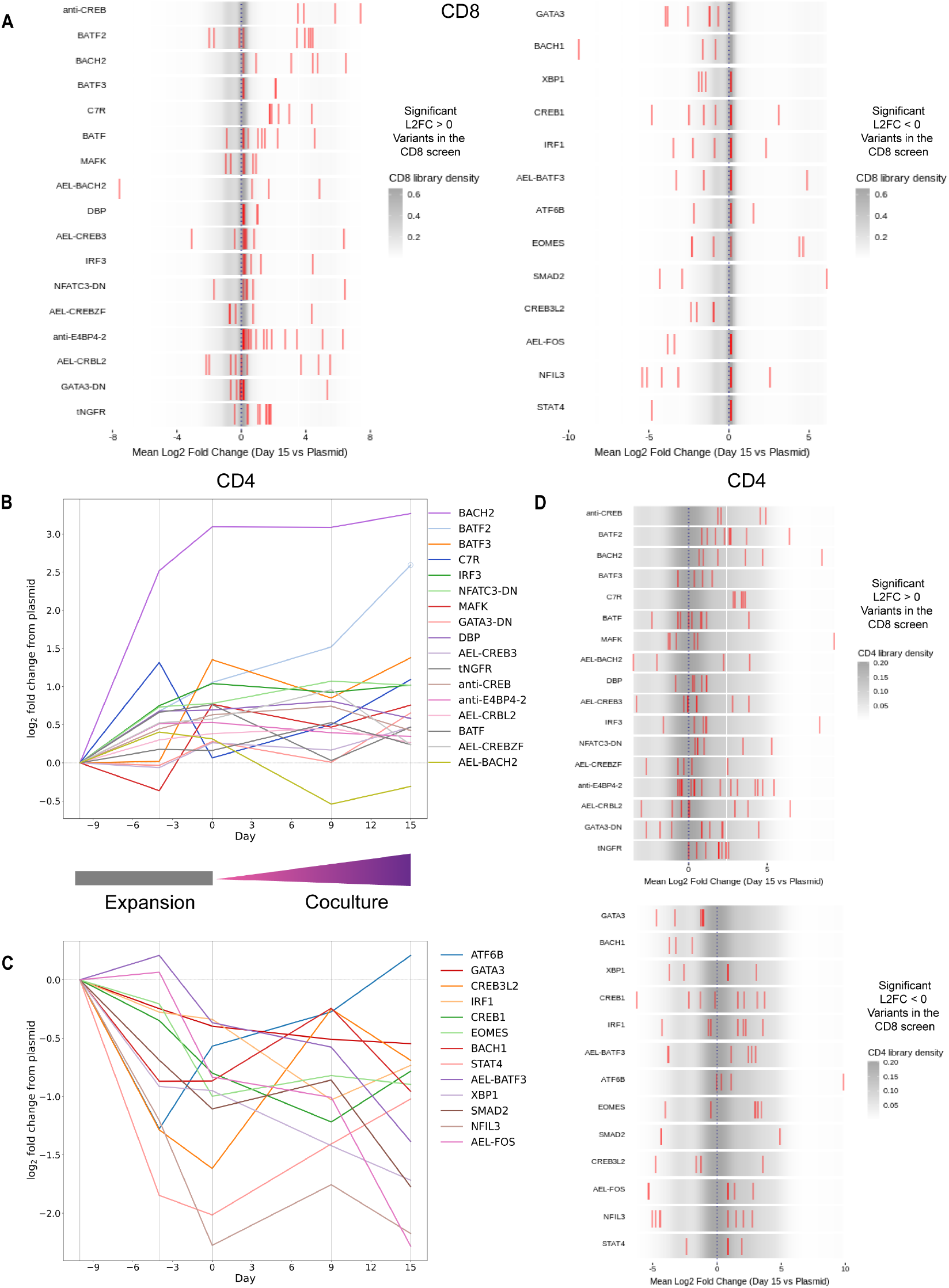
CD4^+^ CAR-T cell bZIP expansion/persistence screen results mirror CD8^+^ results. **(A)** Barcode level CD8^+^ log_2_ fold change (Day 15 vs. Plasmid) values (red bars) for variants significantly enriched or de-enriched in the CD8^+^ screen. The grey gradient in the background shows the distribution of all barcodes in the CD8^+^ screen. **(B)** and **(C)** CD4^+^ log_2_ fold change relative to the frequency in the plasmid pool (Day −10) vs. time for bZIP variants significantly enriched **(B)** or de-enriched **(C)** in CD8^+^ T cells. Log_2_ fold change fit by n = 2 donors, 2 replicates and calculated via DESeq2; significance calculated via Wald test in DESeq2. Open circles indicate timepoints with p_adj_ < 0.05. **(D)** Barcode level CD4^+^ log_2_ fold change (Day 15 vs. Plasmid) values (red bars) for variants significantly enriched or de-enriched in the CD8^+^ screen. The grey gradient in the background shows the distribution of all barcodes in the CD4^+^ screen. The results from one bZIP screen replicate on two donors are shown in (A) and (D).

**Figure S5.**
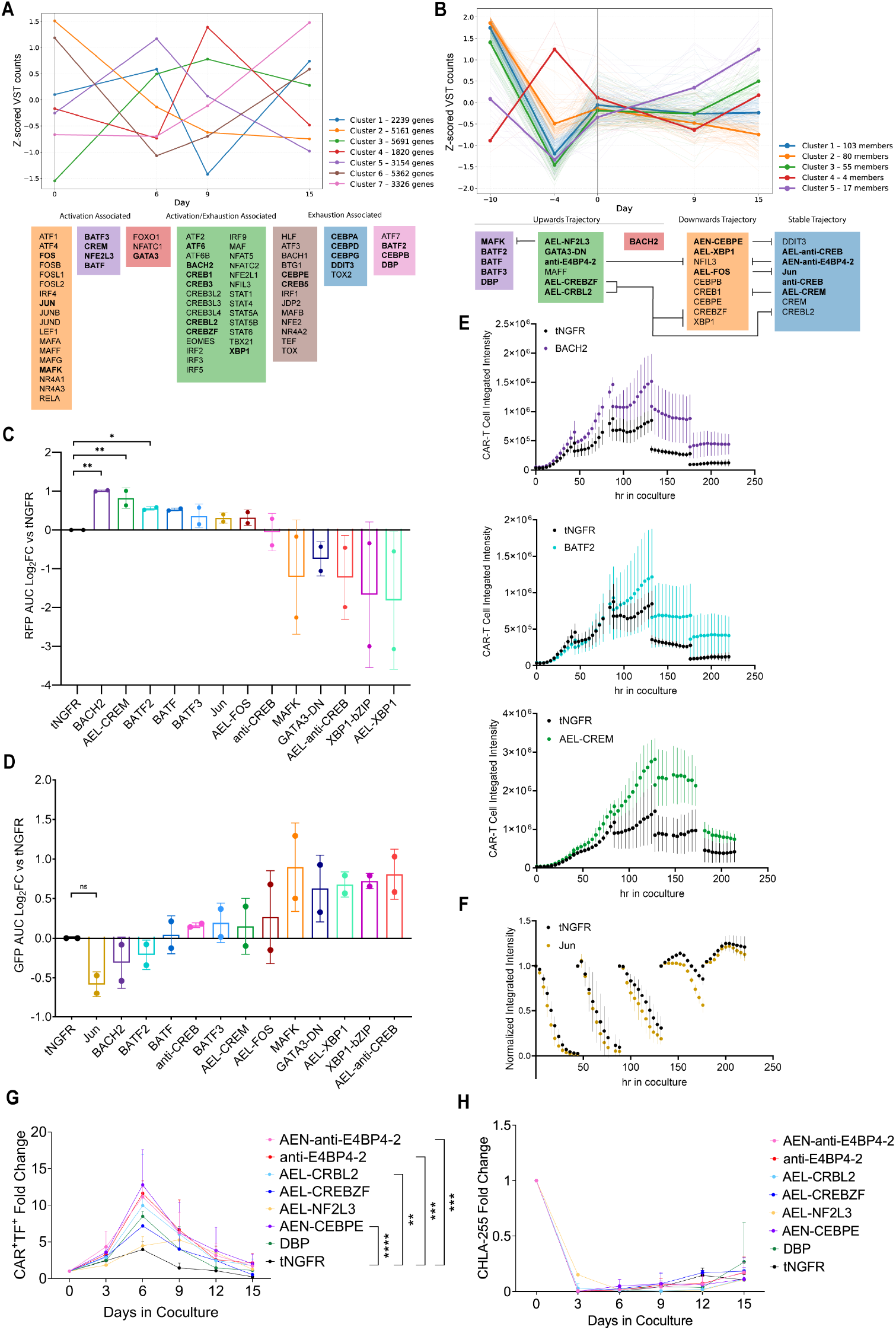
Proliferation and cytotoxicity validation of library hits. **(A)** mFuzz (*104*) soft clustering analysis of RNA sequencing VST counts GD2 CAR T cells in coculture, z-score normalized by shape of growth trajectory. Library variants belonging to each gene cluster are shown in colored boxes, with the putative targets of further tested TFs in bold. **(B)** mFuzz soft clustering analysis of bZIP library screen VST counts, z-score normalized by shape of growth trajectory. TFs that were further tested are in bold. **(C)** mKate2 RFP signal (log_2_ fold-change of area under the curve relative to CAR.tNGFR condition) corresponding to CAR-T cell expansion; n = 2 donors; error bars show SD; significance calculated by one-way ANOVA with Dunnett’s test for multiple comparisons; ** p < .01 * p < .05. **(D)** GFP signal (log_2_ fold-change of area under the curve relative to CAR.tNGFR condition) showing presence of tumor over time; n = 2 donors; error bars show SD; significance calculated via Welch’s test for JUN condition. **(E)** RFP integrated intensity (corresponding to CAR-T expansion) vs. time for best performing TF conditions in one donor. n = 4 technical replicates; error bars show SD **(F)** GFP integrated intensity (corresponding to tumor cell presence) normalized to the starting signal for each round of tumor stimulation for one CAR.JUN condition donor. n = 4 technical replicates; error bars show SD. **(G)** and **(H)** Fold change expansion **(G)** and tumor killing **(H)** of TF-inhibitor expressing CAR T cells and CHLA-255.GFP tumor cells. n=3 donors; error bars show SD; significance calculated by a two-way ANOVA with Dunnett’s test for multiple comparisons; ** = p < .01, *** = p < .001, **** = p < .0001.

**Figure S6.**
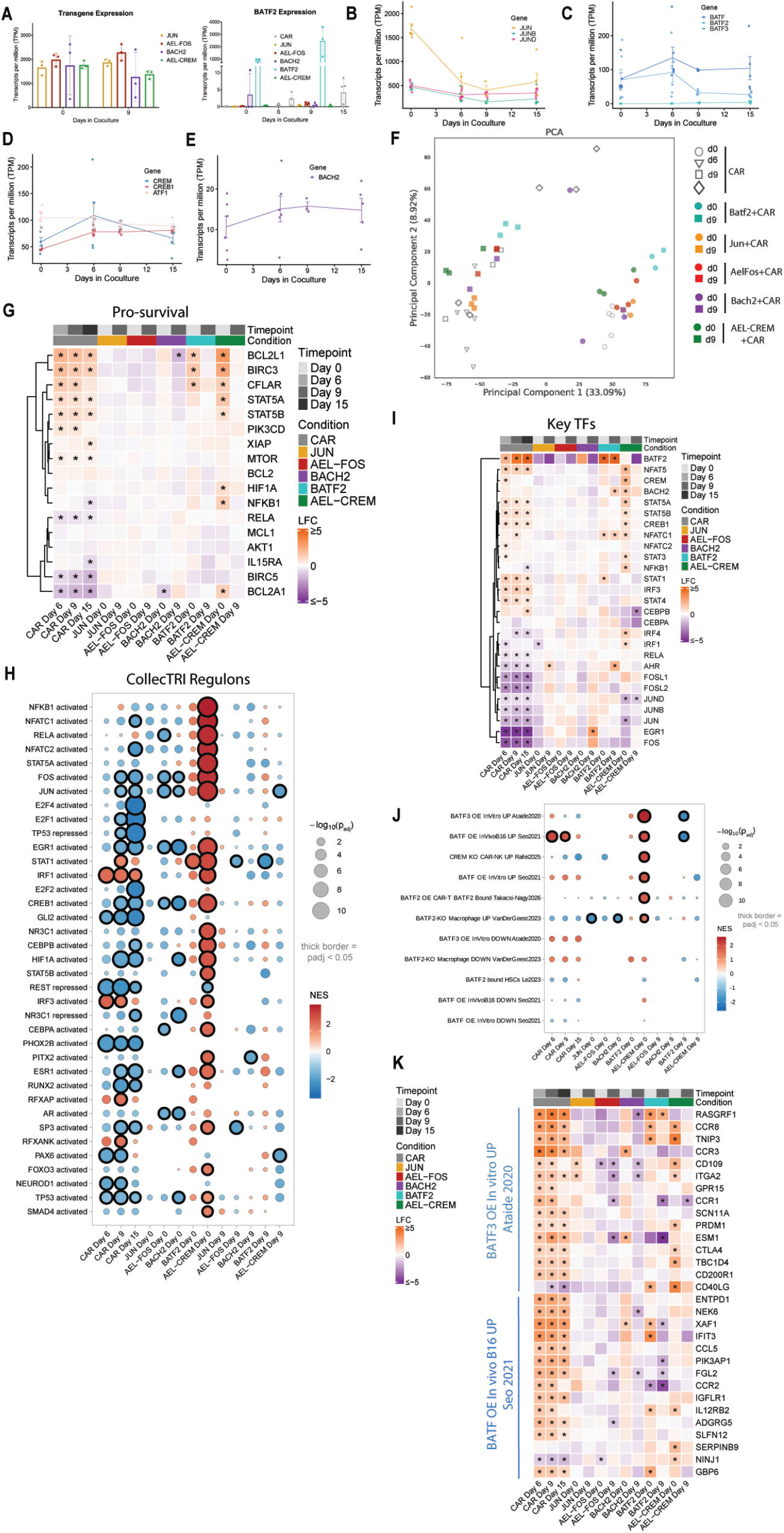
RNA-seq of CAR T cells. **(A)** Expression of TF transgenes (transcripts per million) for codon-optimized full-length bZIPs and inhibitors (left) and expression of BATF2 for all conditions (right). The wild-type BATF2 sequence was overexpressed for the GD2.BATF2 condition. **(B-E)** Expression of Jun **(B),** BATF **(C)**, and CREM **(D)** subfamily members and BACH2 **(E)** in GD2 CAR T cells undergoing chronic tumor exposure **(F)** PCA plot of TF-overexpressing CD8+ CAR T cell transcriptomes overlaid on CAR-only T cells. **(G)** Heatmap of log_2_ fold change (LFC) values for gene expression changes (CAR day 6-15 vs. CAR day 0, TF day 0 vs. CAR day 0, or TF day 9 vs. CAR day 9) for survival genes. **(H)** Gene set enrichment analysis of CollecTRI TF regulons on contrasts: TF day 0 vs. CAR day 0, TF day 9 vs. CAR day 9, or CAR day 9 vs CAR day 0. **(I)** Heatmaps of log_2_ fold change (LFC) values for gene expression changes (CAR day 6-15 vs. CAR day 0, TF day 0 vs. CAR day 0, or TF day 9 vs. CAR day 9) for key TFs **(J)** GSEA of the differentially expressed genes for the contrasts to BATF-family or CREM KO gene signatures (*15*, *18*, *31*, *85*, *86*, *102*). **(K)** Heatmap of LFC values for the leading-edge genes for CAR day 9 or BATF2 day 0 from GSEA comparison to the indicated BATF3 and BATF overexpression (OE) datasets. (E,F,H, *p_adj_<0.05 from DESeq2 Wald test with FDR correction).

**Figure S7.**
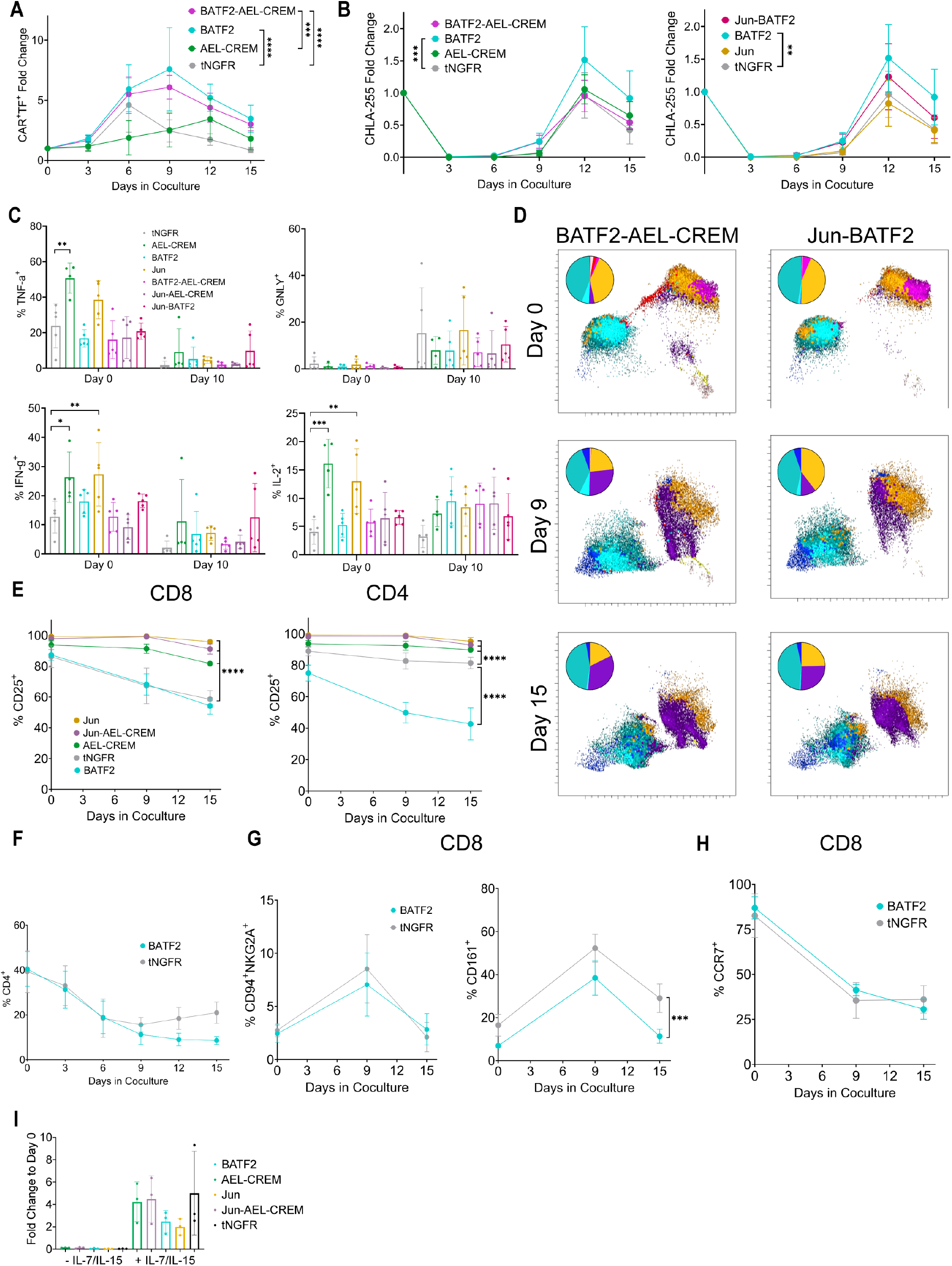
BATF2 Overexpression Improves GD2 CAR-T Cell Expansion But Impairs Activation and Persistence of CD4^+^ T cells. **(A)** Fold-change expansion of TF-overexpressing CAR T cells; n = 4-5 donors; error bars show SD; significance calculated via two-way ANOVA with Tukey’s test for multiple comparisons; **** p < 0.0001, *** p < 0.001 **(B)** Fold-change residual tumor relative to the tumor cells added for each round of stimulation; n = 5 donors; error bars show SD **(C)** Frequency of TNF-α^+^, IFN-γ ^+^, GNLY^+^, and IL-2^+^ TF-overexpressing GD2 CAR T cells at baseline and after 3 rounds of chronic tumor cell stimulation; n = 4-5 donors; error bars show SD; significance calculated via one-way ANOVA with Dunnett’s test for multiple comparisons; **** p < 0.0001, *** p < 0.001, ** p < 0.01, * p < 0.05. **(D)** UMAPs for paired BATF2 conditions per timepoint. Pie charts display the frequency of each FLOWSOM cluster. Cluster colors are the same as in Fig. 3. **(E)** Frequency of CD25^+^ cells for CD8+ (left) and CD4+ (right) GD2 CAR T cells in coculture; n = 5 donors; error bars show SD; significance calculated via two-way ANOVA with Dunnett’s multiple comparison’s test; **** p < 0.0001, *** p < 0.001. **(F)** Overall frequency of CD4^+^ BATF2-overexpressing GD2 CAR T cells in coculture **(G)** Expression of select markers in BATF2-overexpressing and CAR.tNGFR T cells in coculture; n = 5 donors; error bars show SD; significance calculated via mixed-effects model for the effect of condition; *** p < 0.001. **(H)** CCR7^+^ frequency in CD8^+^ CAR.BATF2 and CAR.tNGFR T cells **(I)** Fold-change expansion of TF-overexpressing GD2 CAR T cells cultured with or without IL-7 and IL-15 for 7 days

**Figure S8.**
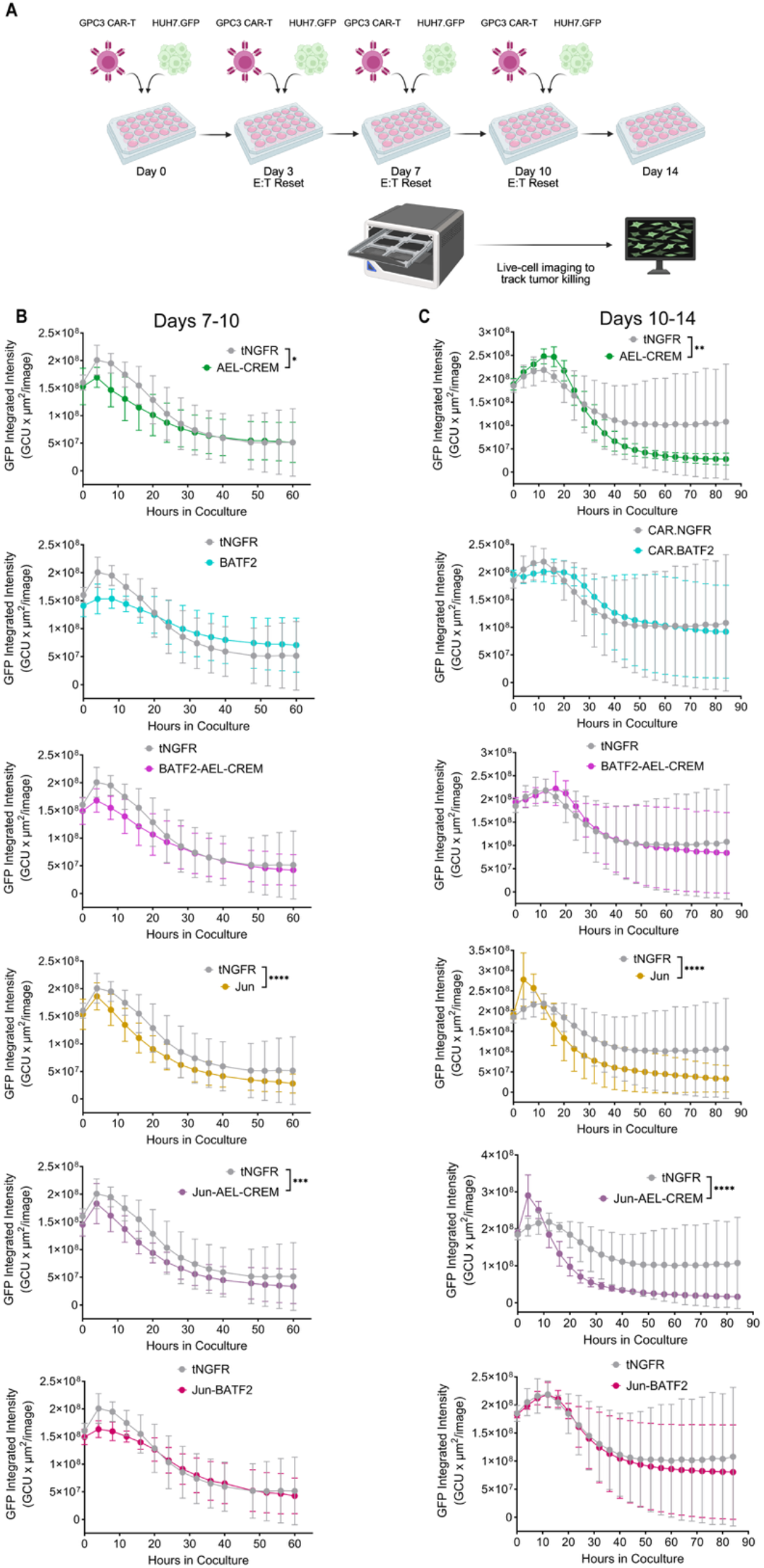
JUN, AEL-CREM, BATF2, and pairs show similar effects on cytotoxicity in GPC3.BBz CAR T cells. **(A)** Schematic of GPC3 CAR.BBz repeat HUH7.GFP tumor cell stimulation assay, with E:T reset to 1:1 every 3-4 days. **(B)** and **(C)** HUH7.GFP tumor cell killing for Days 7-10 **(B)** or 10-14 **(C)** of coculture, tracked by GFP via Incucyte S3 live-cell analysis system. n = 3-5 donors; error bars show SD; significance calculated via two-way ANOVA with Tukey’s multiple comparisons test for paired and constituent conditions plus control condition; **** p < 0.0001, *** p < 0.001, ** p < .01, * p < 0.05; (A) created in BioRender. Foight, G. (2027) https://BioRender.com/9jbphin.

**Figure S9.**
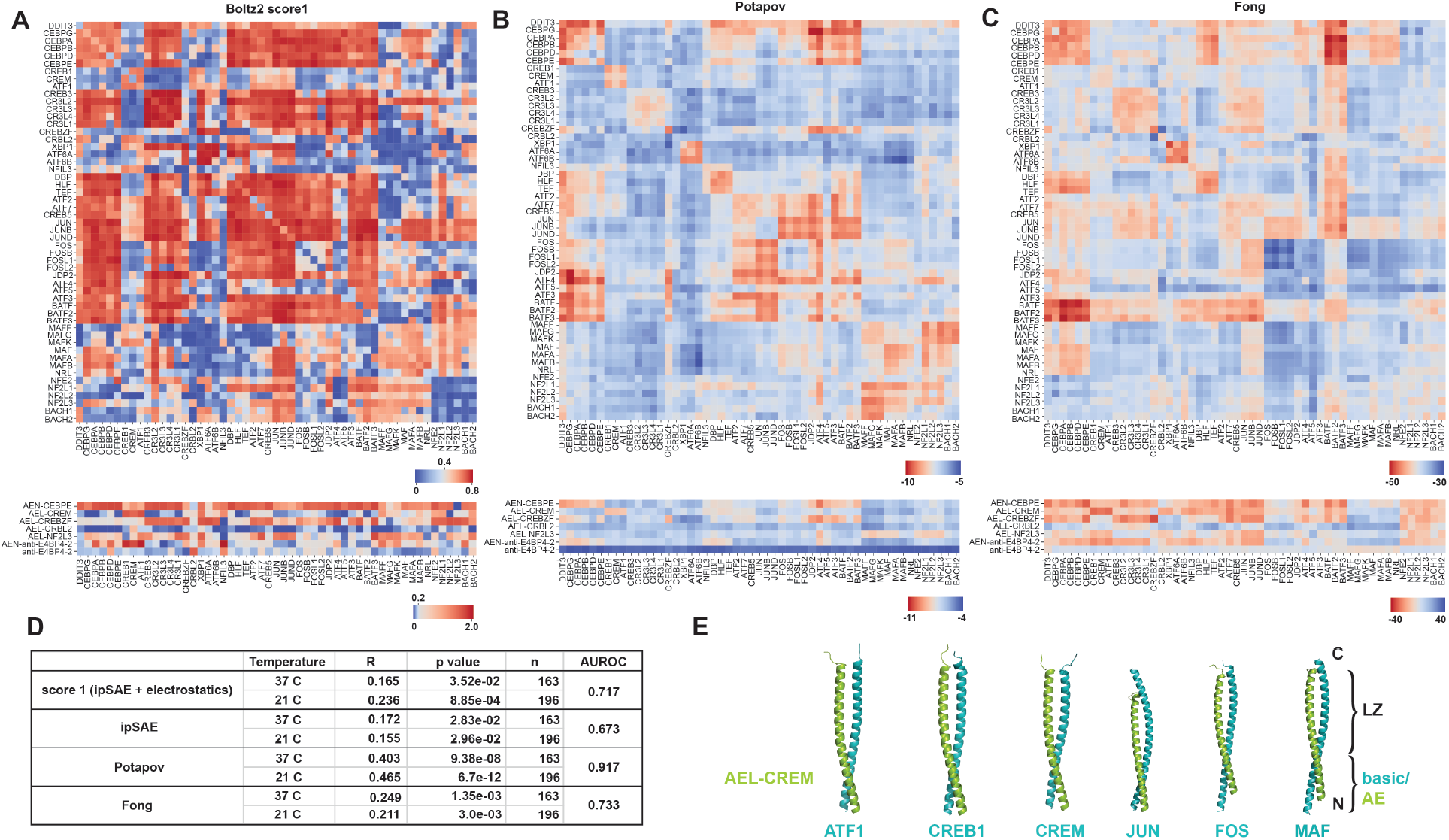
Modeling of bZIP and inhibitor complexes. **(A-C)** Heatmaps of model predictions from **(A)** Boltz-2 score 1, a composite score incorporating the ipSAE score of Boltz-2 predictions and electrostatic complementarity, and support vector machine models from **(B)** Potapov et al. (*34*) and **(C)** Fong et al. (*58*) for human bZIP pairs (top) and bZIP inhibitor library hits (bottom). For all scores, red is a prediction of higher affinity, blue of lower affinity interactions. **(D)** Performance of each model was evaluated against experimental datasets from (*26*) for human bZIP pairs, which comprised K_D_ values measured at two temperatures, 37 °C and 21 °C (n=number of experimental measurements scored). Pearson’s correlation coefficient (R) was calculated using continuous experimental binding measurements (K_D_). AUROC was calculated using binary binder/non-binder classifications, with binders defined as K_d_ ≤ 500 nM at two or more temperatures and non-binders defined as K_D_ > 5000 nM at all temperatures. **(E)** Representative Boltz-2-predicted structures of AEL-CREM (green) paired with selected human bZIPs (teal).

**Figure S10.**
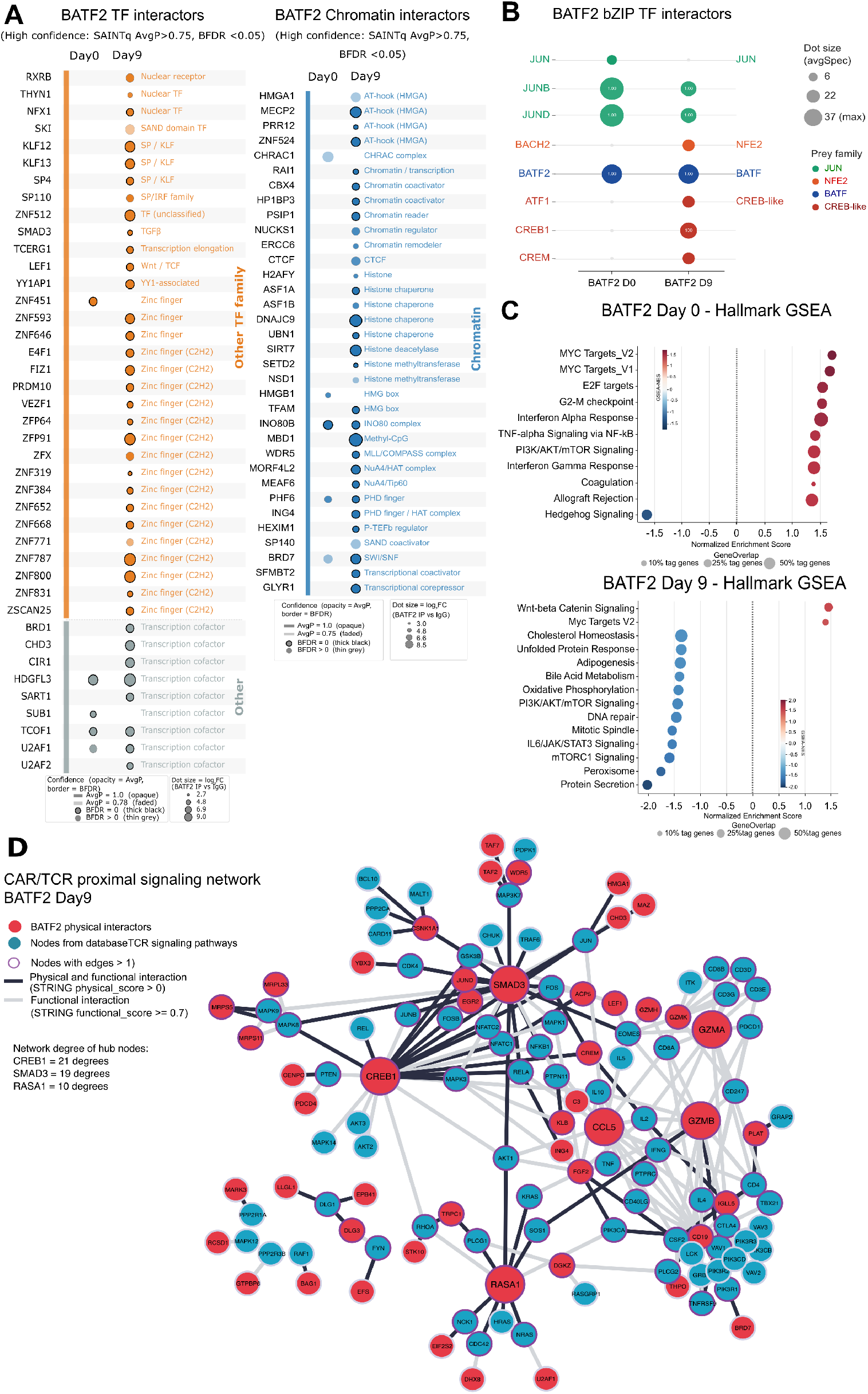
BATF2 interactor analysis. **(A)** Extended dot plot of high-confidence BATF2 TF and chromatin interactors continued from Fig. 4F (SAINTq BFDR<0.05). **(B)** Moderate-confidence BATF2 bZIP interactors (BFDR≤0.05, SAINTScore ≥0.70 with numeric SAINTScore labels on dots for AvgSpec>10 (AvgSpec, averaged spectral counts across replicates)). **(C)** GSEA Hallmark pathway enrichment analysis comparing the BATF2 interactome between resting (Day 0) and chronic stimulation (Day 9) conditions. (**D)** High-confidence BATF2 signalosome network at day 9, showing ultra-high-confidence BATF2 interactors (SAINTq AvgP = 1.0, BFDR = 0, iBAQ log₂FC ≥ 4.5, padj < 0.05) overlaid onto reference CAR/TCR proximal signaling nodes, with edges restricted to STRING v12 functional-association score ≥ 0.7 (128 nodes, 205 edges). Node color indicates origin (red: BATF2 physical interactors, n=48; blue-teal: reference pathway nodes, n=64); node size scales with degree; hub nodes= nodes with high degrees or edge connections in the network; edge shading distinguishes functional-plus-physical (black, n=81); (gray, n=123) from functional-only STRING evidence. See also Supplemental_Text_S2.

**Figure S11.**
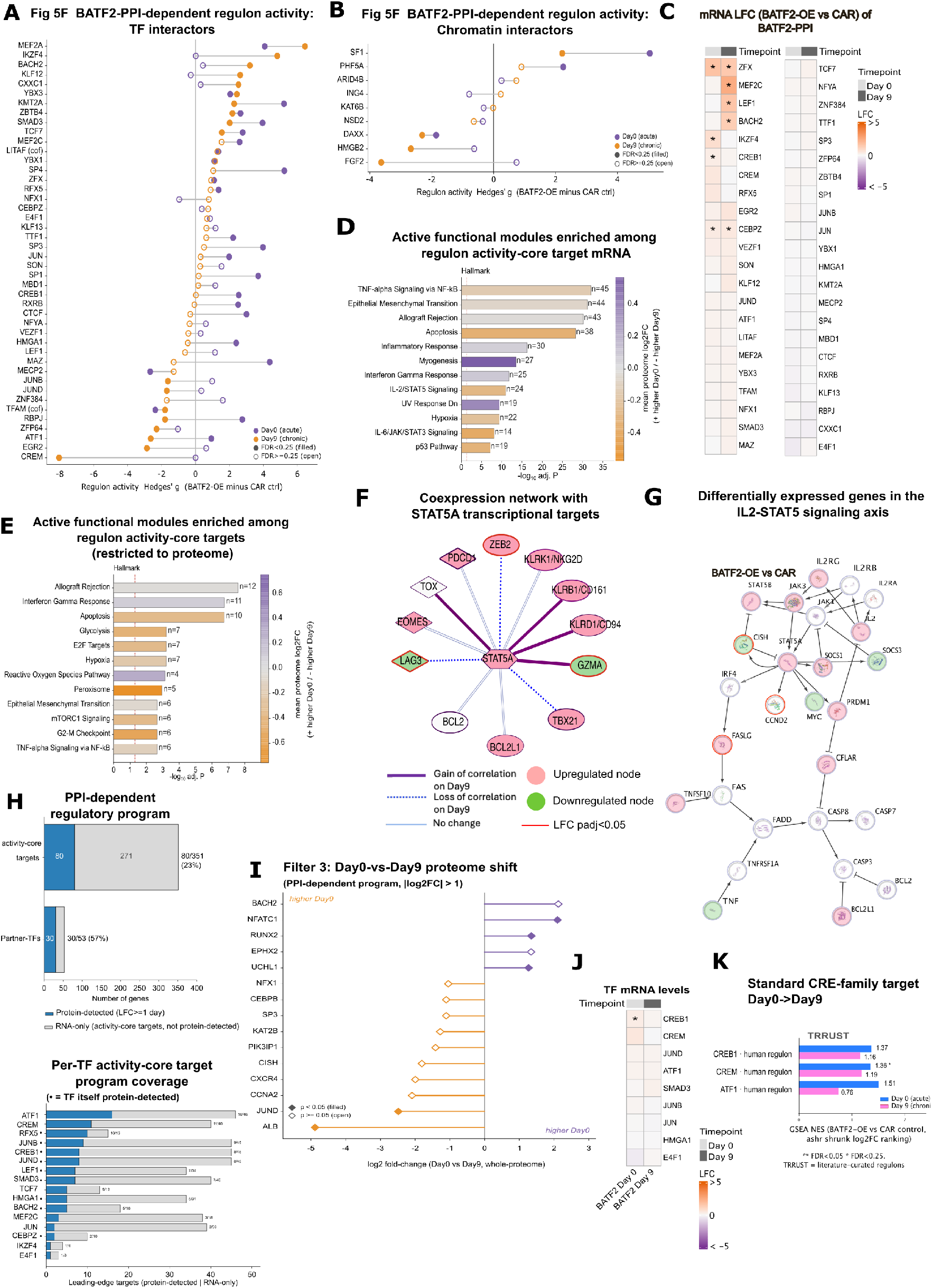
Active functional modules of the BATF2-PPI-dependent regulon. Significant BATF2 AP-MS partner-TFs **(A)** and chromatin modifying enzymes **(B)** with active regulons in the CollecTRI-ULM database, **(C)** their mRNA levels. **(D-E)** Active functional modules of the BATF2-PPI-dependent regulon. Hallmark (MSigDB Hallmark 2020) functional modules significantly over-represented among the activity-core targets of BATF2 AP-MS partner-TFs; TF CollecTRI regulon “activity-core” is defined as the high-contribution set of target genes that account for ≥90% of its net activity signal (ULM-weighted BATF2-vs-CAR effect); enrichment by Fisher’s exact test; Benjamini-Hochberg-adjusted p-values; n = number of activity-core genes overlapping the module. **(D)** Modules enriched among the mRNA activity-core target genes (bars colored by the mean RNA-seq LFC). **(E)** Modules enriched among the proteome-detected activity-core targets (bars colored by the mean whole-proteome LFC). **(F)** STAT5A coexpression networks in BATF2-OE Day9 GD2-CAR-T cells (Spearman rho Δ=day 9-day 0). **(G)** Flow diagram depicting IL-2-STAT5 signaling pathway activity in BATF2-OE GD2-CAR-T cells **(H)** Proteome-level coverage of the full BATF2-PPI-dependent regulatory program (top) and per-TF regulon (bottom, blue = genes detected in proteome, gray = genes detected only by RNA-seq). **(I)** Individual TF day 0 to day 9 shift at the proteome level. **(J)** mRNA levels of key TFs. **(K)** Orthogonal confirmation of CRE regulon activity trends on day 0 vs day 9 from gene lists obtained from TRRUST TF database (*100*).

**Figure S12.**
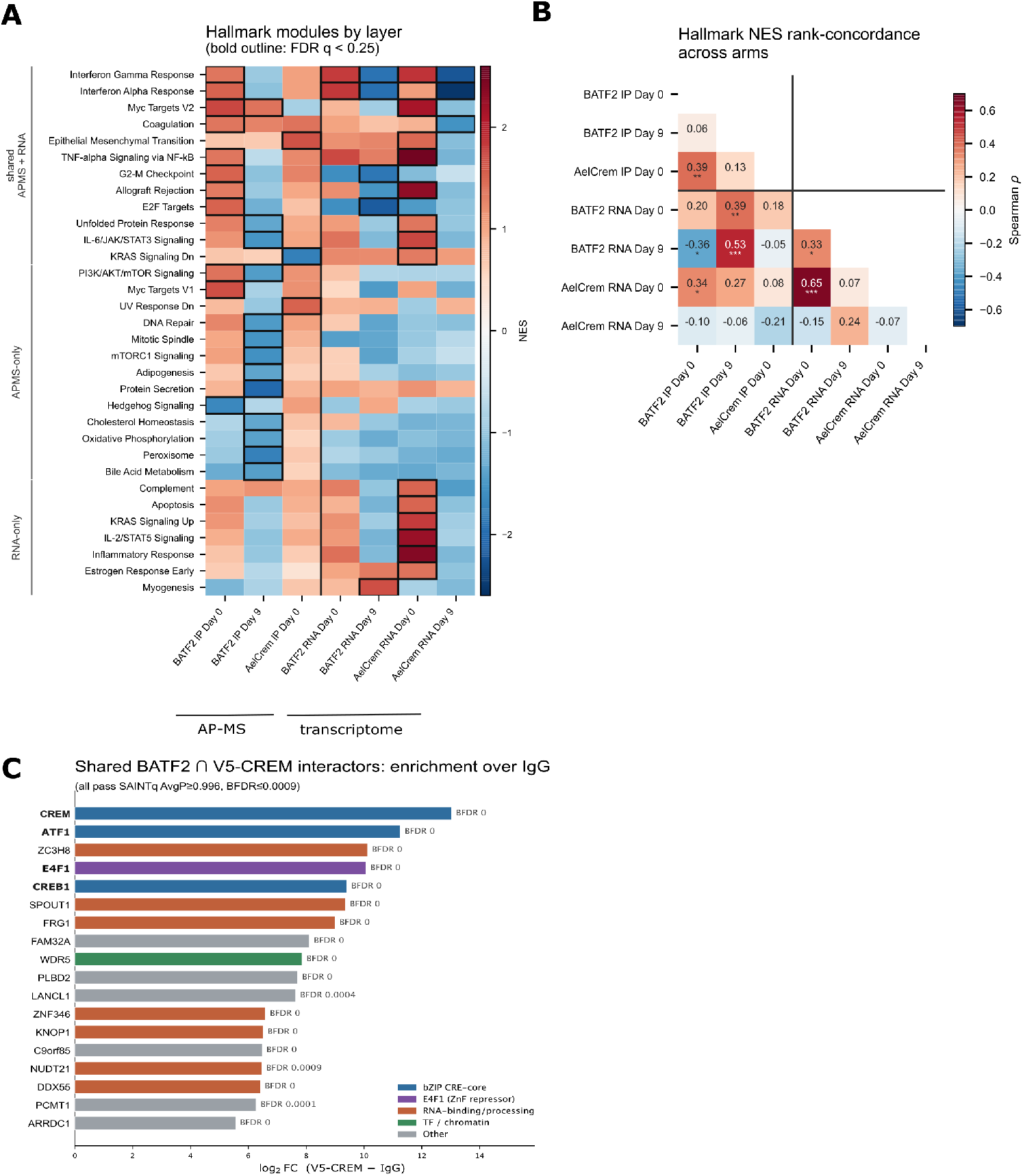
Comparison of BATF2 and AEL-CREM interactomes and transcriptomes in human GD2 CAR T cells. **(A)** Hallmark pathway analysis of AP-MS interactomes and transcriptomes colored by NES scores. Terms grouped by whether enrichment (q < 0.25) appears in both datasets, only the interactome, or only the transcriptome. **(B)** Heatmap of Hallmark NES concordance as measured by Spearman correlation (rho). **(C)** BATF2 and AEL-CREM AP-MS interactomes share 18-members notably CRE/CREB proteins. LFC values from AEL-CREM interactome (BATF2 LFC values in Fig. 5B).

**Figure S13.**
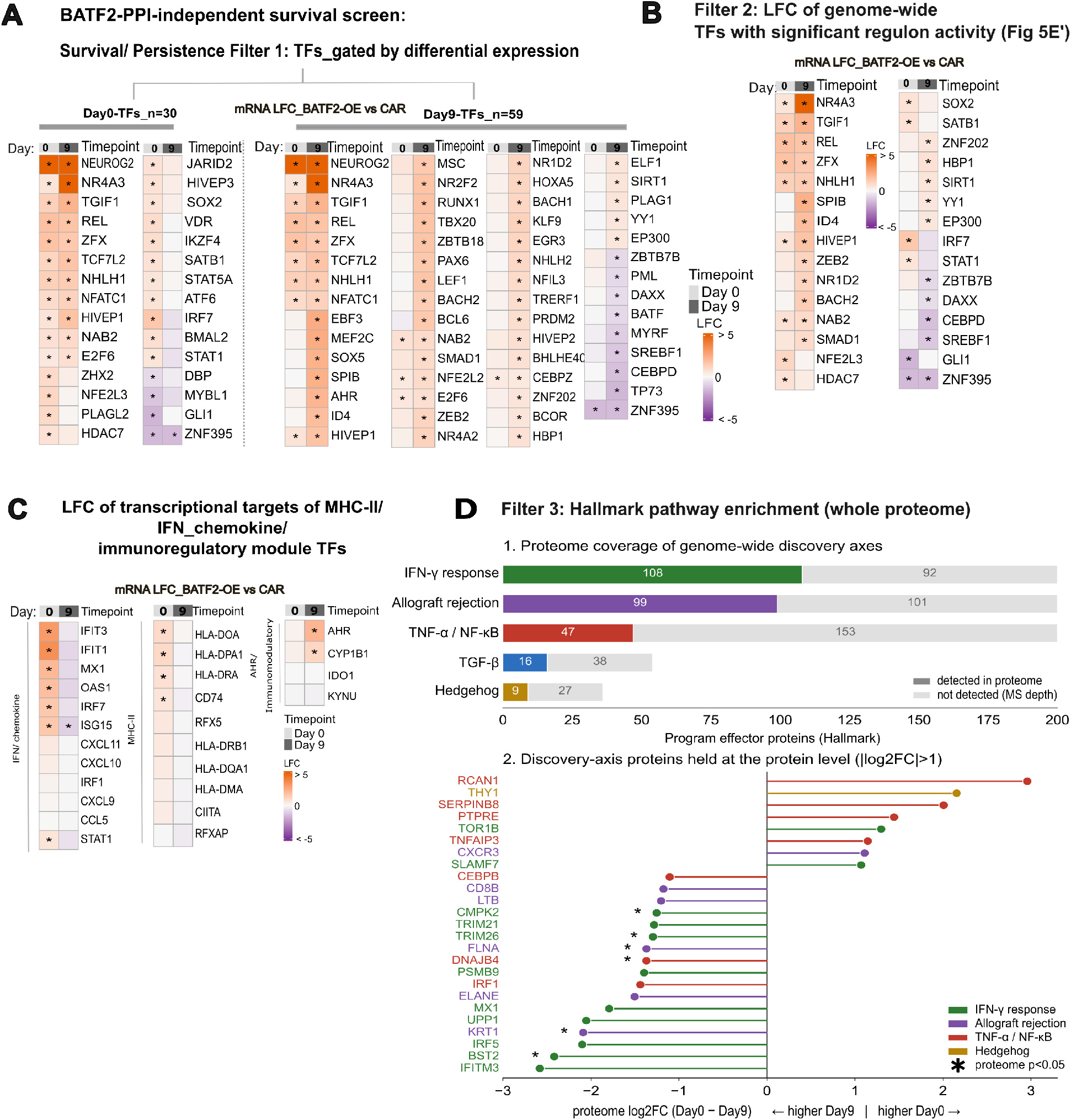
Filters used to define the survival/persistence programs in BATF2 CAR-T cells. **(A-B, D)**. Genome-wide analysis of active TF regulons **(A)**. A survival/persistence filter for differentially expressed TFs that do not associate with BATF2 (BATF2-PPI-independent, BATF2-OE vs CAR). TFs with significant regulon activity that are differentially expressed (|LFC| >0.5, BATF2-OE vs CAR, **B**). **(C)** BATF2-OE vs CAR: manually curated transcriptional targets of MHC-II, IFN_chemokine and AHR/immunoregulatory programs (LFC>0.05, padj<0.05.). **(D)** Enriched Hallmark pathway modules in BATF2 day 0 and day 9 whole proteomes (LFC>1.0, pvalue<0.05).

**Figure S14.**
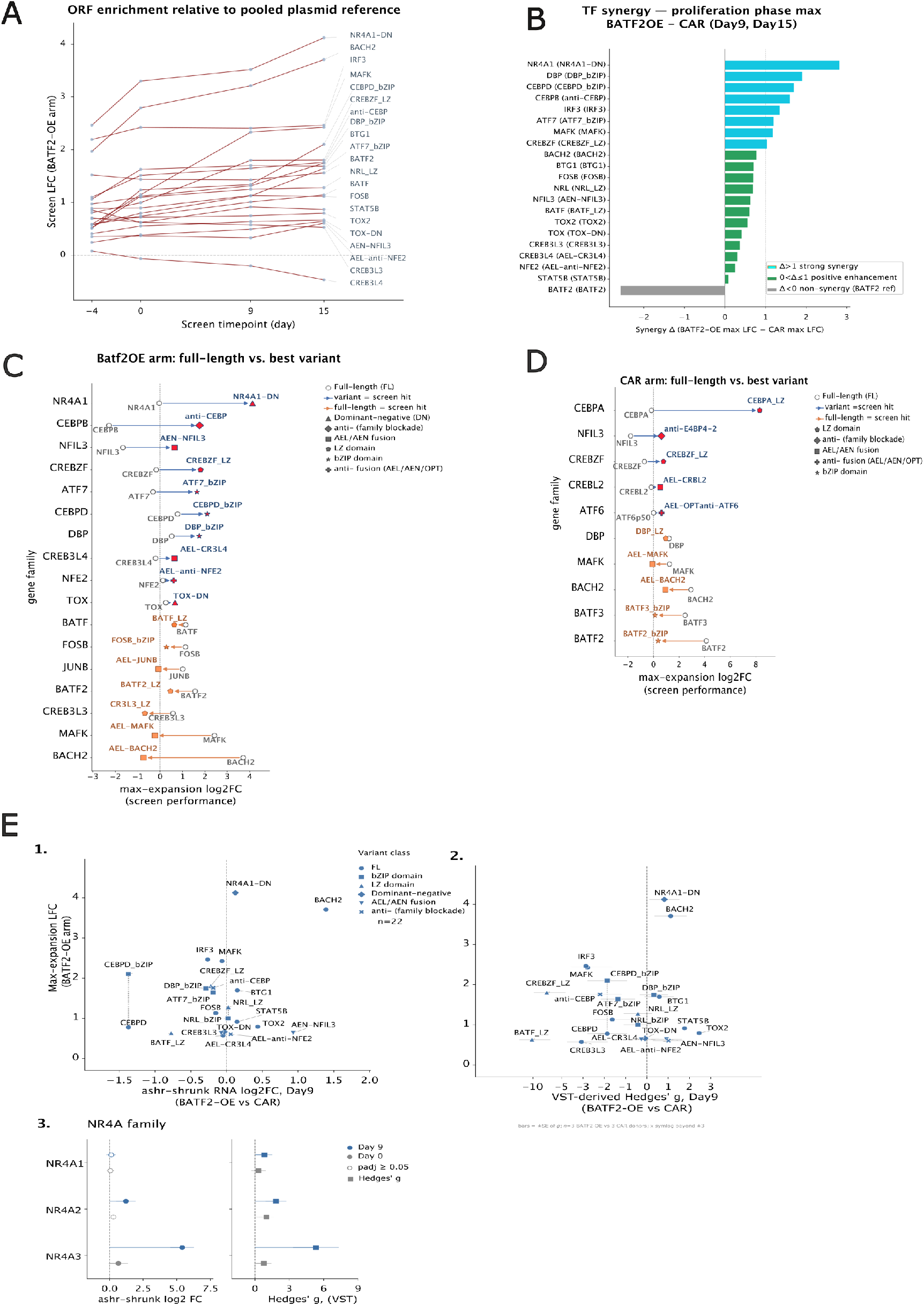
Comparison of pooled bZIP proliferation screens in BATF2 CAR and CAR-only backgrounds. **(A)** Enrichment relative to pooled plasmid library over time in the BATF2 screen for variants that show positive enhancement or synergistic increase in BATF2 CAR over the CAR-only background screen, graphed by **(B)** maximum proliferation (LFC ≥0.5) at day 9 or day 15 timepoints minus the LFC in the CAR-only screen (for trajectory p <0.05). **(C-D)** Enrichment and de-enrichment trends of full-length TFs vs their respective best-performing variants in the BATF2 CAR **(C)** and CAR-only **(D)** proliferation screen backgrounds. Each row represents a gene family. **(E)** Screen enhancement vs. mRNA abundance (1) and CollecTRI TF regulon activities (2) (day 9); (3) mRNA abundance and TF regulon activity of NR4A1/2/3 targeted by NR4A1-DN inhibitor. Horizontal bars = +/− SE of Hedges’ g; n=3 donors (BATF2 vs CAR); x-axis symlog beyond ±3.

**Figure S15.**
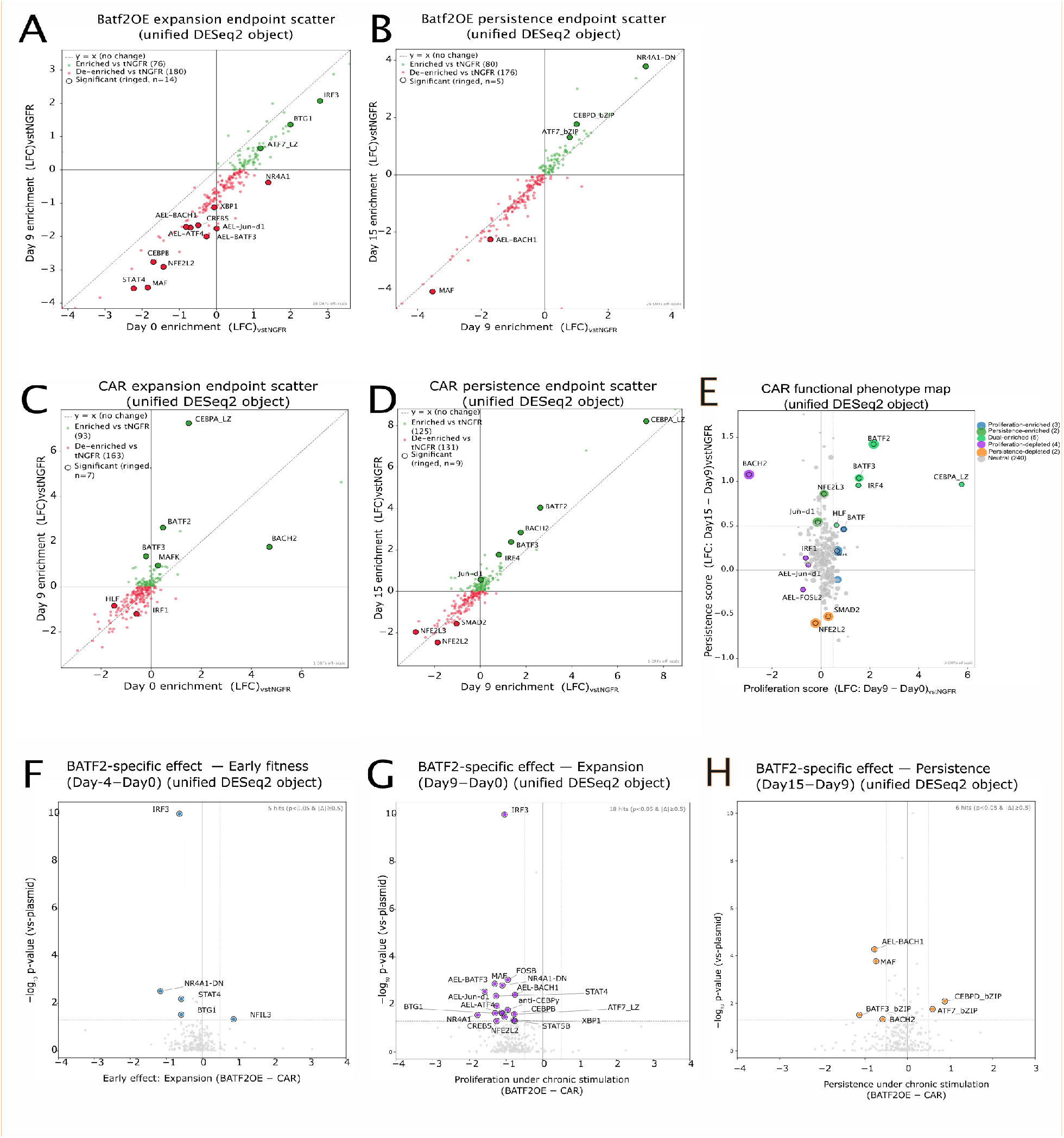
CAR/BATF2 screens proliferation/persistence phenotype phase diagrams. Enriched or de-enriched TF screen hits that enhance or counter proliferation in BATF2 CAR **(A-B)** or CAR-only **(C-E)** T-cells. **(F-H)** Volcano plots depicting hits that respond to BATF2 overexpression status (BATF2-specific effect). TF variant hits in the positive quadrant have enrichment higher than the CAR control and impact on early fitness, proliferation or persistence. Hits that confer early fitness during the expansion phase (enriched, F), proliferation (G, enriched) and late-stage persistence under chronic stimulation (enriched, H). (Significance obtained from unified DESeq2 object for timepoints vs. plasmid Wald raw p<0.05, and LFC ≥0.5). Supplementary information for Main Figure 5H.

**Figure S16.**
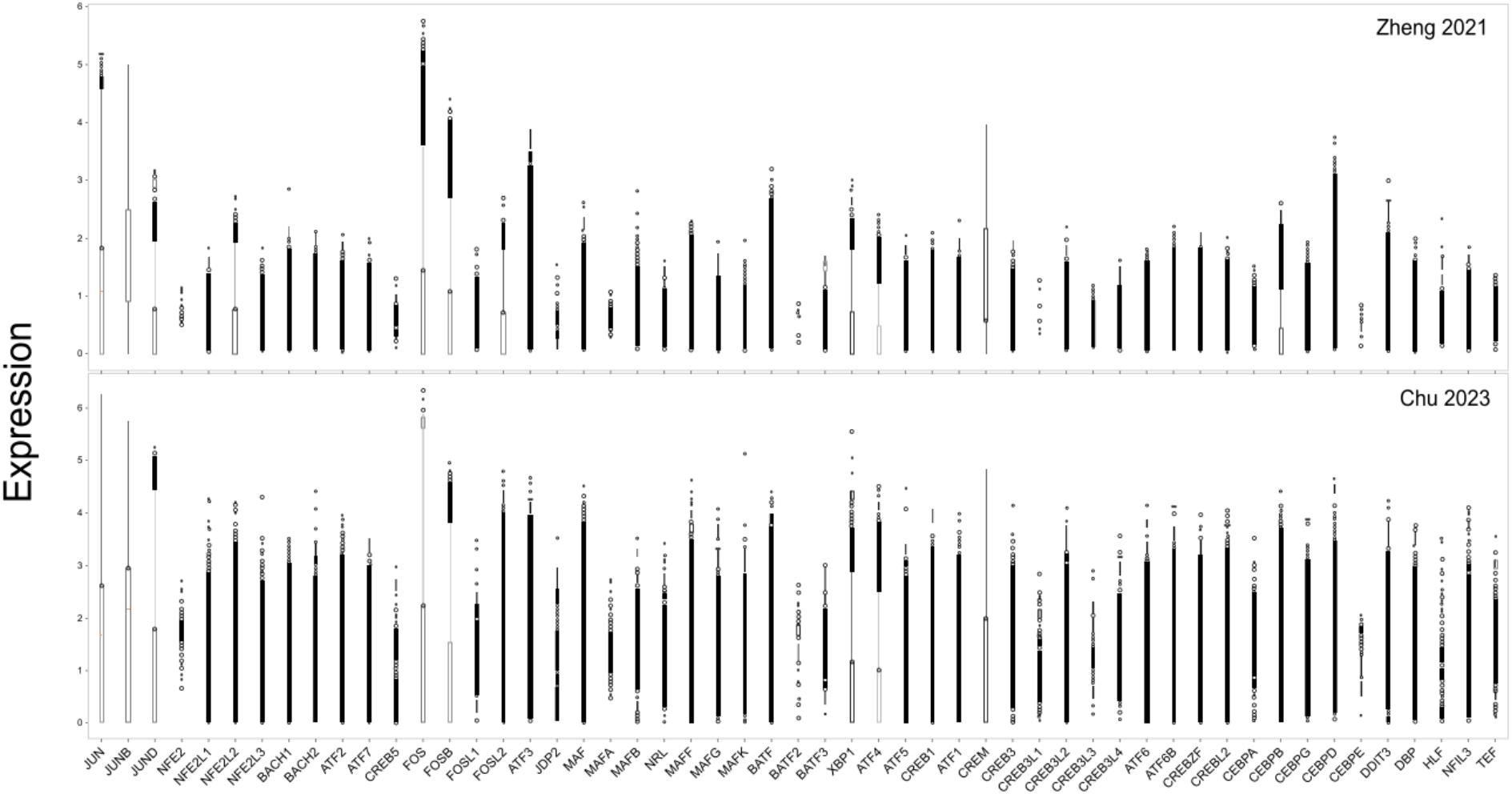
Endogenous bZIP transcription factor expression in human tumor-infiltrating T cells (hTILs). Distribution of log-normalized expression [log10(counts per 10,000)] of 53 human bZIP TFs across single T cells from two pan-cancer scRNAseq atlases: Zheng et al. 2021 (*91*) (n=108,884 cells) and Chu et al. 2023 (*92*) (n=110, 218 cells). Box plots show median and interquartile range. Endogenous BATF2 transcripts were detected in ∼0.02% of TILs (49 out of 219102 cells) indicating low expression rarely detectable by scRNA-seq.

**Figure S17.**
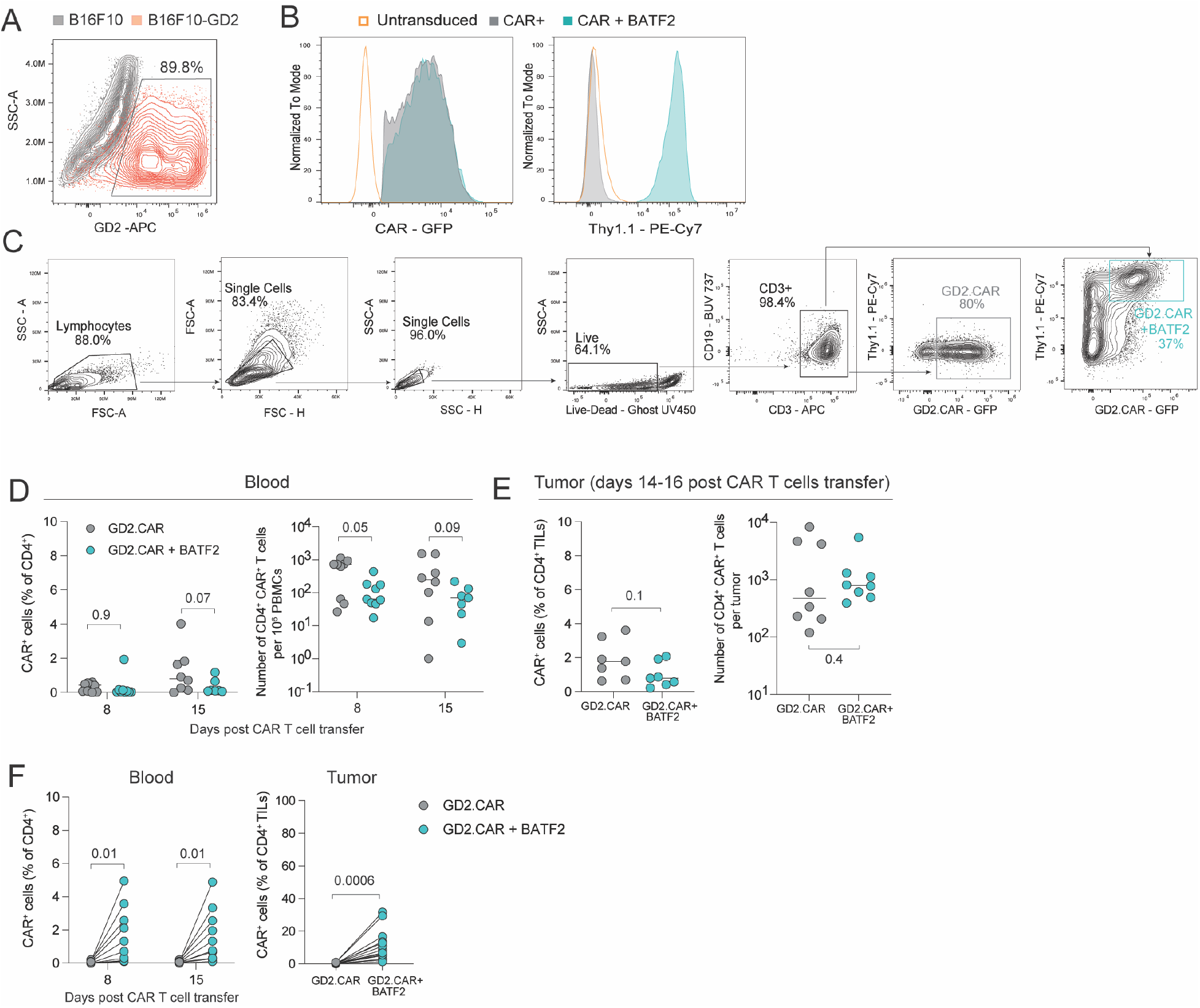
Generation of the murine GD2 CAR T cell model. **(A)** GD2 expression in wild type B16F10 (grey) and transgenic B16F10-GD2 (red) cell lines. **(B)** Relative expression of GFP (CAR) and Thy1.1 (BATF2) in the CD3^+^ T cells transduced with GD2 CAR or simultaneously transduced with GD2 CAR and BATF2. Gated on CAR^+^ T cells. **(C)** Gating strategy for FACS of CAR^+^ or CAR^+^BATF2^+^ T cells. **(D)** Relative frequency and numbers of CAR^+^CD4^+^ T cells in the blood at day 8 and 15 and **(E)** in the tumor at days 14-16 in the tumor control experiment. **(F)** Relative frequency of CAR^+^CD4^+^ T cells in the blood (left) at day 8 and 15 and in the tumor (right) at days 14-16 in the co-transfer experiment. Statistics are two-way ANOVA (D), ratio t-test on log-transformed data (F blood), paired t-test (F tumor), Welch’s t-test (E).

**Figure S18.**
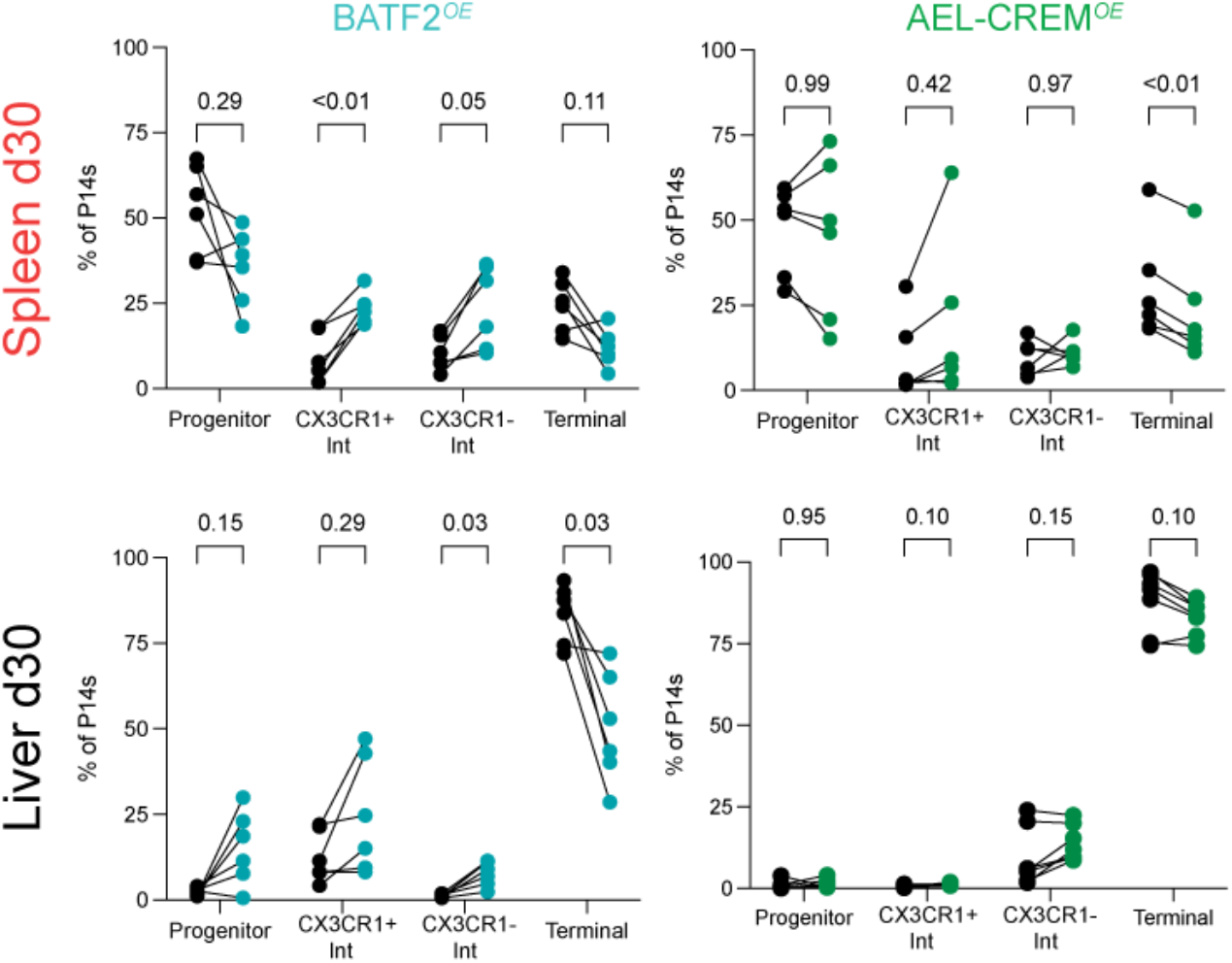
Exhaustion Subsets at Day 30 of P13 infection. Relative frequency of each subset within the indicated P14 population: untransduced (black), AEL-CREM (green), or BATF2 (turquoise) in the spleen and liver at day 30. Data are pooled from 2 experiments of n=2-5 mice per construct. All statistical comparisons were performed using matched 2-way ANOVA with Geisser-Greenhouse correction and Sidak’s multiple comparison’s test.

**Figure S19.**
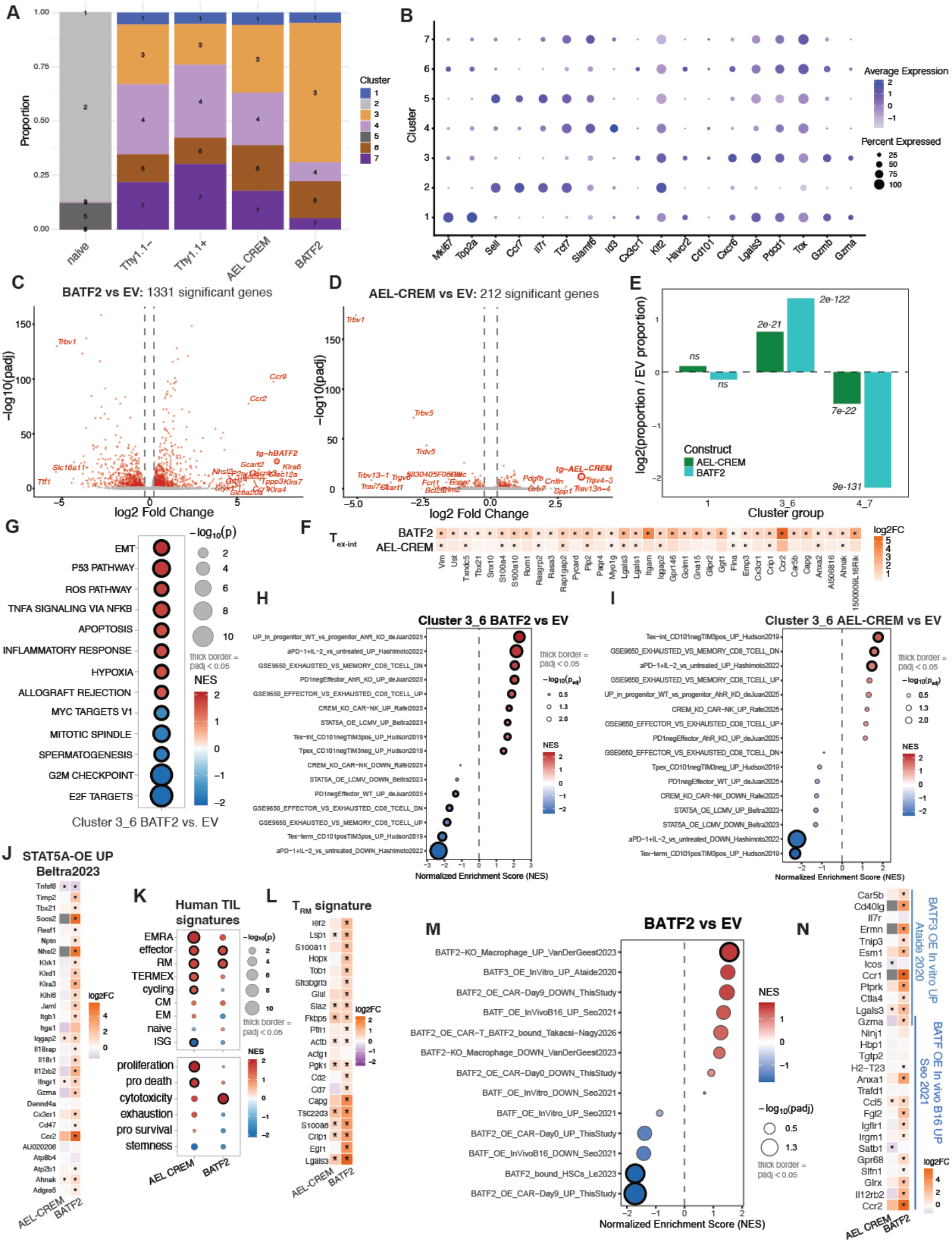
LCMV scRNA-seq analysis. **(A)** Cluster composition by sample. **(B)** Marker dot plot showing expression profile of clusters. Volcano plots of bulk differentially expressed genes (DEGs) for **(C)** AEL-CREM vs. empty vector (EV, Thy1.1+) and **(D)** BATF2 vs. EV (*padj<0.05, |log2fc|>0.25; transgenes denoted as tg-AEL-CREM and tg-hBATF2). **(E)** Enrichment of cells from AEL-CREM and BATF2 samples over the EV sample in cluster1 and grouped clusters 3_6 (Tex-int) and 4_7 (Tpex). P-values from Fisher exact test on cell counts (Supplementary Table STATs). **(F)** GSEA of DEGs in merged cluster 3_6 (Tex-int) for BATF2 vs. EV to MSigDB Hallmark pathways. **(G)** Tex-int state leading-edge genes from GSEA of the Tex-int Hudson2019 gene set to pseudobulk DEGs from BATF2 or AEL-CREM vs. EV. GSEA of DEGs in merged cluster 3_6 (Tex-int) for **(H)** BATF2 vs. EV or **(I)** AEL-CREM vs. EV to murine T cell signatures. **(J)** Heatmap of pseudobulk differential gene expression of BATF2 or AEL-CREM vs. EV for leading edge genes from the GSEA to the STAT5A OE LCMV UP Beltra 2023 gene set. **(K)** GSEA of pseudobulk AEL-CREM or BATF2 vs. EV to curated human tumor infiltrating lymphocyte (TIL) states and functional module gene lists (see Supplementary Table 3 for gene lists). **(L)** Heatmap of pseudobulk differential gene expression of BATF2 or AEL-CREM vs. EV for leading edge genes from the GSEA to the resident memory T cell (RM) TIL signature. **(M)** GSEA of pseudobulk BATF2 vs. EV DEGs to BATF/2/3 gene signatures (*15*, *18*, *31*, *85*, *86*). **(N)** Heatmap of pseudobulk differential gene expression of BATF2 or AEL-CREM vs. EV for leading edge genes of BATF2 GSEA to BATF and BATF3 signatures. (G, J, L, N, *padj<0.05).

## MATERIALS AND METHODS

### General cell culture

Peripheral blood mononuclear cells (PBMCs) were isolated from healthy donor buffy coats obtained from the Gulf Coast Regional Blood Center (Houston, TX, USA) by density gradient centrifugation using Lymphoprep™ (STEMCELL Technologies). T cells were cultured in RPMI containing 10% FBS, supplemented with GlutaMAX^TM^ (Thermo Fisher, cat. #35050061), and IL-7 and IL-15 each at 10 ng/mL. CAR-T cells in coculture did not receive added cytokines. HEK293T cells used for virus production were cultured in IMDM (Gibco™ Thermo Fisher, cat. # 12-440-053) containing 10% FBS and supplemented with GlutaMAX^TM^. CHLA-255 tumor cells were cultured in RPMI containing 10% FBS and supplemented with GlutaMAX^TM^. HUH7 and LN229 tumor cells were cultured in DMEM with 10% FBS.

### General construct design

For human T cell work, all constructs were cloned in an SFG gammaretroviral backbone. The second generation GD2.BBz CAR (hereafter, “GD2.CAR”) construct consisted of a 14g2a scFV, IgG1 hinge, CD8 transmembrane domain, 41-BB costimulatory domain, and CD3z endodomain. It was cloned as SFG-CAR-P2a-tCD19, and truncated CD19 (tCD19) was used to detect cells positive for CAR transduction. TFs were cloned as SFG-tNGFR-P2a-V5-TF, where truncated NGFR (tNGFR) is a transduction marker used to detect TF-positive cells, and V5 is an epitope tag for direct detection of TF expression. TF-pair constructs were cloned as SFG-tNGFR-P2a-V5-TF1-T2a-HA-TF2, where HA is an epitope tag for detection of TF2 expression.

For murine T cell work, a pMSCV retroviral backbone was used (MIGR1, a gift from Warren Pear (Addgene plasmid # 27490)). TFs were cloned as pMSCV-Thy1.1-P2a-V5-TF, or, for the construct coexpressing JUN and AEL-CREM, as pMSCV-Thy1.1-P2a-V5-cJUN-T2a-HA-AEL-CREM. A murine sequence-based equivalent of the human GD2.BBz CAR was designed as 14g2a, murine IgG2 hinge (UniProt P01863, residues 97-114), CD8 transmembrane (UniProt P01731, residues 193-222), 4-1BB (UniProt P20334, residues 212-256), CD3z (UniProt P24161, residues 52-164). It was expressed as pMSCV-CAR-P2a-mGreenLantern.

### Retrovirus production and transduction

Generation of human CAR T cells using gammaretrovirus was performed according to (*101*). Serial co-transduction of a GD2.BBz CAR followed 16-24 hr later by a TF variant was followed by expansion for 10 days before coculture.

For murine retroviral production, 293T cells were seeded in a 100mm dish (Corning) and incubated for 18–24 h until reaching ∼70% confluence, then co-transfected with pCL-Eco and a pMSCV-based retroviral transfer vector using GeneJuice transfection reagent according to the manufacturer’s protocol. Viral supernatant was collected 48 and 72 h post-transfection and filtered through a 0.45 μm PES filter. Supernatants were stored at 4°C in the dark between collections, then pooled, aliquoted, snap-frozen, and stored at −80°C until use. Frozen stocks were used to titer each batch of retrovirus prior to experimental transduction.

### bZIP library construction

bZIP/LZ domain annotations were based on (*28*) with slight modifications. bZIP domain constructs encompassed the 1^st^ **d** position of the bZIP domain (2^nd^ amino acid of the (*28*) annotation) to the **f** position at the end of the bZIP domain. LZ domain constructs encompassed the 2^nd^ **f** position before the LZ annotation in (*28*) to the **f** position at the end of the bZIP domain. For bZIPs that were not included in (*28*), we added one amino acid before and 2 amino acids after the Uniprot-annotated bZIP domain to match the format of the (*28*) annotation. We used this method because the bZIPs annotated in (*28*) all followed this pattern relative to the Uniprot annotations unless they had a shorter LZ domain.

AEN- and AEL-acidic extension peptide sequences are from (*37*) and (*38*), respectively. The AEN-version was used for AE variants targeting bZIPs containing asparagine, glutamine, aspartic acid, or glutamate (all polar amino acids) at the first **a** position N-terminal of the first **d** position of the LZ. The AEN-version contains asparagine at the corresponding amino acid to the **a** position in the target bZIP, thus creating a polar interaction. The AEN-version we used also included the E to R modification at the **g** position immediately N-terminal to a position **a** to create a more stable salt bridge. For AE-variants targeting bZIPs with polar lysine or arginine in the critical **a** position, the AEL-version (with leucine in the critical **a** position and the E to R replacement at the adjacent **g** position) was used because we inferred that the charged sidechain would interact with water distally from the inside of the coiled coil, while the aliphatic regions of the amino acid sidechains would form nonpolar interactions with leucine in the coiled coil. For all other AE variants, the AEL-version was employed to create a strong nonpolar interaction. The wild-type leucine zippers and anti-bZIPs fused to the AEs started at the second **g** position of the LZ domain, cutting off the first 8 N-terminal residues. Because many of the LZs fused to AEs could dimerize with multiple target bZIPs, we decided which AE version to fuse to the LZ based on the expected binding partners according to (*56*) and (*26*) [37 °C dataset]. In cases where expected binding partners had a mix of polar/nonpolar residues at the critical **a** position, we chose the AE version based on the number of polar vs nonpolar targets and the strongest binding partners. If binding partner data was not available for certain bZIPs from (*26*) or (*56*), we assigned the AE based on protein interactions from Uniprot or the protein interactions of a closely related bZIP. For a complete list of all peptide sequences in the library, see Supplementary Table 1.

The library was constructed as SFG-tNGFR-P2a-V5-TF-stop-barcode. For all non-full-length TF variants, a strong NLS (*102*) was added between the V5 and TF variant. Full-length TF sequences came from the DNASU repository, while subdomains and inhibitors were codon-optimized for human expression and ordered as gene fragments from IDT. See Table S1 for all peptide sequences. Each TF variant was first cloned into a Golden Gate donor vector with PaqCI sites and sequence confirmed. A receiver vector was constructed as SFG-tNGFR-P2a-V5-[PaqCI-bacterial_GFP-PaqCI]. Random 18-mer barcodes were ordered as a long oligonucleotide with flanking PaqCI sites and made double-stranded using Klenow polymerase. DNA of donor vectors encoding TFs were pooled at equimolar levels with an Echo Acoustic Liquid Handler. Pooled donor DNA, double-stranded barcode fragment, and receiver vector were mixed, and the Golden Gate reaction was performed with NEBridge^®^ Ligase Master Mix, PaqCI enzyme and activator (New England Biolabs), with 30 cycles of 37°C for 1 min and 16°C for 1 min, followed by an end soak at 60°C for 5 min. Reaction was cleaned and electroporated into NEB 10-beta electrocompetent cells, according to manufacturer’s recommendations. The library was bottlenecked to get an expected average of 10 barcodes per library member. The final pooled bZIP library plasmid was amplified with primers covering the TF-barcode region, randomly fragmented, and sequenced with a MiSeq 2×300bp protocol to map barcodes to bZIP library members. Sequencing identified an average of 40 barcodes per TF variant present in at least two copies in the sequencing data. Constructs encoding SFG-tNGFR-stop-barcode negative control and SFG-tNGFR-P2a-C7R-stop-barcode positive control (C7R, constitutive IL-7R) (*39*) were spiked into the final library.

### bZIP library screens and sequencing

Gammaretrovirus containing the bZIP library was titrated to achieve 20-25% transduction-positive cells, corresponding to a multiplicity of infection ∼0.2. The titrated amount of virus was utilized for the second transduction in the CAR-T generation protocol. NGFR^+^ cells containing bZIP library members were enriched using the EasySep™ Human CD271 Positive Selection Kit II (Stemcell, cat. #17849) 3 days after the second transduction. A coculture of GD2.CAR-T cells and CHLA-255.GFP neuroblastoma target cells was set up at an E:T ratio of 1:4, with the same number of target cells added once every 3 days for the 15-day coculture. Live CD8 and CD4 CAR(tCD19)^+^NGFR^+^ T cell pools were collected using a Sony SH800 on day −4 (day 6 of expansion), 0, 9, and 15 of the coculture. Genomic DNA was extracted with a Quick-DNA Microprep Kit (Zymo Resesarch, cat. #D3021). The barcode region was amplified in two PCR rounds, with the first round adding standard sequencing primer regions and the second round adding demultiplexing indices and p5/p7 adaptors. SYBR^TM^ Green I (Thermo Fisher Scientific) was used to track product and stop amplification in the exponential phase. The screen was performed in two donors, but with two independent runs starting from virus production, for a total of four biological replicates. Log2 fold change and p_adj_ values for each library member are in Table S2.

For the library screen in the background of BATF2 overexpression, a GD2.CAR-P2a-tCD19-T2a-HA-BATF2 construct was used in place of the usual GD2.CAR construct for the transduction protocol, and all other steps were completed as above. The BATF2 screen was completed concurrently in three donors.

### Library screen data analysis

Barcode abundance was quantified using a custom next-generation sequencing analysis pipeline. Raw paired-end FASTQ files were obtained from Azenta/GENEWIZ, reads merged using PEAR, and barcode-containing sequences were extracted using cutadapt based on invariant flanking sequences surrounding the barcode region. Exact-length 18-bp barcode sequences were retained, allowing one base pair mismatch. Barcode sequences differing by a single nucleotide were collapsed to their parent barcode to reduce sequencing and PCR artifacts. Barcode identities were mapped to their corresponding transcription factor constructs using the barcode lookup table generated during initial library validation, requiring a minimum of two independent observations per barcode for assignment.

Collapsed barcode counts were aggregated to the transcription factor level to generate sample-wise TF count matrices. Downstream statistical analysis on the TF level was performed separately for CD4 and CD8 T cell datasets using pyDESeq2 (version 0.4.11) (*103*), comparing TF abundances across timepoints relative to plasmid input controls or Day 0 samples. Filtering on low-count TFs was applied to reduce noise prior to normalization and dispersion fitting. Differential abundance statistics, including log2 fold-change (to plasmid or Day 0) and Wald test adjusted p-values, were generated from pyDESeq2 outputs and used for downstream visualization of transcription factor enrichment dynamics over time. For the initial bZIP library screen performed in the GD2.CAR background, data were fit considering the two donors and two replicates as batches. For the bZIP library screen performed in the GD2.CAR/BATF2 background, three donors were fit, and an overall gene dispersion model was fit including the CAR- and CAR/BATF2-background libraries to allow direct comparison of log2 fold-change values.

### Combined CAR/BATF2 and CAR-only bZIP library data analysis

TF-level counts were generated for the pooled BATF2-overexpression and CAR-T proliferation screens each sampled at Day-4, Day 0, Day 9, and Day15 timepoints of coculture with the CHLA-255 neuroblastoma cell line. Raw TF-level counts were combined from both screens and fit in a single pyDESeq2 object with a design combining screen (CAR/BATF2 or CAR-only) and timepoint, with the pooled plasmid library as the reference. ORFs with total counts across samples of ≥ 10 were included. Size factors and dispersions were jointly estimated across both screens to arrive at a common counts scale. For reporting within the BATF2-OE screen, the DESeq2 Wald test of the timepoint_vs_plasmid contrast for each screen_timepoint yielded log2FC with Wald standard error and p-val/padj (Benjamini-Hochberg adjusted) values. For cross-screen comparisons, tNGFR-referenced LFC for each TF variant was calculated post-hoc by subtracting log2FC(tNGFR vs plasmid) from log2FC (TF_ORF vs plasmid) within the same contrast to yield LFC_vs_tNGFR_ (TF_ORF enrichment relative to tNGFR).

TF variant enrichment and dynamics were further analyzed by three approaches: (1) Temporal dynamics of individual TF_ORF-vs-tNGFR contrasts from the two screens were assessed using Fisher’s Combined Probability test which yielded screen-specific trajectory p-values. (2) Maximum TF_ORF/variant effect across the full time-course was defined as the maximum LFC observed across Day−4, Day 0, Day 9, and Day15. (3) Phase-specific enrichment scores: Effect sizes (LFC) were used to identify TF variants with the strongest effect during the proliferation or persistence phase and to support downstream functional phenotype assignment (expansion= ΔLFC_vstNGFR_ (Day-4 - Day 0), proliferation= ΔLFC_vstNGFR_ (Day 9 - Day 0), and persistence = ΔLFC_vstNGFR_ (Day15 - Day 9). For functional phenotypic phase maps built from the proliferation screens, phase-specific enrichment scores within the BATF2-OE or CAR condition were deduced by ΔLFC_vs_tNGFR_ between consecutive timepoints and represented as scatter plots. A positive or negative score in the proliferation or persistence quadrants indicated pro-function or antagonizing effects, respectively. Additionally, BATF2-specific effect was derived by comparing phase enrichment scores between BATF2-OE and CAR screens, thereby distinguishing it from the baseline CAR program.

### IncuCyte proliferation and killing assay

GD2.BBz-P2a-mKate2 CAR-T cells were transduced with TF variants and expanded as described above. After expansion, CAR-T cells were cocultured with CHLA-255.GFP cells at an E:T ratio of 1:8 (25,000 CAR+:200,000 target cells, n = 4 technical replicates per donor condition). An additional 200,000 tumor cells were added every other day for a total of 5 rounds of stimulation (10 days). The Incucyte S3 was utilized to track CAR-T expansion and CHLA-255 killing via mKate2 and GFP integrated intensity, respectively. Area under the curve (AUC) analysis was performed for the mean integrated intensity and normalized to the CAR.NGFR control condition within each donor.

### Flow cytometry

#### CAR-T Proliferation and Tumor Killing Flow Beads Counting Assay

GD2.CAR-T cells were cocultured with CHLA-255.GFP target cells (E:T = 1:4) such that each condition per donor had 1 well for each timepoint. The same amount of additional target cells was added to each well every 3 days for 5 rounds of tumor exposure total (15-day coculture). For each timepoint, the entire well was harvested, stained, and mixed with CountBright^TM^ Absolute Counting Beads (Thermo Fisher Scientific, cat. # C36950) diluted 10x in PBS before acquisition on a Cytoflex LX.

#### Annexin V Flow Cytometry

CAR-T cells were taken from culture into flow cytometry tubes, washed, and stained for surface markers. Each tube was washed with 1 mL of Annexin V binding buffer (BD, cat. #556454) before AnnV PE stain (BD, cat. #556422, 3 uL per tube) was added. 7-AAD viability stain (BD, cat. #559925, 2 uL per tube) was added shortly before acquisition on a Cytoflex LX.

#### Proliferation Dye Flow Cytometry

GD2.CAR-T cells at a concentration of 2e6/mL were stained in pre-warmed PBS solution containing 1.25 µM CellTrace^TM^ Violet (Thermo Fisher, cat. #C34557) proliferation dye for 10 minutes at 37 °C. The staining was quenched with RPMI containing 10% FBS (5x the staining solution volume). After exchanging cRPMI media to add IL-7 and IL-15 (10 ng/mL), 2e6 T cells were plated per well on a non-TC-treated plate that had been coated overnight at 4 °C with 1 mL/well of 1A7 anti-idiotype (NCI) antibody solution (.25 µg/mL) to stimulate the CAR T cells. CTV-stained T cells were surface-stained and data were acquired on a Cytoflex LX at baseline and after 3 days of 1A7-idiotype stimulation.

#### Spectral flow immunophenotyping

GD2.CAR-T cells were cocultured with CHLA-255.WT target cells in a 15-day coculture, as described above for the flow bead counting assay. At baseline, day 9, and day 15 of coculture, CAR-T cells were collected for each condition and washed with FACS buffer (PBS containing 2% FBS and .002 M EDTA). Fc block (Biolegend, cat. #422302, 1 µL/tube) and Brilliant stain buffer (BD, cat. #659611) were added to each tube according to manufacturer protocols before surface marker staining and a FACS buffer wash. Fixation and permeabilization were performed using the eBioscience^TM^ Foxp3/Transcription Factor Staining Buffer Set (Thermo Fisher, cat. #00-5523-00). Mouse serum (Thermo Fisher, cat. #24-5544-94) was added to minimize non-specific binding before intracellular marker staining. After washing with permeabilization buffer and FACS buffer, data acquisition was performed on a Cytek Aurora. Downsampling to 5000 events (2500 each of CD4^+^ and CD8^+^ CAR^+^TF^+^ cells) for each sample was performed prior to concatenation and clustering analysis with UMAP (*104*) and FLOWSOM (*49*).

#### LCMV sample immunophenotyping

After leukocyte isolation, staining was performed by surface staining cells with indicated antibodies and Live/Dead Ghost UV450 Fixable Viability Dye (Tonbo) for 30 min at 4°C, followed by washing with FACS buffer. Intracellular staining was performed using the intracellular staining Foxp3/Transcription Factor Staining Buffer per manufacturer’s instructions. Samples were acquired using a 5-laser Cytek Aurora and analyzed using FlowJo, and summary graphs and statistics were prepared in GraphPad Prism v10.

#### Cytotoxicity Marker Coculture and Intracellular Staining

GD2.CAR T cells were analyzed for cytokine and effector molecules at baseline (day 0), or after 9 days of coculture with CHLA-255 (E:T = 1:2), followed by 1 day of rest in pure RPMI. Baseline or rested T cells were acutely stimulated for 5 hr at a 1:1 E:T with CHLA-255.GFP target cells with Brefelden A (GolgiPlug^TM^, BD, cat. #555029) and Monensin (GolgiStop^TM^, BD, cat. #554724) at manufacturer’s recommended concentrations. After acute stimulation, T cells were harvested and washed with FACS buffer before surface staining, fixation/permeabilization (Cytofix/Cytoperm^TM^ Fixation/Permeabilization Kit, BD, cat. #554714), and intracellular staining. Due to the presence of multiple Brilliant Violet fluorophores for intracellular markers, True-stain^TM^ Multifluor Buffer (Biolegend, cat. #426105) was added immediately prior to intracellular staining. Data were acquired on a Cytoflex LX.

### Bulk RNA sequencing experiments

#### Culturing and RNA extraction

GD2.CAR T cells with or without TF-P2a-tNGFR cotransduction were cocultured with CHLA-255.GFP cells at 1:4 E-T, with fresh CHLA-255 added every 3 days. For the day 0 timepoint, T-cells were exposed to CHLA-255.GFP cells in co-cultures for ∼3 hours prior to sorting. All other timepoints were harvested directly from coculture at the end of a 3-day coculture period. GD2.CAR-only samples were harvested at day 0, 6, 9, and 15, while GD2.CAR cotransduced with JUN, BATF2, BACH2, AEL-CREM, or AEL-FOS were harvested at days 0 and 9. CD8^+^ GD2.CAR T cells (live/T-cells(GFP^−^)/CAR^+^(tCD19^+^)/TF^+^(tNGFR^+^)/CD8^+^) using a SONY SH800S sorter. The number of isolated CD8^+^ T cells for each pool was ∼500,000 – 2,500,000. The cells were then lysed with RLT lysis buffer from the Qiagen RNEasy kit (cat# 74104), RNA extracted according to manufacturer’s instructions and sequenced with Azenta or Admera Health sequencing services using standard poly-A capture protocols.

#### RNA-seq data analysis

Raw FASTQ files were aligned to the human reference genome (GRCh38/hg38) using STAR (v2.7.10) (*105*) with default parameters with a genome index built from Gencode v44 primary assembly annotations (with custom CAR, tNGFR, and TF transgenes added), producing coordinate-sorted BAM files for downstream analysis. Raw count matrices were adjusted for batch effects using pyComBatSeq (*106*, *107*). Batch-corrected counts were subsequently processed using pyDESeq2 (v0.5.4) (*103*) to perform variance-stabilizing transformation (VST). Principal component analysis (PCA) was conducted on VST-transformed expression values including all expressed genes. Gene dispersions were jointly fit across all CAR-only and transcription factor (TF) datasets to account for shared sources of biological variability. Transgenes were excluded from the gene dispersion fitting step, as their extreme counts resulted in poor fits. The exception to this was BATF2, for which the endogenous sequence was used for the overexpression vector, disallowing separation of endogenous TF and transgene counts. Differential expression between TF and CAR-only conditions, or across timepoints, was subsequently estimated by computing log2 fold-changes (LFC) with Wald testing within this unified dispersion model, and adjusted p-values (padj) calculated using the Benjamini-Hochberg method. Heatmaps of LFCs reported raw LFCs and padj values, as shrunk LFCs were found to contain anomalies for some conditions with high donor variance (BACH2). For Figure 5 BATF2 bioinformatic and network analyses, ashr-shrunk LFC values from an earlier fit that excluded the AEL-CREM dataset were used.

#### Gene signature analysis

Gene set variation analysis (GSVA) (*108*) was performed in R using the GSVA Bioconductor package (v2.4.7) on variance-stabilized RNA-seq expression matrices of GD2.CAR T cells overexpressing JUN, BACH2, AEL-FOS, BATF2, or AEL-CREM to quantify enrichment of predefined T cell functional and cell-state transcriptional programs across samples. Gene signatures were curated from published human tumor-infiltrating lymphocyte (TIL) datasets (*91–93*) (“cell state signatures”) and custom functional modules (Table S3). To minimize redundancy between T cell state signatures from the TIL datasets, overlapping genes were resolved using weighted filtering that assigned genes uniquely to the most strongly supported cell state based on literature frequency and overlap metrics. GSVA scores were calculated using the dense classical algorithm with Gaussian kernel-based empirical cumulative distribution function estimation, excluding transgenes and genes with constant expression across samples. Statistical analysis was performed using linear mixed-effects models implemented in R (lmer), fitting the model score ∼ condition * timepoint + (1|donor), where condition and timepoint were treated as fixed effects and donor as a random effect (Table S4). P-values were adjusted for multiple testing using the Benjamini–Hochberg method, and results were visualized in radar plots of the mean across donors of z-score normalized GSVA scores.

Pre-ranked gene set enrichment analysis (GSEA) was performed using the fgsea R package (v1.36.2) (*46*, *109*, *110*). For each CAR T cell dataset contrast (TF Day 0 vs. CAR Day 0, TF Day 9 vs. CAR Day 9, CAR Day 6, 9, 05 15 vs. CAR Day 0), genes were ranked by raw pyDESeq2 LFC and restricted to genes with adjusted p-value < 0.05 to focus the GSEA analysis on pathways enriched amongst the most strongly regulated genes for each TF. For LCMV contrasts (BATF2 vs. EV, AEL-CREM vs. EV), DEGs were ranked by (avg_log2FC)*-log10(p_val). Gene sets for comparison were drawn from the MSigDB Hallmark collection accessed via msigdbr (v26.1.0), or ImmuneSigDB C7 collection (*46*, *96*). Additional gene sets were defined from literature sources by taking DEGs with padj <0.05; |log2FC|>1.5 split into UP or DOWN sets (*15*, *18*, *82*, *95*); a BATF2 CUT&RUN dataset selected for genes near peaks bound by BATF2 in wild-type hematopoietic stem cells and not in BATF2 knockout (*81*). Consensus UP and DOWN BACH2 regulons were built by finding genes that appeared in at least two sources of: UP: genes upregulated in BACH2-overexpressing T cells (*85*, *87*), ARCHS4 BACH2 coexpression, or genes repressed in more than one BACH2 knockout study sourced from the Enrichr TF Perturbation database (*111*), DOWN: genes downregulated in BACH2-overexpressing T cells (*85*, *87*), or genes upregulated in more than one BACH2 knockout study sourced from the Enrichr TF Perturbation database. Transcription factor activity was assessed using regulons from the CollecTRI database (*45*), split into activated and repressed gene sets, and fgsea was run using the same parameters. Gene sets were converted to human or murine orthologs using g:orth (*112*) to run against the human CAR or murine P14 T cell datasets, respectively.

#### Functional enrichment analysis of differentially expressed genes

Enrichment analysis was performed using Enrichr (*111*). Statistical significance was calculated using a Fisher exact test, and false discovery rate (FDR) was controlled using the Benjamini–Hochberg procedure. Only terms meeting an adjusted p-value (FDR) threshold of < 0.05 and a combined score > 10, which integrates significance and deviation from expected rank, were retained for downstream analyses. Gene lists from enriched terms were used for downstream network analyses.

#### Network construction and differential correlation analyses

For each condition (BATF2 Day 0, BATF2 Day 9, CAR Day 0, CAR Day 9, n ≥ 3 per group), we constructed gene coexpression networks from the protein-coding gene set using pairwise Spearman rank correlations across biological replicates. Edges were retained when the absolute correlation coefficient exceeded a predefined threshold (|ρ| ≥ 0.5) and the corresponding nominal P value was <0.05, yielding condition-specific networks that capture robust gene–gene associations. Differential correlation (Δρ) networks were then generated by matching edges between condition pairs (for example, BATF2 Day 0 vs BATF2 Day 9, or BATF2 vs CAR at a fixed time point) and computing the difference in correlation (Δρ) for each shared gene pair. We applied Fisher z-transformation–based tests to estimate the statistical significance of Δρ, prioritizing edges with both large magnitude changes and low P values. Final networks and subnetworks, annotated with condition-specific ρ, Δρ, and P values, were exported as edge tables and visualized in Python using standard graph libraries, with node colors denoting curated functional categories.

### Co-immunoprecipitation coupled to mass spectrometry (CoIP/MS)

#### Sample preparation

GD2.CAR T cells were transduced with CAR-P2a-HA:BATF2 or CAR-P2a-V5:AEL-CREM and expanded in a 6-well GRex chamber (Wilson Wolf P/N 80240M) for ten days before harvesting for “day 0” timepoint samples. For the day 9 BATF2 sample, CAR-P2a-HA:BATF2 T cells were co-cultured with CHLA-255.GFP tumor cells in 40 mL cultures in a GRex at E:T of 1:4 at baseline and the same number of added tumor targets added on Day 3 and 6 of coculture for 3 rounds of stimulation total.

CD8+ T cells were enriched by bead-based positive selection (Stem Cell Technologies, cat#17853) and pelleted as ∼100e6 cell aliquots; a small aliquot was reserved for flow cytometric assessment of purity. Flash-frozen pellets were retrieved from −80°C and all subsequent steps were performed on ice or at 4°C. Pellets were resuspended in chilled NTEN lysis buffer supplemented with 1X protease and phosphatase inhibitors (50mM Tris pH 7.3, 170mM NaCl, 1mM EDTA, 0.5% NP-40) and lysed by probe sonication using the Branson sonicator (20% output, 6 cycles, 30s ON/1 min OFF). Lysates were clarified by ultracentrifugation at 100,000 × g for 20 min at 4°C, and the supernatant was immediately transferred to a pre-chilled tube. Protein concentration was estimated spectrophotometrically using Bradford reagent (BioRad; cat# 5000205, absorbance at 595 nm). 1-6 mg total protein from ∼80-400e6 enriched cells were used per IP.

Magnetic beads (30 μL) coupled to anti-V5 (Millipore Sigma, cat# SAE0203), anti-HA (Pierce, cat# 88836), or mouse IgG1 isotype control (MBL International, cat# M075-11) were equilibrated by washing 2X with NTEN buffer prior to use. Clarified lysate was evenly divided between the experimental IP (anti-V5 or anti-HA) and isotype control (IgG1) fractions and incubated on a rotating wheel for 1 hr at 4°C. Following magnetic separation, the unbound flow-through fraction was aspirated. Beads were washed 3X with NTEN buffer. Bead-bound fractions were immediately stored at −80°C pending mass spectrometry analysis.

#### Sample and data processing

The protein complex was eluted from the beads in NuPAGE™ LDS Sample Buffer and resolved on 10% Bis-Tris NuPAGE gel (Life Technologies), visualized with Coomassie Brilliant blue-stain, excised into gel pieces. The whole lane was in-gel digested with trypsin and LysC protease mixture as described previously (*113*). The peptide pools were concentrated in a speed vac and dissolved in 5% methanol containing 0.1% formic acid loading solution. The LC-MS/MS analysis was carried out on a nano-LC 1200 system (Thermo Fisher Scientific, San Jose,CA) coupled to Orbitrap Fusion Lumos (Thermo Fisher Scientific, San Jose, CA) mass spectrometer. The data acquisition was carried out for a 110min gradient time in a data dependent mode. The full MS scan was acquired in Orbitrap in the range of 300 to 1400 m/z at 120,000 resolution followed by MS2 in IonTrap with HCD fragmentation. The MS raw data was searched in Proteome Discoverer software (v2.1, Thermo Scientific, San Jose, CA) with Mascot algorithm (v2.4, Matrix Science) (*114*) against the human NCBI refseq database (updated 2021_12_23). The precursor and fragment mass tolerance were set to 20ppm and 0.5Da respectively. Maximum miscut of 2 with trypsin enzyme, dynamic modification of oxidation on (M), protein N-term acetylation, deamidation (N/Q) and destreak (C) was allowed. The peptides identified from mascot result file were validated with 5% false discover rate (FDR) in Percolator. The protein-level quantification was conducted with label-free iBAQ approach using ‘gpGrouper’ algorithm (*115*). The log-transformed iBAQ values were used for differential analysis. For statistical assessment, missing value imputation was employed through sampling a normal distribution N(μ-1.8 σ, 0.8σ), where μ, σ are the mean and standard deviation of the quantified values. For differential analysis, we used the moderated t-test and log2 fold changes as implemented in the R package.

#### Protein interactome filtering

Following mass spectrometric acquisition, protein abundance was quantified using intensity-based absolute quantification (iBAQ). Putative non-specific and background prey proteins with ≥50% FT (frequency of occurance) in the Jurkat CRAPome 2.0 reference set (*116*) were filtered. Filtered prey lists were retained for downstream high-confidence interactome scoring, independently scored by SAINTexpress (*117*) and SAINTq (*118*). SAINTexpress was applied to spectral counts to calculate AvgSpec and BFDR for each bait-prey pair (AvgSpec= raw/averaged spectral counts/abundance signal across replicates). SAINTq was independently applied to iBAQ-derived to calculate AvgP and BFDR (AvgP= Avg probability of interaction). Both calculate per bait-prey pair across replicates comparing prey abundance in IP samples vs corresponding IgG controls. Expected proportion of false positives estimated by Bayesian False Discovery Rate BFDR <=0.05. High-confidence HA-Batf2 and V5-AelCrem interactors were scored by the following criteria: SAINTq average probability (AvgP) ≥ 0.75, Bayesian false discovery rate (BFDR) < 0.05, log₂ fold-change (IP/IgG) ≥ 1.0, and adjusted p-value ≤ 0.05 relative to the day-matched IgG control. For HA-Batf2, a total of 623 proteins were quantified at Day9 (378 significant) and 147 at Day0 (57 significant; 41 shared across timepoints) and for V5-AelCrem, 176 proteins were recovered.

Pre-ranked Gene Set Enrichment Analysis (*119*) was performed using the MSigDB Hallmark gene set collection (*46*) to identify pathways differentially enriched in the BATF2 interactome between Day 0 and chronic (Day 9) stimulation conditions, or in the AEL-CREM interactome at day 0. TF/cofactor annotations towards the functional sorting of the BATF2 and AELCREM interactomes is based on (*94*) which contains a census of curated ∼1,639-gene human TFs.

### Whole proteome Profiling

A portion of the same CD8 bead enriched cell pellet used for co-IP/MS was lysed in 50mM ammonium bicarbonate buffer by repeated freeze thaw cycle in liquid nitrogen and boiling for 5min at 90 °C. The protein lysate mixture was quantified using the Bradford assay. The protein sample was digested with trypsin and LysC protease mixture overnight at 37 °C. The digested peptide was acidifed in 1% formic and quantified using the Pierce™ Quantitative Colorimetric Peptide Assay (Thermo Scientific 23275). The LC-MS/MS analysis was carried out on a nanoElute2 UHPLC system (Bruker Daltonics, Germany) coupled online to timsTOF Ultra2 mass spectrometer (Bruker Daltonics, Germany) equipped with a CaptiveSpray nano-electrospray ion source. The data acquisition was performed in diaPASEF mode. The peptide samples were directly loaded onto a 25cm x 75µm PepSep Ultra C18 column (Bruker Scientific LLC) and separated using a 30-min total run time at 250nl/min. The diaPASEF raw data was processed using DIA-NN (v 2.2.0) (*120*) in a library-free search mode. The fasta file containing 20,529 human proteins was downloaded from UniprotKB (downloaded 2024_10_22). The precursor ions were generated from FASTA in silico digest with deep learning-based spectra, retention times (RTs), and ion mobilities (IMs) prediction. The proteolytic enzyme Trypsin/P was selected, allowing for up to one missed cleavage. Variable modification of oxidation (M) and protein N-term acetylation was allowed. Parameter specifics include peptide lengths ranging from 7–30, precursor charges from 1–4, precursor m/z from 300–1800, and fragment ion m/z from 200–1800. The MBR was enabled and precursor identification was set at 1% FDR. The DIA-NN precursor-level report file was processed through gpGrouper algorithm (*115*) to obtain protein level quantification using the label-free iBAQ approach. The Ms1.Normalised values were used for iBAQ calculation. The iBAQ expression was median normalized and log-transformed for statistical analysis. For statistical assessment, missing value imputation was employed through sampling a normal distribution N(μ-1.8 σ, 0.8σ), where μ, σ are the mean and standard deviation of the quantified values. For differential analysis, we used the moderated t-test and log2 fold changes as implemented in the R package.

### Integrative analyses of proteomics and transcriptomics datasets

Five transcriptome/proteome-wide analyses were integrated to arrive at the persistence filter, rank active TF programs and nominate driver TFs based on the following criteria: (1) Active TFs in the BATF2-OE CAR-T cell context were filtered by differential expression (ashr-shrunkLFC >0.5, padj<0.05) in BATF2-OE vs CAR for Day 0 (acute) and Day 9 (chronic stimulation, per-gene layer). The pre-filter of differential expression (LFC >0.5, padj<0.05) was applied to exclude developmentally restricted/neuroendocrine TFs that are not T-cell context-specific. (2) Deducing the persistent TF regulon activity from Day 0 to Day 9: TF regulon activity (scorable regulon ≥5 targets per TF) gated by differential expression was obtained by decoupler-ULM ES (Univariate Liner Model, padj <0.05, 0.25, signed net measuring both transcriptional upregulation and repression) to rank active TF programs per day and nominate driver TFs. Regulon activity was quantified with a univariate linear model (ULM). For each regulon, target-gene weights were regressed gainst the gene-wise expression shift (mean_BATF2 – mean_CAR, VST normalized), restricted to that regulon’s targets. The resulting t-statistic of the regression slope is the activity score: positive values indicate net activation in BATF2-OE CAR-T cells, negative values represent net repression. (3) Whole-proteome LFC (BATF2-OE Day 9 normalized to Day 0, p-value <0.05) and Hallmark (*67*) or Gene Ontology (*121*) pathway enrichment; (4) manually curated target effector lists from canonical immunology and cross-referenced against origin-traceable database sets programmatically fetched from Enrichr (MSigDB_Hallmark_2020, Reactome_2022); (5) a BATF2-PPI-independent filter limited to regulon activities of TFs that do not associate with BATF2 protein.

The driver TF was nominated by regulon-activity ranking, and the curated targets of effector TFs were assembled from that TF’s canonical program. Effector membership was manually curated and cross-referenced to origin-traceable database sets MSigDB_Hallmark_2020, Reactome_2022 from Enrichr and the CRE arm target sets were fetched from TRRUSTv2, NAR/ChEA3, and NAR)/ENCODE (*100*, *122*, *123*). Subsequently, a persistence filter in the pipeline nominated driver TFs based on regulon activity ranking, coordinated target effector induction, Day 9-retained regulon and BATF2-PPI independent TFs, as TFs interacting with BATF2 appear largely suppressed.

#### Transcription factor activity inference

For each TF, per-sample VST-normalized expression values were regressed against signed CollecTRI regulon weights (+1 for activating targets, −1 for repressive targets) via ordinary least squares resulting in a weighted signed network (36,740 activating / 5,789 repressing). Only TF regulons that passed the minimum target size threshold of ≥5 target genes were included in the analyses. The t-value of the regression slope was taken as the Univariate Linear Model (ULM) enrichment score (ES). ULM was applied independently to each replicate sample, yielding a per-replicate ES distribution for every TF regulon where its sign indicates the direction of inferred TF activity (positive = activation, negative = repression). For each TF and timepoint, the BATF2-OE vs CAR contrast was summarized as ΔES= mean(ES_BATF2-OE) - mean(ES_CAR). Because ΔES is on a scale dependent on regulon-size hence not comparable across TFs, it was standardized to a bias-corrected standardized mean difference (Hedges’ g =ΔES/SD_pooled * J, J=1-3[4(n1+n2)-9]), appropriate for a small replicate number of n=3. Hedges’ g is a post-hoc effect size metric to compare the standardized enrichment scores (ES) between two distinct experimental or clinical groups. g values were computed on native CollecTRI net with the exception of BACH2 and AHR regulons which were derived from a curated gene list of 40 and 6 targets, respectively. Conventional g thresholds of ∼0.2 (small), ∼0.5 (medium) and ∼0.8 (large) were applied to determine effect size of each TF regulon activity. Statistical significance was assessed using Benjamini-Hochberg-adjusted p-values (ULM BH-padj) which are independently derived from the same per-replicate contrasts and are reported independently of Hedges’ g magnitude. Occasionally, since Hedges’ g can be inflated when across-replicate variance is very small, significance was gated on BH-padj instead of Hedges’ g.

For each BATF2-partner TF, we computed a per-day CollecTRI-ULM activity score (weighted mean of target BATF2-vs-CAR expression difference). The targets driving that score were ranked by signed contribution, and the regulon activity-core is defined as the minimal set of a TF’s concordant CollecTRI regulon genes that account for ≥90% (cumulative contribution) of its net activity signal (ULM-weighted BATF2-vs-CAR effect) on a given day. This is a deterministic cut on the regulon activity shift unlike the GSEA leading edge and does not involve a running enrichment score, ranked whole-genome walk or permutation null. Significant testing is separately done by Hedges’ *g =*bias-corrected, BH_FDR_perday.

#### Prerank normalized enrichment score (NES)

Regulons were restricted to positive (activating) targets only (weight > 0), excluding repression targets as per standard GSEA convention. Running-sum enrichment scores were computed using GSEApy prerank (seed = 42, 1000 permutations, minimum regulon size ≥ 5) and normalized to yield NES, with typical values ranging from approximately −2.5 to +2.5. Significance was determined by permutation-based FDR q-values.

### Mice

All mouse strains are on the C57BL/6J background. All animals used were treated according to the provisions of the Animal Welfare Act, PHS Animal Welfare Policy, NIH Guide for Care and Use of Laboratory Animals, and the policies and procedures of the Center for Comparative Medicine (CCM) at Baylor College of Medicine (BCM), which is fully accredited by the Association of Assessment and Accreditation of Animal Care, International (AAALAC). Studies were approved by the Baylor College of Medicine Institutional Animal Care and Use Committee under protocol number AN-8935.

#### Murine B16-GD2 melanoma model

The lentiviral plasmid encoding murine GD3 synthase (ST8SIA1) and GD2 synthase (B4GALNT1) (pLV-EF1a-GD3synth-P2a-GD2synth-P2a-FFluciferase-WPRE-mPGK-Puro) was designed in Metelitsa Laboratory and manufactured by VectorBuilder (Chicago, IL, Vector ID: VB230221-1434qqj). Lenti-X HEK293T cells (Takara Bio, cat. no. 632180) were maintained in DMEM medium supplemented with 10% fetal bovine serum (FBS), 1% GlutaMAX, 1mM sodium pyruvate, 1% MEM non-essential amino acids, and 1 mM HEPES at 37 °C in 5% CO2 and were passaged at approximately 60% confluence. Lentiviral particles were generated as previously described (*30*) and concentrated using Lenti-X Concentrator (Takara Bio, cat. no. 631232) according to the manufacturer’s instructions and used to transduce B16F10 *(ATCC - CRL-6475).* To isolate GD2+ cells, cells were stained with human anti-GD2 antibodies, and single-cell sorting was performed using an FACS AriaI cell sorter (BD Biosciences) into TC-treated U-bottom 96-well plates. Cultures derived from single cells were screened for GD2 expression, expanded, and cryopreserved in liquid nitrogen. For mouse tumor experiments, B16F10-GD2 cells were trypsinized and washed upon reaching 90% confluency. 3*10⁵ B16F10-GD2 cells were injected subcutaneously into C57BL/6J mice at 6-10 weeks of age.

For murine CAR T-cell generation, CD3⁺ T cells were isolated from the spleens of donor mice using the MojoSort™ Mouse CD3 T Cell Isolation Kit (BioLegend, Cat. #480024) according to the manufacturer instructions and activated for 24 hours in the same way like for LCMV experiments. Activated CD3⁺ T cells were transduced with an empty vector expressing Thy1.1 (EV), the murine GD2.CAR (expressing mGreenLantern), or both GD2.CAR and BATF2 (Thy1.1 transduction marker). Cells were cultured for 5 days post-transduction. When tumor nodules were established (8–10 days after B16F10-GD2 cell injection), mice received 1*10⁶ cells of each indicated T cell population intravenously for tumor-control experiments. A group of mice remained untreated to monitor tumor growth in the absence of CAR T-cell treatment. Tumor length and width were measured for 20 days following CAR T-cell transfer. Tumor volume was calculated using the following formula: 0.5 × L × W². Blood samples were collected on day 8 after CAR T cell transfer and analyzed by spectral flow cytometry (Cytek Aurora 5L; Cytek) to assess CAR T-cell proliferation and persistence.

To compare the immunophenotypes of murine GD2.CAR and GD2.CAR/BATF2 T cells, congenically distinct (*Cd45^.1/.2^* or *Cd45^.1/.1^*) donor CD3⁺ T cells were used. On day 5 of culture, transduced cells were sorted using a BD FACSAria S8 cell sorter (BD Biosciences) and co-transferred into tumor-bearing mice at 5*10⁵ cells of each population under the same conditions described above. Tumors, spleens, tumor-draining lymph nodes, and blood were collected 14–20 days after cell transfer and analyzed by flow cytometry.

#### LCMV experiments

For LCMV infection experiments, mouse CD8^+^ T cells were transduced and transferred into LCMV clone 13 infected mice according to (*124*). Briefly, spleens were collected from congenically distinct P14 (*Cd45^.1/.2^* or *Cd45^.1/.1^*) mice. CD8^+^ T cells were isolated by negative selection using the MojoSort Mouse CD8 T Cell Isolation Kit (Biolegend) according to manufacturer’s instructions. Isolated CD8^+^ cells were seeded in a non-TC-treated 24 well plate at 1-3 million cells/mL and stimulated in T cell stimulation media containing anti-CD3 (1 µg/mL) and anti-CD28 (0.5 µg/mL) antibodies in cRPMI (10% FBS + 1% pen-strep) with 1% MEM Non-Essential Amino Acids (Gibco), 1 mM sodium pyruvate (Gibco), 50 µM beta-mercaptoethanol, 1 ng/mL rhIL-2 and rhIL-7 for 24 hours. Activated cells were washed, resuspended in 30% Percoll (Cytivia), underlaid with 60% Percoll, and spun at 800 *x g* for 20 minutes with the centrifuge brake off. Cells were resuspended in stimulation media, added to a non-TC-treated plate coated with 7-20 µg of RetroNectin (Takara). Titered viral supernatants were added to the culture at an MOI of 1 and incubated at 37°C for 20-30 hours. After transduction, cells were washed, stained with anti-CD90.1, and sort-purified using a BD S8 or FACSAriaII. Transduced cells were pooled at a 1:1 ratio and 50,000-0,000 cells were transferred into CD45^.2/.2^ B6 recipients which were infected with LCMV clone 13 1-2 days prior to transfer. Mice were euthanized at 15 or 30 days post transfer for flow cytometry and single-cell RNA sequencing analysis.

#### LCMV Sample Processing

To collect tissues, mice were euthanized and perfused with PBS via the hepatic vein and the right ventricle prior to collection of tissues. Spleens were mechanically dissociated, treated with RBC Lysis buffer (Tonbo), washed, and filtered to generate a single-cell suspension. Livers were mechanically dissociated, and leukocytes were subjected to density gradient centrifugation using a 44%/67% Percoll gradient (Cytiva).

#### Single cell RNA-seq data analysis

Single-cell RNA sequencing data were generated from P14 T cells transduced with BATF2 or AEL-CREM retroviral constructs and compared with empty vector (Thy1.1+) and untransduced (Thy1.1−) controls, all isolated at day 15 of chronic LCMV infection, alongside naïve P14 T cells from uninfected mice. Libraries were processed using the 10x Genomics platform with hashtag oligonucleotide (HTO) multiplexing and aligned using Cell Ranger Count v8.0.1 (10x Genomics) to a custom GRCm39-based reference genome containing transgene sequences for human BATF2 and AEL-CREM. Downstream analysis was performed in R using Seurat v5 (*125*). Cells were assigned to samples by manual HTO demultiplexing using thresholds of ≥100 total HTO counts and ≥80% dominant hashtag signal. Quality-control filtering retained cells with <7.5% mitochondrial transcripts, >600 detected genes, and >1000 UMIs. SCTransform v2 normalization with regression of nCount_RNA was performed prior to dimensionality reduction and clustering. Principal component analysis, UMAP visualization, and nearest-neighbor clustering were optimized by parameter grid search, with final clustering performed using 20 principal components, 100 neighbors, a minimum distance of 0.3, and clustering resolution of 0.2. All clusters except cluster 1 (cycling) were found to have a mirrored cluster distinguished by UMI counts and nuclear-encoded mitochondrial genes. Attempts to regress on these features did not resolve this mirroring, so mirrored clusters were grouped based on similar marker gene profiles for cluster-based analysis: naïve clusters 2 and 5, Tpex clusters 3 and 7, Tinter clusters 3 and 6. Differential gene expression analysis was performed using Seurat’s FindMarkers function with Wilcoxon rank-sum testing (logfc.threshold = 0) for bulk and cluster-specific comparisons between experimental conditions. Functional enrichment analysis was conducted as described above.

### Statistics

Flow cytometry data were analyzed using Flowjo v10. The UMAP (*104*) and FLOWSOM (*49*) plugins were used for clustering analysis of spectral flow cytometry data. Summary graphs and statistical analysis were performed in GraphPad Prism v10.

Time-course experiment comparisons for one experimental condition versus control were analyzed for significance via mixed-effects models with time and condition as factors. Significance is reported for the main effect of condition, which estimates the overall difference between conditions across all sampled timepoints. One-way ANOVA with Dunnett’s multiple comparisons test was employed for experiments comparing multiple conditions to a control at a single timepoint. Two-way ANOVA with Tukey’s multiple comparisons test was utilized for time-course experiments requiring comparisons between all conditions. Two-way ANOVA with Dunnett’s multiple comparisons test was utilized for time-course experiments requiring comparisons to a single control condition. Statistical analyses are listed in Table S13. Statistics for large datasets are reported in dedicated supplementary tables.

### Protein modeling

*Boltz2 modeling.* Structures of human bZIP pairs and designed bZIP variants paired with human bZIPs were predicted using Boltz-2 (v2.2.1) (*126*). For each protein pair, Boltz-2 was run with three random seeds, and three predicted models were retained for scoring. Predicted complexes were evaluated using the interface prediction metric ipSAE (*127*), which was calculated from the Boltz-2-predicted structures and averaged across models for each protein pair.

To further weight electrostatic interactions, which are important for determining bZIP binding specificity, a custom electrostatic score was calculated. Lys and Arg residues were assigned a charge of +1 and Asp and Glu residues a charge of −1, using only the terminal side-chain atoms (NZ for Lys; NH1/NH2 for Arg; OE1/OE2 for Glu; OD1/OD2 for Asp). Charged residues in opposing chains were considered to be in contact when the minimum distance between their terminal side-chain atoms was within 8 Å. Oppositely charged residue pairs were classified as favorable contacts, whereas contacts between residues of the same charge were classified as unfavorable. Each contact was weighted according to the inverse square of the inter-residue distance (1/*d*^2^), where d is the minimum atom-to-atom distance between the charged side chains, with a lower bound of 2.5 Å to avoid singularities. A raw electrostatic score was calculated as the distance-weighted fraction of favorable minus unfavorable contacts, normalized by the total weight of all contacts:

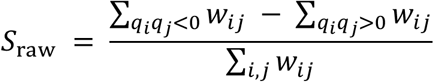

where *w_ij_* is the weight (1/*d*_ij_^2^) of the contact between residues *i* and *j*, and *q_i_* and *q_j_* are their charges.

*S*_raw_ ranges from −1 (all contacts unfavorable) to +1 (all contacts favorable). To give greater weight to interfaces with more contacts, the final electrostatic score was calculated as:

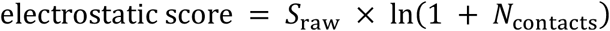

where N_contacts_ is the total count of residue pairs within the 8 Å cutoff.

A composite interaction metric, termed score1, was then calculated to combine predicted interface quality with electrostatic interactions:

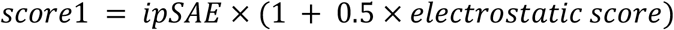

using the mean ipSAE and mean electrostatic score calculated across the three predicted models for each protein pair. Score1 values were used to generate the Boltz2 interaction heatmaps.

#### Potapov and Fong modeling

Predictions from previously published (*34*) and (*58*) bZIP interaction models were generated for the same set of human bZIP pairs and designed bZIP variants analyzed with Boltz2. Because more negative scores indicate stronger predicted binding, the heatmap color scale was inverted to maintain consistent visualization across models.

#### Model validation

To assess accuracy of the models, predictions for human bZIP pairs were compared with experimentally measured binding affinities for 36 human bZIPs reported by Reinke et al. (2013). Pearson’s correlation coefficient (R) was calculated between model scores and experimental (*K_d_*) values measured at two temperatures, 37 and 21 °C. Model scores were correlated against –*K_d_* to align the direction of the experimental binding metric with the model scores, such that stronger binding corresponded to higher values. Classification performance was evaluated using the area under the receiver operating characteristic curve (AUROC). For this analysis, pairs were classified as binders if *K_d_* ≤ 500 nM at two or more temperatures and as non-binders if *Kd* > 5000 nM at all measured temperatures. Pairs that did not satisfy either criterion were excluded from AUROC analysis.

#### Structure visualization

Representative Boltz2-predicted complexes were selected for visualization for AEL-CREM and human bZIP pairs to compare high-scoring (ATF1, CREB1, CREM) and low-scoring (FOS, JUN, MAF) interactors. Structures were visualized in PyMOL, with AEL-CREM and the interacting human bZIP shown as separate chains.

## Acknowledgements

Data analysis was performed on the HPC cluster that is managed by the Biostatistics and Informatics Shared Resource (BISR) and supported by an NIH S10 Shared Instrument Grant S10-OD032185, NCI P30-CA125123 and institutional funds from the Dan L Duncan Comprehensive Cancer Center and Baylor College of Medicine. BCM Mass Spectrometry Proteomics Core (RRID:SCR_027015) is supported by the Dan L. Duncan Comprehensive Cancer Center NIH award (P30 CA125123), CPRIT Core Facility Award (RP210227). This project was supported by the Cytometry and Cell Sorting Core at Baylor College of Medicine with funding from the CPRIT Core Facility Support Award RP240432 and the NIH (CA125123, OD036336, and OD038251), and the assistance of Joel M. Sederstrom and Claude C. Chew. BCM Mass Spectrometry Proteomics Core (RRID:SCR_027015) is supported by the Dan L. Duncan Comprehensive Cancer Center Award (P30 CA125123), CPRIT Core Facility Awards (RP210227), Intellectual Developmental Disabilities Research Center Award (P50 HD103555), and NIH High End Instrument Award (S10 OD026804, Orbitrap Exploris 480). We thank the Texas Children’s Cancer and Hematology Center, Texas Children’s Hospital Flow Cytometry Core Facility for technical assistance and instrument access, specifically, the Sony SH800 sorter. We acknowledge a Texas Childrens’ Hospital Research Institute instrument grant for a Celleca cell counter used in this work. We acknowledge the use of an Echo Acoustic Liquid Handler in the Therapeutic Innovation Center (THINC) at Baylor College of Medicine and the assistance of Dr. Calla Olson. We thank members of Andras Heczey’s lab for assistance with the GPC3.CAR protocol, the GPC3.BBz plasmid, and HUH7.GFP tumor cells. R.S. wishes to thank Spyridoula Vasileiou and Joel Christie for technical discussions. We thank the Rooney lab for sharing an initial protocol for the CHLA-255.GFP repeat tumor stimulation model.

## Author Contributions

Conceptualization: J.M., R. S., W.H., G.F.

Methodology: J.M., R.S., P.G., Y.W., L.M., S.H., M.S., W.H., G.F.

Investigation: J.M., R.S., S.H., M.S., P.G., A.Z., S.Z., A.C., L.L., V.Y.

Visualization: J.M., R.S., S.H., M.S., P.G., G.F., A.Z., S.Z., L.L.

Supervision: V.Y., L.M., W.H., G.F.

Writing—original draft: J.M., R.S., S.H., M.S., and G.F.

Writing—review & editing: J.M., R.S., S.H., M.S., W.H., and G.F.

## Data and materials availability

Datasets and code will be made available upon final journal publication.

## AI Usage

Microsoft CoPilot (Basic), Open AI (GPT 5.4), Claude (Sonnet 4.6 and 5.0, Opus 4.8) were used to edit text for grammar and conciseness, and to draft code. Microsoft CoPilot (Basic), Open AI (GPT 5.4), Claude (Sonnet 4.6 and 5.0, Opus 4.8) were used to generate standalone scripts for RNAseq, AP-MS, proteomic, bZIP library data analyses, coexpression network analyses and visualization. AI-assisted scripts were executed in Google Colaboratory or R studio (v.). All scripts and outputs were reviewed and verified by the authors.

